# ntRoot: Computational Inference of Human Ancestry at Scale from Genomic Data

**DOI:** 10.1101/2024.03.26.586646

**Authors:** René L Warren, Lauren Coombe, Johnathan Wong, Parham Kazemi, Inanc Birol

## Abstract

Ancestry information is essential to large cohort studies, yet it is often unavailable or inconsistently measured. For studies with a genome sequencing component, current ancestry prediction approaches are hindered by high computational demands and complex input requirements. We present ntRoot, a computationally-lightweight method for inferring human super-population-level ancestry from whole genome assemblies or raw short or long sequencing data. Utilizing an alignment-free variant detection framework, ntRoot employs a succinct Bloom filter data structure to efficiently query diverse genomic data inputs. Demonstrated on over 600 human genome sequencing datasets—including complete genomes, draft assemblies, and over 280 independently-generated datasets—ntRoot accurately predicts geographic labels, a descriptor of human populations, and shows high concordance with traditional methods such as ADMIXTURE (*R*^*2*^ = 0.9567) when predicting ancestry fractions. It achieves these predictions within 30 minutes for complete and draft genomes and within 1 hour and 15 minutes for 30X sequencing data, using a maximum of 13GB and 68GB of RAM, respectively. ntRoot offers both global and local ancestry inference, delivering high-resolution predictions across genomic loci. This paradigm fills a critical gap in cohort studies by enabling rapid, resource-efficient, and accurate ancestry inference at scale, advancing the characterization of continental-level ancestry in the genomic era.

**Author Summary:** Study concept: RLW. Software implementation: RLW, LC, JW, PK. Data analysis: RLW, LC. Manuscript development: RLW, LC. Manuscript editing: RLW, LC, JW, PK, IB. Funding acquisition: IB.

## Introduction

Our ancestry information, encoded within our DNA, is crucial for genomic studies. Individuals from different ancestries may have a different genetic makeup and associated disease risk factors, yet association studies do not always take ancestry into consideration or simply leave it as a confounding variable, either because that information is costly to compute, is based on incomplete genotype data, or is not reliably measured. On a population scale, the HapMap Project (1) was the first to profile common patterns of variation between individuals. Subsequently, the 1000 Genomes Project (1kGP) (2) and the Simons Genome Diversity Project (SGDP) (3) catalogued human genomic variation across the whole genomes of healthy individuals from various populations, to generate a baseline for the study of population genetics.

Ancestry admixture inference can be divided into Local Ancestry Inference (LAI) and Global Ancestry Inference (GAI). LAI is the estimation of ancestry along a DNA sequence. This paradigm is based on the fact that during chromosomal crossover, proximal regions of DNA are carried together. Hence, even if an individual’s ancestors were admixed generations ago, DNA fragments may be sourced from single ancestry. While GAI offers a genome-wide summary, LAI provides higher resolution ancestry inferences across genomic loci (4).

Most existing computational approaches developed for ancestry inference are based on single nucleotide variants (SNVs), leveraging the availability of population-scale genotype data from HapMap (1) and 1kGP projects (5). Tools for GAI analysis include the widely-applied model-based STRUCTURE program (6), principal component analysis-based utilities Rye (7) and EIGENSTRAT (8), and ADMIXTURE for likelihood model-based estimation of ancestry in unrelated individuals (9). HAPAA (10) and HAPMIX (11), both designed for LAI, use Hidden Markov Models (HMMs) and explicitly incorporate linkage disequilibrium (LD), the degree of correlation between allelic SNVs. RFMix (12) is an HMM-based admixture model with conditional random forest that leverages phase and LD information (13) for both GAI and LAI, and SNVstory (14) uses machine learning-based models to compute GAI from sequencing data. Additional LAI methods include G-Nomix, FLARE, and SALAI-Net (15–17). While these utilities have their strengths, large-scale studies and routine applications may benefit from more flexible genomic input and smoother execution to better align with current data throughputs and computational demands in the genomic era.

Here, we present ntRoot, a scalable and flexible method for super-population-level (continental, as defined by 1kGP) LAI and GAI analysis from whole genome assemblies or raw genome sequencing datasets. It uses a sequence alignment-free genome variant detection framework (18) that employs a succinct Bloom filter (19) data structure, which contains a reduced sequence representation. This filter is first used to identify SNVs and then leverages integrated variant call sets (IVC) from 1kGP to estimate continental ancestry based on established 1kGP labels (Figs. 1a, and S1). Bloom filters are probabilistic data structures commonly used in bioin-formatics software because they offer a low-memory alternative, enabling scalable applications in the era of large sequencing datasets (20,21). The ntRoot paradigm offers both global and local ancestry inference, providing high-resolution predictions across genomic loci. In our manuscript, the term “ancestry” does not refer to genetic ancestry. Instead, it refers to group labels of geographical locations used to describe a population, as established by the 1kGP and in accordance to The National Academies of Sciences, Engineering, and Medicine guidelines (22).

**Fig. 1.**
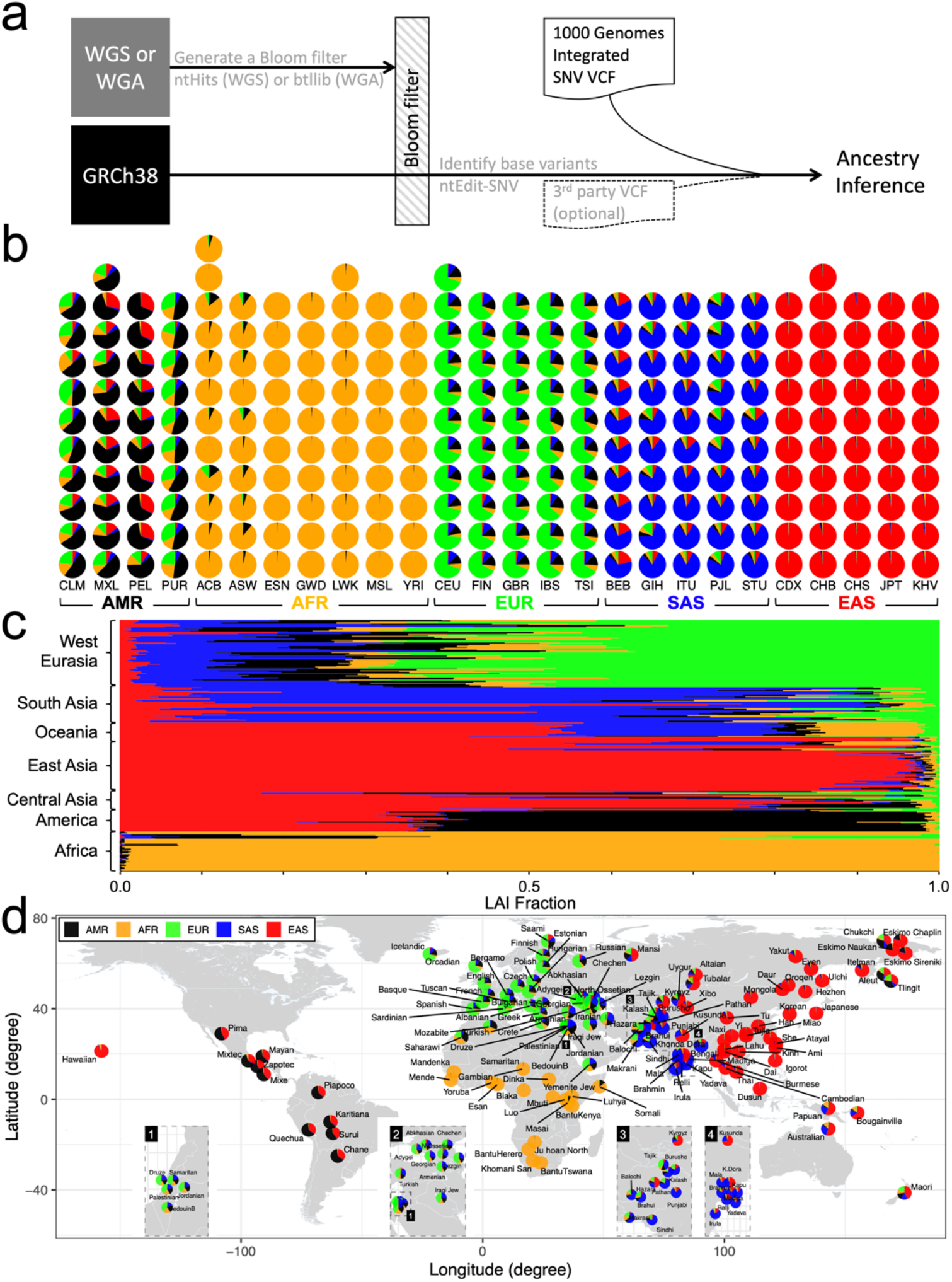
Ancestry inference from genomic data. **a**, Schematic of the ntRoot ancestry inference workflow using whole genome sequencing (WGS) or assembly (WGA) data. Using ntEdit in SNV mode (ntEdit-SNV), each genomic base of the human genome reference GRCh38 queries a Bloom filter built from an individual WGS or WGA data with either ntHits or btllib, respectively; Identified SNVs are cross-referenced with the 1kGP IVC *(23,24)*. Optionally, ntRoot can take a third-party VCF input (dashed lines) to cross-reference the 1kGP integrated variant call set for ancestry predictions (not presented in this study). Human ancestry composition estimates derived from LAI on **b**, the 266 WGS 1kGP validation set grouped by 5 super-populations (1kGP continental ancestry labels AMR:American; AFR:African; EUR:European; SAS:South Asian; EAS:East Asian) and 26 populations (1kGP labels CLM:Colombian; MXL:Mexican-American; PEL:Peruvian; PUR:Puerto Rican; ACB:African-Caribbean; ASW:African-American South West; ESN:Esan; GWD:Gambian; LWK:Luhya; MSL:Mende; YRI:Yoruba; CEU:CEPH (Centre d’Étude du Polymorphisme Humain – Utah); FIN:Finnish; GBR:British; IBS:Spanish; TSI:Tuscan; BEB:Bengali; GIH:Gujarati; ITU:Indian; PJL:Punjabi; STU:Sri Lankan; CDX:Dai Chinese; CHB:Han Chinese; CHS:Southern Han Chinese; JPT:Japanese; KHV:Kinh Vietnamese), and **c**, 279 SGDP WGS discovery set from 129 distinct populations **d**, shown in the global context (using 1kGP continental ancestry labels), with inset regions zoomed-in.

## Methods

We adapted the base-editing utility ntEdit (18) for SNV prediction and ancestry inference (Figs. 1a and S1). This mode leverages k-mer (sequence words of length k)-based analysis to identify alternate base possibilities across genomic datasets, bypassing the error-checking step otherwise used for genome polishing in ntEdit. As with the polishing mode, the SNV calling feature of ntEdit requires a primary Bloom filter (BF) built from k-mers of a user-specified length k, derived from a sequencing data input. After BF construction, each and every k-mer of the provided reference sequence is queried, 5’ to 3’, and the last, 3’-most base is permutated to identify potential alternate bases at that position (Fig. S1). Alternate bases are reported in a variant call format (VCF) file when their observed frequency surpasses a user-defined threshold. The method does not explicitly rely on paired-end read information or alignment-based positional context, focusing on k-mer presence/absence from the BF instead (more details below).

After cross-referencing the putative SNVs identified with the 1kGP IVC (23,24), we average the SNV allele frequency (AF) within each of the five continental super-populations labels, globally for the genome under scrutiny, and locally for genomic windows (details below). We also keep an independent tally of non-zero AF SNVs at both local (per genomic window of a certain size, referred to as *tile* hereafter) and global (whole genome) levels. We rank the super-populations separately for each tile and the genome using those combined metrics, as an indicator of the most likely ancestry. We note that ntRoot can also predict ancestry from a user input VCF file generated from other sources (i.e., not just ntEdit; Fig. 1a), but this feature is not presented in the current study.

### Single nucleotide variant detection with ntEdit v2

The functionality of ntEdit was extended to include single nucleotide variant detection. In this mode, each genomic base of the human reference genome (GRCh38), represented by individual k-mers, is interrogated in turn using the k-mer representation from the individual of interest, stored in a BF. Reference k-mers and their constituent overlapping k-mers (e.g., a 25-mer contains 25 overlapping k-mers) query the BF for presence/absence. Putative SNVs are recorded when alternate 3’-end A, T, C, or G base positions on the original k-mer show sufficient support. Specifically, at any given position in the genome, each k-mer variant and its associated overlapping k-mers are queried, while shifting to the next k-mer by j bases. Once k/j k-mers have been queried, a possible SNV is recorded after assessing the proportion of successful Bloom filter k-mer hits (k_ct_/j) and confirming that it meets or exceeds the user-defined −*Y* parameter (set to 0.55 or 55% by default). Putative SNVs are then cross-referenced with integrated variant callsets from the 1kGP, marking the concordant positions in an output VCF file (Fig. S1).

### Ancestry inference workflow

On whole genome sequencing (WGS) read datasets, the ntRoot workflow (driven by the python script, ntroot --reads) launches ntEdit variant calling, followed by the ancestry inference step. The ntEdit pipeline first executes ntHits (v1.0.2, user-defined parameter k=55, https://github.com/bcgsc/ntHits), which outputs a BF (19) (BF) of robust (non-error) read k-mers. ntEdit (v2.0.0, https://github.com/bcgsc/ntEdit) loads the generated BF and, using the human genome (hg) reference GRCh38, interrogates every k-mer it comprises in turn, querying the BF for presence/absence as per above to generate the variants in VCF format. ntRoot loads a succinct version of the 1kGP integrated variant call sets where reported SNVs have an allele frequency (AF) of 1% or more in at least one human super-population label (EAS, AFR, EUR, SAS or AMR) and the SNVs reported by ntEdit are cross-referenced against the 1kGP call set input VCF. On genome sequence assemblies (WGA) and other 1X representation of an individual’s genome, ntRoot builds a BF using the btllib common code library (25) (ntroot --genome). The remaining steps in WGA follow the same process as WGS (Fig. 1a). Once the ntEdit execution has completed, the ntEdit output VCF file is parsed, summing and averaging the variant AF for each of the five super-populations while tracking the number and rate of non-zero (termed nz in [1] and [2]) AF SNVs over all cross-referenced SNVs. Separate tallies are kept for the whole genome and for each 5 Mbp tile (user-defined parameter --tile, default 5 Mbp). Global (GAI) and local (LAI) ancestry inference scores are calculated as:

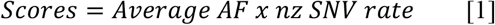

nz: non-zero

SNV: single nucleotide variant AF: SNV allele frequency

Where Average AF=(Sum of AF for each SNV) / (Total number of SNVs) for a given label

for the whole genome and locally, for each tile. For each super-population label, the non-zero SNV rate is defined as:

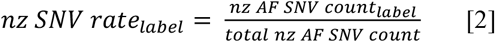

The GAI and LAI scores are bounded within the range [0, 1], with 0 and 1 representing lowest and highest confidence, respectively. The highest-scoring ancestry is reported for the former (GAI), globally, and the associated super-population label assigned to each tile for the latter (LAI), locally. When each tile has been assigned, the LAI super-population fraction for each label is reported as:

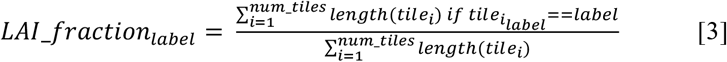

The GAI and LAI scores are output in a tab-separated variable (TSV) output file (_ancestry-predictions_tile<tile size>.tsv). When the --lai option is specified, ntRoot will additionally output the ancestry predictions and scores per tile (_ancestry-predictions-tile-resolution_tile<tile size>.tsv).

### Data

In our study we tested ntRoot on over 600 data sets, which we broadly separated into whole genome sequence (WGS) and whole genome assembly (WGA). The WGS set included 3 different cohorts: **validation, discovery** and **benchmark**. The WGS validation and benchmark sets consisted of 266 and 100 1kGP sequence datasets, respectively and were used to test the accuracy and compute performance of the method, respectively. The WGS discovery dataset (n=279) is independent from 1kGP and was sequenced as part of the Simons Genome Diversity Project (SGDP). The WGA sets comprised complete and assembled genomes in draft stages from both independent (e.g., HuRef, KOREF, CN1) and 1kGP sources (e.g., HG02055 and HG002). Details on the datasets, software commands and run parameters are included in Table S1 and Supplementary Methods.

### Circular genome representations

For all aforementioned experiments and for each human super-population, we tracked tile assignments and SNV density in TSV files for all non-zero AF SNVs reported by ntRoot and cross-referenced with the 1kGP integrated variant call set. That information was then plotted in R using circlize (26). All scripts developed to support our study are available on zenodo (10.5281/zenodo.10869033).

### Benchmarking

ntRoot was mainly (WGA and 1kGP validation set) benchmarked on a server with 144 Intel(R) Xeon(R) Gold 6150 2.70GHz CPUs with 2.1TB RAM, using 48 threads. Additionally, we used a server-class system with 144 Intel(R) Xeon(R) Gold 6254 3.1 GhZ CPUs with 2.9 TB RAM for processing the large SGDP WGS data volumes and benchmarking SNVstory and ADMIXTURE. We extracted the wall clock run time and peak memory from the output of the UNIX time command preceding all ntRoot, SNVstory and ADMIXTURE processes.

## Results and Discussion

To benchmark the performance of ntRoot, we derived super-population-level (label-based continental/geographic) ancestry for a random subset of the 1kGP whole genome sequencing (WGS) dataset (27) organized into the 26 human populations as defined by the 1kGP (WGS validation set, Fig. S2, Table S2, n=266, 10-12 per population) and observed congruent ancestry inferences for all 266 individuals compared to 1kGP assigned labels when derived globally (Fig. S3, Tables S2-S4). The ancestry composition was inferred locally (i.e., LAI) using 5Mbp or 2Mbp tiles (Table S5), and the estimated super-population contributions were reported for the entire genomes of the 266 and 279 WGS 1kGP and SGDP datasets, respectively (Fig. 1b-d, Fig. S4, Table S6). We note that, based on LAI, the ancestry compositions of Americans (Fig. 1b, AMR), Europeans (EUR) and South Asians (SAS) are more diverse than are those of Africans (AFR) and East Asians (EAS), consistent with historical accounts of population movement (28). Inferring super-population ancestries locally (i.e., LAI) is information-rich (Fig. 2) but we also caution about deriving a GAI label from LAI (i.e., largest pie chart slice) because, out of the 266 1kGP WGS validation datasets, one (CLM:Columbian, 0.4%) and three labels (1 CLM, 2 PUR:Puerto Rican, 1.1%) are inconsistent with the highest super-population fraction, when using 5Mbp vs. 2Mbp tiles, respectively (Table S5). Interestingly, all four incongruent samples represent individuals from the Americas, where population admixture has occurred (28) and hence, genetic ancestry is likely less homogeneous. This is the case for HG01243, a Puerto Rican individual with a recently documented (29) mixed, but primarily African (AFR) ancestry, originally assigned an American (AMR) label by the 1kGP consortium. ADMIXTURE and ntRoot GAI and ancestry fractions for this individual are consistent and align with the revised and recently published AFR ancestry assignment (29) (Fig. S5, Tables S7-S8).

**Fig. 2.**
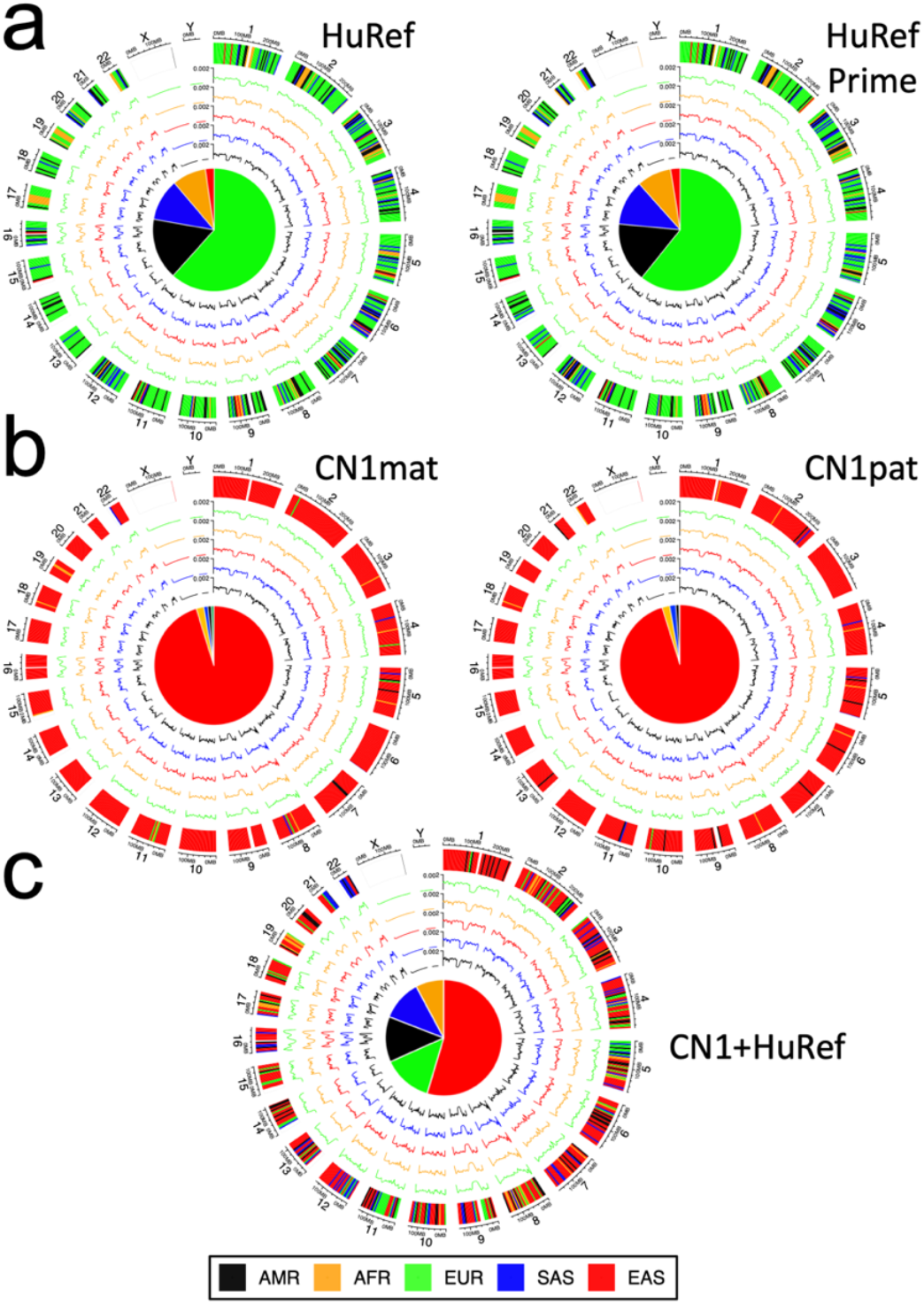
Circular genome representations of ancestry inference. using WGA for **a**, European HuRef (Levy *et al*., 2007) and HuRefPrime haplotypes, **b**, Han Chinese CN1 (Yang *et al*., 2023) (mat:maternal, pat:paternal) haplotypes and **c**, simulated HuRef/CN1 diploid rearrangement. The outer track shows ntRoot local ancestry assignments (LAI) along each chromosome using 5Mbp tiles (AMR:American; AFR:African; EUR:European; SAS:South Asian; EAS:East Asian), with each subsequent inner track showing the corresponding SNV density for each super-population. Inner pie charts report the predicted global ancestry composition (GAI).

ntRoot LAI using the chromosome-scale diploid HuRef (30) (Fig. 2a), CN1 (31) (Fig. 2b) and haploid KOREF (32) (Fig. S6) genome assemblies of three individuals revealed prevalent ancestry based on their highest composition (EUR:61%, EAS:95%, EAS:95%, respectively), consistent with both the highest GAI scores, and the published European (30), Chinese (31) and Korean (32) ancestry for each (Tables S7-S8). Furthermore, ntRoot correctly inferred EAS and EUR as the two most prevalent super-population ancestries from synthetic diploid genome admixtures of HuRef and CN1 (Fig. 2c, Table S8).

Using orthogonal datasets distinct from the 1kGP data, WGS-based SGDP results (WGS discovery set, Table S6) and WGA-based GAI/LAI results (Tables S7 and S8, respectively) independently validate our approach. The SGDP discovery data set uses slightly different geographic names compared to 1kGP, but despite this and the fact that we are using 1kGP integrated variant call VCFs to analyze the continental-level composition of SGDP, the location and ancestry fraction is congruent (Fig. 1d). Further, the full SGDP set comprises multiple individuals from the same populations, and the ntRoot predictions are consistent between these biological replicates (Table S6).

Unlike existing tools, ntRoot can infer ancestries directly from a genome assembly, even when it is in a draft stage (Fig. S7). Processing haploid WGA with ntRoot averaged 29m and consumed at most 13 GB of RAM, while it ran in under 1h36m requiring at most 68 GB of RAM, on average, on 279 SGDP WGS datasets (22–87-fold genome coverage, average 44-fold). Despite varying WGS sequencing coverage, ntRoot produced robust predictions, using as little as 22X coverage (Table S6). A more controlled sequencing coverage titration experiment using Illumina WGS data from the HG002 (also known as GM24385 and huAA53E0) cell line derived from an Ashkenazi Jewish individual of primarily European ancestry reveals that ntRoot’s predictions remain robust even at 12.5-fold coverage. The LAI EUR fraction, which represents the GAI and the largest ancestry fraction, shows a delta of 2.62% between full coverage (44.83%) and 12.5-fold coverage (42.21%), demonstrating the stability of ntRoot’s ancestry predictions at reduced sequencing depths (Table S10). Further, the predictions appear to be sequencing data type-agnostic, as long-read WGS data from Illumina, PacBio (HiFi), and Oxford Nanopore Technologies (ONT, V14 kit) instruments yield consistent ntRoot predictions of ancestry fractions for HG002. The largest percentage variation observed is 0.7%, seen in both the LAI AFR fraction (Illumina vs. ONT) and the LAI EUR fraction (Illumina vs. ONT). This consistency is also observed in KOREF, where both Illumina and PacBio WGS data yield nearly identical ancestry fraction predictions. For example, the LAI EAS fraction, which represents the GAI and the largest ancestry fraction for KOREF, is 97.72% for Illumina and 97.90% for PacBio, with the LAI AFR, AMR, and EUR fractions showing 100% consistency across both platforms (Tables S1, S7-S8). ntRoot ancestry predictions are influenced by the parameter k, with the GAI being accurately inferred at higher k values (e.g., k45-k70). However, as k decreases, the fraction compositions can fluctuate. This is expected, as smaller k values generally increase the recall of ancestry-discriminant SNVs, but at the cost of precision. Using the Genome in a Bottle SNV benchmarks for HG002 (33), we found k=55 to be a Pareto-optimal choice, representing a good balance between sensitivity (0.8051) and precision (0.8262) for identifying ancestry-discriminant SNVs (Figs. S8 and Table S11). Based on these findings, we recommend running ntRoot with the default or higher k values on all datasets.

Compared to SNVstory (14), one of the more recent GAI-based utility that was shown to have more consistent performance on continental and subcontinental ancestry inference tasks compared to RFMix and ADMIXTURE, ntRoot has the same or similar accuracy as SNVstory and ADMIXTURE (100% and 99% concordance with 1kGP labels, respectively) while running ~4X and ~250X faster (1h09m vs. 4h25m vs. 273h42m, for ntRoot, SNVstory and ADMIXTURE) and with 1.3-6.7X less compute memory when inferring the ancestry of 100 1kGP individuals from WGS data (WGS benchmark set, Tables S9, S12-S13). We note that while SNVstory outputs probabilities for GAI predictions—often 0.99 or 1 on the WGS benchmark set—these values do not provide insights into ancestry composition. In contrast, ntRoot and ADMIXTURE offer more detailed ancestry fractions, which are useful for characterizing populations and individuals with significant ancestry admixtures (Fig. S9, Table S12). Moreover, ntRoot demonstrates high concordance with ADMIXTURE overall (Fig. S10a, *R*^2^ = 0.9567) and across each ancestry (Fig. S10b), with *R*^2^ values ranging from 0.9237 (AMR) to 0.9974 (AFR). Additionally, a mean squared error (MSE) analysis further supports this concordance, with an overall MSE of 0.78%, and individual MSE values for each ancestry as follows: EAS – 0.44%, SAS – 0.70%, AMR – 1.19%, AFR – 0.06%, and EUR – 1.48%. We note, however, that ntRoot’s LAI-based fraction is different from the fine-scale genetic structure analysis previously reported (34) and, in its current implementation, does not have the resolution to characterize genetic ancestry at that level. Because ntRoot cross-references the SNVs identified with an integrated variant call set collated from 2,709 1kGP individuals (23,24), exact ancestry compositions are not expected. This is especially true given that the continental (and population) labels have been assigned by 1kGP based on the geographical location where the original 1kGP samples were taken/where the individuals lived at the time of collection, and that the information that ntRoot uses to cross-reference its SNVs is built on a relatively limited number (n=2,709) of those individuals. Given a human population size of eight billion in mid-November 2022 (35) and that large human migrations have taken place in the past 10,000 years, 2,709 individuals are not expected to represent the full breadth of nucleotide variations in humans. We also stress that ntRoot is designed to predict the likely continental (super-population)-level ancestry based on pre-assigned labels, which may or may not be representative of true genetic ancestry. To mitigate these limitations, potential strategies include providing ntRoot with an IVC built from other or additional sources (e.g., the Genome Aggregation Database - gnomAD or The International Genome Sample Resource - IGSR), ensuring it is compiled from more comprehensive and diverse reference panels (36,37).

The ntRoot framework automates each step via a driver script, markedly simplifying the execution of ancestry prediction pipelines compared to the often convoluted workflows of other modern predictors (Table S13). It builds on top of the sequence alignment-free ntEdit paradigm (18), for identifying and cross-referencing sequence variants in whole genome sequencing and whole genome assembly datasets with an efficient use of computational resources. It accomplishes the feat by using Bloom filters generated from either ntHits (38) (in WGS mode) or btllib (25) (in WGA mode). Bloom filters, widely used in bioinformatics applications and the genomics realm (21,39), are popular, succinct probabilistic data structures that accommodate large datasets with a light compute memory footprint, as demonstrated herein. In previous work (18), we showed how Bloom filter false positive rates do not affect downstream ntEdit analyses. This is because ntEdit requires k-mer redundancy at each base for its nucleotide base identification. Further, ntRoot cross-references the variants it identifies (or inputs) with integrated variant call sets, which only looks at a subset of human variations characterized and curated by third parties (23,24). As observed with ntRoot predictions from WGA, even a 1X genome representation is sufficient to accurately predict continental-level ancestry. We also find that, in a typical ntRoot run that uses either WGA or WGS as input, roughly 2-3M SNVs are cross-referenced with the 1kGP integrated variant call sets and used to derive adequate ancestry inferences (Table S7; SNV count column).

## Conclusion

With its streamlined, memory-efficient, fast, and easy-to-execute workflow, ntRoot generates GAI and LAI-based admixture profiles that provide detailed insights into population genetics. We anticipate that ntRoot will broadly facilitate geographic-level ancestry inference, offering reliable and objective demographic information from the sequencing data of modern technologies. By enabling rapid and accurate pedigree inference at scale, ntRoot addresses a critical gap in cohort studies and holds promise for advancing super-population-level ancestry predictions for association studies in the genomic era.

## Data and Code Availability

The data used in this study is summarized in Supplementary Material, Table S1. ntRoot is freely available on GitHub (https://github.com/bcgsc/ntroot).

## Supporting information captions

Supplementary Methods

Supplementary Table S1. **Data Accessions**

Supplementary Table S2. **Continental, super-population level summary of ntRoot GAI results on representative 1kGP whole-genome sequencing datasets** (n=266, WGS validation set) arranged by population, compared to their Assigned Label Counts

Supplementary Table S3. **Continental, super-population level summary of ntRoot GAI results on representative 1kGP whole-genome sequencing datasets (n=266, WGS validation dataset)**, compared to their Assigned Label Counts

Supplementary Table S4. **ntRoot super-population level GAI summary on the 1kGP WGS validation dataset**

Supplementary Table S5. **ntRoot super-population level LAI-based ancestry fraction estimates on the 1kGP WGS validation** set using 5Mbp (5Mbt) or 2Mbp (2Mbt) tiles

Supplementary Table S6. **ntRoot super-population level LAI-based ancestry fraction estimates on the SGDP WGS discovery set**

Supplementary Table S7. **ntRoot super-population level GAI summary on complete and draft genome sequences**, with matching raw whole-genome read sequencing data

Supplementary Table S8. **ntRoot (unless specified) super-population level LAI-based ancestry fraction estimates on complete and draft genome sequences**, with matching raw whole-genome read sequencing data

Supplementary Table S9. **Resource usage** for ntRoot, SNVstory and ADMIXTURE

Supplementary Table S10. **Effect of HG002 (NA24385, Illumina) WGS coverage** on ntRoot ancestry predictions

Supplementary Table S11. **Rationale for selecting the parameter k in ntRoot** (k sweeps)

Supplementary Table S12. **ADMIXTURE (AdM), SNVstory (SnvS) and ntRoot (ntR) ancestry prediction results** on the balanced 1kGP benchmark set of 100 samples (20 per super-population)

Supplementary Table S13. **Resource usage breakdown** by steps for ADMIXTURE on the balanced set of 100, 1kGP samples

Supplementary Fig. S1. **Overview of the SNV detection algorithm** in ntRoot.

Supplementary Fig. S2. **Geolocation of the 26 human populations**, as defined by the 1000 Genomes Project

Supplementary Fig. S3. **ntRoot super-population level GAI compared to 1kGP labels for the 1kGP WGS validation set**

Supplementary Fig. S4. **ntRoot ancestry composition estimates derived from LAI on SGDP WGS discovery data set** from 129 distinct populations shown in the global context

Supplementary Fig. S5. **ntRoot LAI predictions on HG01243 Illumina whole genome sequencing data from 1kGP for a Puerto Rican individual** of mixed, primarily African ancestry, labeling the likely human super-population matching each 5Mbp tile alongside the genome

Supplementary Fig. S6. **ntRoot LAI predictions on KOREF WGA and WGS**

Supplementary Fig. S7. **ntRoot LAI predictions on HG02055 WGA**

Supplementary Fig. S8. **Rationale for selecting the parameter k in ntRoot** (k sweeps)

Supplementary Fig. S9. a, **ADMIXTURE and b, ntRoot ancestry fraction predictions** (y-axis) on the balanced set of 100, 1kGP samples

Supplementary Fig. S10. **Correlation between ntRoot and ADMIXTURE ancestry fractions** on the balanced set of 100, 1kGP samples

## References

1. Gibbs RA, Belmont JW, Hardenbol P, Willis TD, Yu F, Yang H, et al. The International HapMap Project. Nature. 2003 Dec 1;426(6968):789–96.

2. The 1000 Genomes Project Consortium, Corresponding authors, Auton A, Abecasis GR, Steering committee, Altshuler DM, et al. A global reference for human genetic variation. Nature. 2015 Oct 1;526(7571):68–74.

3. Mallick S, Li H, Lipson M, Mathieson I, Gymrek M, Racimo F, et al. The Simons Genome Diversity Project: 300 genomes from 142 diverse populations. Nature. 2016 Oct 13;538(7624):201–6.

4. Tang H, Peng J, Wang P, Risch NJ. Estimation of individual admixture: analytical and study design considerations. Genet Epidemiol Off Publ Int Genet Epidemiol Soc. 2005;28(4):289–301.

5. McVean GA, Altshuler (Co-Chair) DM, Durbin (Co-Chair) RM, Abecasis GR, Bentley DR, Chakravarti A, et al. An integrated map of genetic variation from 1,092 human genomes. Nature. 2012 Nov 1;491(7422):56–65.

6. Pritchard JK, Stephens M, Donnelly P. Inference of population structure using multilocus genotype data. Genetics. 2000;155(2):945–59.

7. Conley AB, Rishishwar L, Ahmad M, Sharma S, Norris ET, Jordan IK, et al. Rye: genetic ancestry inference at biobank scale. Nucleic Acids Res. 2023 May 8;51(8):e44.

8. Price AL, Patterson NJ, Plenge RM, Weinblatt ME, Shadick NA, Reich D. Principal components analysis corrects for stratification in genome-wide association studies. Nat Genet. 2006;38(8):904–9.

9. Alexander DH, Novembre J, Lange K. Fast model-based estimation of ancestry in unrelated individuals. Genome Res. 2009;19(9):1655–64.

10. Sundquist A, Fratkin E, Do CB, Batzoglou S. Effect of genetic divergence in identifying ancestral origin using HAPAA. Genome Res. 2008;18(4):676–82.

11. Price AL, Tandon A, Patterson N, Barnes KC, Rafaels N, Ruczinski I, et al. Sensitive detection of chromosomal segments of distinct ancestry in admixed populations. PLoS Genet. 2009;5(6):e1000519.

12. Maples BK, Gravel S, Kenny EE, Bustamante CD. RFMix: a discriminative modeling approach for rapid and robust local-ancestry inference. Am J Hum Genet. 2013;93(2):278–88.

13. Uren C, Hoal EG, Möller M. Putting RFMix and ADMIXTURE to the test in a complex admixed population. BMC Genet. 2020;21(1):1–8.

14. Bollas AE, Rajkovic A, Ceyhan D, Gaither JB, Mardis ER, White P. SNVstory: inferring genetic ancestry from genome sequencing data. BMC Bioinformatics. 2024 Feb 20;25(1):76.

15. Hilmarsson H, Kumar AS, Rastogi R, Bustamante CD, Montserrat DM, Ioannidis AG. High Resolution Ancestry Deconvolution for Next Generation Genomic Data [Internet]. 2021 [cited 2025 Mar 18]. Available from: http://biorxiv.org/lookup/doi/10.1101/2021.09.19.460980

16. Oriol Sabat B, Mas Montserrat D, Giro-i-Nieto X, Ioannidis AG. SALAI-Net: species-agnostic local ancestry inference network. Bioinformatics. 2022;38(Supplement_2):ii27–33.

17. Browning SR, Waples RK, Browning BL. Fast, accurate local ancestry inference with FLARE. Am J Hum Genet. 2023 Feb;110(2):326–35.

18. Warren RL, Coombe L, Mohamadi H, Zhang J, Jaquish B, Isabel N, et al. ntEdit: scalable genome sequence polishing. Bioinforma Oxf Engl. 2019 Nov 1;35(21):4430–2.

19. Bloom BH. Space/time trade-offs in hash coding with allowable errors. Commun ACM. 1970;13(7):422–6.

20. Warren RL, Yang C, Vandervalk BP, Behsaz B, Lagman A, Jones SJM, et al. LINKS: Scalable, alignment-free scaffolding of draft genomes with long reads. Gigascience. 2015 Dec 1;4(1):s13742–015-0076–3.

21. Jackman SD, Vandervalk BP, Mohamadi H, Chu J, Yeo S, Hammond SA, et al. ABySS 2.0: resource-efficient assembly of large genomes using a Bloom filter. Genome Res. 2017;27(5):768–77.

22. Committee on the Use of Race, Ethnicity, and Ancestry as Population Descriptors in Genomics Research, Board on Health Sciences Policy, Committee on Population, Health and Medicine Division, Division of Behavioral and Social Sciences and Education, National Academies of Sciences, Engineering, and Medicine. Using Population Descriptors in Genetics and Genomics Research: A New Framework for an Evolving Field [Internet]. Washington, D.C.: National Academies Press; 2023 [cited 2024 Aug 27]. Available from: https://www.nap.edu/catalog/26902

23. Zheng-Bradley X, Streeter I, Fairley S, Richardson D, Clarke L, Flicek P, et al. Alignment of 1000 Genomes Project reads to reference assembly GRCh38. GigaScience [Internet]. 2017 Jul 1 [cited 2024 Aug 26];6(7). Available from: https://academic.oup.com/gigascience/article/doi/10.1093/gigascience/gix038/3836916

24. Lowy-Gallego E, Fairley S, Zheng-Bradley X, Ruffier M, Clarke L, Flicek P, et al. Variant calling on the GRCh38 assembly with the data from phase three of the 1000 Genomes Project. Wellcome Open Res. 2019 Dec 30;4:50.

25. Nikolić V, Kazemi P, Coombe L, Wong J, Afshinfard A, Chu J, et al. btllib: A C++ library with Python interface forefficient genomic sequence processing. J Open Source Softw. 2022 Nov 4;7(79):4720.

26. Gu Z, Gu L, Eils R, Schlesner M, Brors B. circlize Implements and enhances circular visualization in R. Bioinforma Oxf Engl. 2014 Oct;30(19):2811–2.

27. Byrska-Bishop M, Evani US, Zhao X, Basile AO, Abel HJ, Regier AA, et al. High-coverage whole-genome sequencing of the expanded 1000 Genomes Project cohort including 602 trios. Cell. 2022 Sep;185(18):3426–3440.e19.

28. Willerslev E, Meltzer DJ. Peopling of the Americas as inferred from ancient genomics. Nature. 2021 Jun 17;594(7863):356–64.

29. Zimin AV, Shumate A, Shinder I, Heinz J, Puiu D, Pertea M, et al. A reference-quality, fully annotated genome from a Puerto Rican individual. Novembre J, editor. Genetics. 2022 Feb 4;220(2):iyab227.

30. Levy S, Sutton G, Ng PC, Feuk L, Halpern AL, Walenz BP, et al. The Diploid Genome Sequence of an Individual Human. Rubin EM, editor. PLoS Biol. 2007 Sep 4;5(10):e254.

31. Yang C, Zhou Y, Song Y, Wu D, Zeng Y, Nie L, et al. The complete and fully-phased diploid genome of a male Han Chinese. Cell Res. 2023 Jul 14;33(10):745–61.

32. Kim HS, Jeon S, Kim Y, Kim C, Bhak J, Bhak J. KOREF_S1: phased, parental trio-binned Korean reference genome using long reads and Hi-C sequencing methods. GigaScience. 2022 Mar 24;11:giac022.

33. Zook JM, McDaniel J, Olson ND, Wagner J, Parikh H, Heaton H, et al. An open resource for accurately benchmarking small variant and reference calls. Nat Biotechnol. 2019 May;37(5):561–6.

34. Wellcome Trust Case Control Consortium 2, International Multiple Sclerosis Genetics Consortium, Leslie S, Winney B, Hellenthal G, Davison D, et al. The fine-scale genetic structure of the British population. Nature. 2015 Mar;519(7543):309–14.

35. “World Population Prospects 2022, Graphs / Profiles”. United Nations Department of Economic and Social Affairs, Population Division. 2022. [Internet]. Available from: https://population.un.org/wpp/Graphs/Probabilistic/POP/TOT/900

36. Fairley S, Lowy-Gallego E, Perry E, Flicek P. The International Genome Sample Resource (IGSR) collection of open human genomic variation resources. Nucleic Acids Res. 2020 Jan 8;48(D1):D941–7.

37. Chen S, Francioli LC, Goodrich JK, Collins RL, Kanai M, Wang Q, et al. A genomic mutational constraint map using variation in 76,156 human genomes. Nature. 2024 Jan 4;625(7993):92–100.

38. Mohamadi H, Chu J, Coombe L, Warren R, Birol I. ntHits: de novo repeat identification of genomics data using a streaming approach [Internet]. Genomics; 2020 Nov [cited 2024 Mar 6]. Available from: http://biorxiv.org/lookup/doi/10.1101/2020.11.02.365809

39. Wong J, Coombe L, Nikolić V, Zhang E, Nip KM, Sidhu P, et al. Linear time complexity de novo long read genome assembly with GoldRush. Nat Commun. 2023 May 22;14(1):2906.

