## Supplementary Material for "ntRoot: Computational Inference of Human Ancestry at Scale from Genomic Data"

### Supplementary Methods

***Ancestry inference of 266 1kGP individuals, using whole genome sequencing datasets (WGS validation set)***

The 30X WGS data was downloaded from https://www.internationalgenome.org/data-portal/data-collection/30x-grch38 (Table S1). We randomly selected 266 individuals, with 10-12 representatives per each 26 human populations (defined in [1,2]) and downloaded the associated WGS data with enaBrowserTools (https://github.com/enasequence/enaBrowserTools). We executed ntRoot while tracking the wall-clock run time and memory. An example ntRoot command on a 1kGP WGS dataset:

*ntroot -k 55 --draft GRCh38.fa.gz --reads ERR3242308_ -t 48 -Y 0.55 -l 1000GP_integrated_snv_v2a_27022019.GRCh38.phased_gt1.vcf.gz*

where GRCh38.fa.gz contains the GRCh38 human genome reference sequence with chromosome numbers as FASTA header, ERR3242308_ is the fastq file prefix for the sequencing reads and 1000GP_integrated_snv_v2a_27022019.GRCh38.phased_gt1.vcf.gz the integrated variant call set from 1kGP. We also downloaded Illumina-only (HG01243) and Illumina NovaSeq 6000 and ONT kit v14 Dorado base-called reads for NA24385 (HG002). See Table S1 for the file source and Tables S2-S5 for associated results.

***Ancestry inference on 279 WGS datasets (1kGP-independent, WGS discovery set) from the Simons Genome Diversity Project***

We downloaded all available 279 genomes sequencing datasets (WGS) from the diverse populations of the Simons Genome Diversity project[10] using enaBrowserTools and executed ntRoot (v1.0.0; ntroot --reads -t 48) on each read set separately while tracking the wall-clock run time and memory. We tabulated the fractions of ntRoot ancestry inference from every tsv output (in _ancestry-predictions_tile<tile size>.tsv) within the cohort and detailed in Table S6. Subsequently, we represented these fractions for a randomly-selected representative individual, for each of the distinct 129 populations represented in the main manuscript, Figure 1d. This large data served as a large-scale and independent validation of our approach.

***Ancestry inference on reference, 1kGP-independent WGS datasets***

In separate experiments, we downloaded the whole-genome sequencing datasets (Table S1) of an individual of European ancestry, HuRef (Illumina HiSeq X Five) and of Korean ancestry, KOREF (Illumina HiSeq X Ten / PacBio_SMRT Sequel II) and executed ntRoot (v1.0.0; ntroot --reads -t 48) on those datasets while tracking the wall-clock run time and memory. This experiment served as an independent validation of our 1kGP-based approach, since HuRef and KOREF are not part of the 1kGP cohort. See Tables S7-S8 for results.

***Ancestry inferences using WGA, complete and draft whole genome sequences***

In separate experiments, we downloaded the complete diploid genome assemblies (Table S1) of an individuals of European ancestry (Super-population EUR: European), HG002 (NA24385 complete genome, maternal/paternal haplotypes) and HuRef/HuRefPrime (complete genome, principal/alternate haplotypes), of Han-Chinese (EAS: East Asian) ancestry, CN1maternal/paternal (complete genome, maternal/paternal haplotypes), Korean (EAS: East Asian) ancestry, KOREF (complete genome, pseudohaploid), Puerto Rican (HG01243, AMR: Admixed American) ancestry and African (AFR: African) ancestry, HG02055 (complete genome, maternal and paternal haplotypes) and executed ntRoot (v1.0.0; ntroot --genome -t 48) while tracking the wall-clock run time and memory. We also ran ntRoot on a synthetic HuRef-CN1 admix diploid WGA and three pseudohaploid genome assemblies[11] of HG02055 to test ntRoot on admixed and pseudohaploid assemblies vs. separated, maternal and paternal haplotypes. For the synthetic HuRef-CN1 admix diploid experiment, using a custom script (available on zenodo), we first randomly selected complete paternal or maternal chromosome sequences from HuRef and HuRefPrime (random_chromosome_mix.py GCA_000002125.1_HuRef_genomic.chr-renamed.fa GCA_000212995.1_HuRefPrime_genomic.chr-renamed.fa HuRef-hap.fa), and then separately from CN1mat and CN1pat (random_chromosome_mix.py GWHCBHM00000000.genome.fasta GWHCBHQ00000000.genome.fasta CH1hap). We then executed ntRoot (v1.0.0; ntroot --genome -t 48) on the combined HuRef-hap.fa and CN1-hap.fa FASTA files. See Tables S7-S8 for results.

***HG002 WGS coverage titration, k sweeps and sequencing data type experiments***

For the coverage titration experiment, we subsampled the NA24385 (HG002) Illumina WGS datasets from 2.5 to 55-fold coverage (seqtk sample -s100 7500000 to 165000000000) and ran ntRoot (v1.0.0; ntroot --reads -t 48 -k 55) on each individual read partitions. We also investigated the effect of the parameter k on ancestry predictions using the full-coverage (~60-fold) Illumina WGS data, running ntRoot (v1.0.0; ntroot --reads -t 48) sweeping on k from 30 to 70 in separate runs, with a step value of 5. The sensitivity, precision and F1 metrics were calculated at each k by first making a tally of all ancestry-discriminant SNVs from the ntRoot VCF output, and counting the number of true positives, false positives, and true negatives by comparing ntRoot's SNV predictions against a benchmark from the Genome in a Bottle Consortium[12] (available from: https://www.nist.gov/programs-projects/genome-bottle and https://ftp-trace.ncbi.nlm.nih.gov/ReferenceSamples/giab/release/AshkenazimTrio/HG002_NA24385_son/NISTv4.2.1/GRCh38/HG002_GRCh38_1_22_v4.2.1_benchmark.vcf.gz). Finally, in separate experiments, we ran ntRoot (v1.0.0; ntroot --reads -t 48 -k 55) on recent PacBio Revio (Sequel II) and Oxford Nanopore (kit V14) HG002 long read datasets. See Tables S7-S8 and S10-S11 for results.

***Ancestry inference of 100 1kGP individuals, using whole genome sequencing datasets (WGS benchmarking set)***

The 30X WGS data was downloaded from https://www.internationalgenome.org/data-portal/data-collection/30x-grch38. We randomly selected 100 individuals in total, 20 for each of the five super-populations[1,2], ensuring no overlap with the 1kGP WGS validation set, and downloaded the associated WGS data with enaBrowserTools (https://github.com/enasequence/enaBrowserTools). We executed ntRoot (v1.0.0; ntroot --reads -t 48) on each while tracking the wall-clock run time and memory. To perform ancestry inference using SNVstory[3], we followed the approach described in the SNVstory manuscript. For each of the 100 1kGP individuals, we aligned the corresponding short reads to the human reference GRCh38 using bwa mem[4] (v 0.7.17-r1188; -t 48). Next, we generated a VCF file from the short-read alignments using the GATK[5,6] suite of tools. The necessary files for the GRCh38 reference genome were generated using samtools[7] faidx (v1.17) and gatk CreateSequenceDictionary (v4.5.0.0). Next, the variants were called using gatk HaplotypeCaller (v4.5.0.0), with separate runs launched for each chromosome (option -L), and at most 12 parallel jobs running at a given time. For each individual, all chromosome-specific VCF files generated using gatk HaplotypeCaller were merged using bcftools[7] concat (v 1.17; --threads 48). Finally, the ancestry for the individual was predicted using SNVstory (v3.0.2; --genome-ver 38 --mode WGS), providing the generated VCF file as input. The GAI inference for SNVstory was taken as the 1kGP continental prediction with the highest probability. We also inferred the ancestry of the same 100 1kGP individuals using ADMIXTURE[8]. We used the reference genotype VCF files available from the 1kGP project (https://ftp.1000genomes.ebi.ac.uk/vol1/ftp/data_collections/1000G_2504_high_coverage/working/20201028_3202_raw_GT_with_annot/), excluding the 100 1kGP query individuals. The VCF files generated for SNVstory were merged with this combined reference VCF using bcftools merge (v1.17; --threads 48). This merged VCF file was then filtered to only retain autosomes and biallelic sites using bcftools view (--threads 48 --max-alleles 2). Next, plink2[9] (v2.00a5.12LM; --threads 48 --make-bed --geno 0.1) was used to generate the required input files for ADMIXTURE from the combined VCF file. Finally, the ancestral fractions in the 100 benchmark samples were estimated using ADMIXTURE (v1.3.0; -j48 --supervised), supplying the bed file generated by plink2, and indicating the query samples using a .pop file. See Tables S9, S12-S13 for results.

### Supplementary Tables

**Table S1.** Data Accessions.

| Individual | Type | Samples | Accession | Resource URL |
|---|---|---|---|---|
| **GRCh38** | WGA | NA | GCA_000001405.15 and 10.5281/zenodo.10869033 | https://www.ncbi.nlm.nih.gov/datasets/genome/GCF_000001405.15/ |
| **HuRef[13]** | WGS | 1 | SRR8595488 | https://www.ebi.ac.uk/ena/browser/view/SRR8595488 |
| **HuRef[13]‡** | WGA | 1 | GCA_000002125.1 | https://ftp.ncbi.nlm.nih.gov/genomes/all/GCA/000/002/125/GCA_000002125.1_HuRef/GCA_000002125.1_HuRef_genomic.fna.gz |
| **HuRefPrime[13]‡** | WGA | 1 | GCA_000212995.1 | https://ftp.ncbi.nlm.nih.gov/genomes/all/GCA/000/212/995/GCA_000212995.1_HuRefPrime/GCA_000212995.1_HuRefPrime_genomic.fna.gz |
| **CN1[14]** | WGA maternal | 1 | GWHCBHM00000000 | https://download.cncb.ac.cn/gwh/Animals/Homo_sapiens_CN1_v0.8_mat_GWHCBHM00000000/GWHCBHM00000000.genome.fasta.gz |
| **CN1[14]** | WGA paternal | 1 | GWHCBHQ00000000 | https://download.cncb.ac.cn/gwh/Animals/Homo_sapiens_CN1_v0.8_pat_GWHCBHQ00000000/GWHCBHQ00000000.genome.fasta.gz |
| **HuRef/CN1 simulated diploid** | WGA | 1 | 10.5281/zenodo.10869033 | https://doi.org/10.5281/zenodo.10976332 |
| **KOREF[15]*** | WGA | 1 | GCA_020497085.1 | https://www.ebi.ac.uk/ena/browser/api/fasta/GCA_020497085.1 |
| **KOREF[15]** | WGS (Illumina) | 1 | SRR10861698 | https://www.ebi.ac.uk/ena/browser/view/SRR10861698 |
| **KOREF[15]** | WGS (PacBio Sequel II) | 1 | SRR14759118-21, 23-24 | https://www.ebi.ac.uk/ena/browser/view/SRR14759118 to https://www.ebi.ac.uk/ena/browser/view/SRR14759118 |
| **HG01243** | WGS | 1 | ERR3988858 | https://www.ebi.ac.uk/ena/browser/view/ERR3988858 |
| **NA24385 (HG002)** | WGA maternal | 1 | GCA_018852615.3 | https://www.ebi.ac.uk/ena/browser/view/GCA_018852615.3 |
| **NA24385 (HG002)** | WGA paternal | 1 | GCA_018852605.3 | https://www.ebi.ac.uk/ena/browser/view/GCA_018852605.3 |
| **NA24385 (HG002)** | WGS (llumina NovaSeq 2x250bp) | 1 | SRR11321732 | https://www.ebi.ac.uk/ena/browser/view/SRR11321732 |
| **NA24385 (HG002)** | WGS (ONT V14 Dorado) | 1 | NA | https://labs.epi2me.io/askenazi-kit14-2022-12/ and s3://ont-open-data/giab_lsk114_2022.12/ |
| **NA24385 (HG002)** | WGS (PacBio Revio HiFi Sequel II) | 1 | SRR26402938 | https://www.ebi.ac.uk/ena/browser/view/SRR26402938 |
| **HG02055[1,2]** | WGS | 1 | ERR3988979 | https://www.ebi.ac.uk/ena/browser/view/ERR3988979 |
| **HG02055[11]*** | WGA (GoldRush*,^) | 1 | 10.5281/zenodo.7884681 | https://doi.org/10.5281/zenodo.7884681 |
| **HG02055[11]*** | WGA (Shasta*,^) | 1 | 10.5281/zenodo.7884681 | https://doi.org/10.5281/zenodo.7884681 |
| **HG02055[11]*** | WGA (Flye*,^) | 1 | 10.5281/zenodo.7884681 | https://doi.org/10.5281/zenodo.7884681 |
| **HG02055[16]** | WGA maternal | 1 | JAHEPJ01 | https://www.ncbi.nlm.nih.gov/Traces/wgs/JAHEPJ01 |
| **HG02055[16]** | WGA paternal | 1 | JAHEPK01 | https://www.ncbi.nlm.nih.gov/Traces/wgs/JAHEPK01 |
| **1kGP[1,2]** | WGS | 266 and 100 | Table S4 and S9 | https://www.internationalgenome.org/data-portal/data-collection/30x-grch38 |

| | | | | |
|---|---|---|---|---|
| **SGDP[10#]** | WGS | 279 | PRJEB9586 | https://www.simonsfoundation.org/simons-genome-diversity-project/ https://ddbj.nig.ac.jp/resource/bioproject/PRJEB9586 https://www.ebi.ac.uk/ena/browser/view/PRJEB9586 |
| **1kGP integrated variant call set** | VCF | 2709 | 10.5281/zenodo.10869033 | https://www.internationalgenome.org/data-portal/data-collection/30x-grch38 |

***Pseudohaploid assembly. ^Assembled with WGS Oxford Nanopore datasets[11] (Wong et al., 2022, Table 6). The mat/pat suffix indicates maternal and paternal haplotypes, respectively. ‡HuRef and HuRefPrime haploid WGAs each refer to the principal and alternate haplotypes respectively. #The primary SGDP dataset consists of data from 260 genomes from 127 populations: 39 Africans, 23 Native Americans, 27 Central Asians or Siberians, 49 East Asians, 27 Oceanians, 38 South Asians and 71 West Eurasians according to https://www.simonsfoundation.org/simons-genome-diversity-project/; We identified data (and report results in Table S5) for 279 distinct individuals organized into 129 unique populations (Figure 1d).**

**Table S2.** Continental, super-population level summary of ntRoot GAI results on representative 1kGP whole-genome sequencing datasets (n=266, WGS validation set) arranged by population, compared to their Assigned Label Counts.

| Super-population Code | Population Code | Population Name | Assigned Label Counts | ntRoot GAI Correct Prediction Counts |
|---|---|---|---|---|
| **AMR** | MXL | Mexican-American | 11 | 11 |
| **AMR** | PUR | Puerto Rican | 10 | 10 |
| **AMR** | PEL | Peruvian | 10 | 10 |
| **AMR** | CLM | Colombian | 10 | 10 |
| **AFR** | MSL | Mende | 10 | 10 |
| **AFR** | LWK | Luhya | 11 | 11 |
| **AFR** | ASW | African-American South West | 10 | 10 |
| **AFR** | YRI | Yoruba | 10 | 10 |
| **AFR** | GWD | Gambian | 10 | 10 |
| **AFR** | ESN | Esan | 10 | 10 |
| **AFR** | ACB | African-Caribbean | 12 | 12 |
| **SAS** | STU | Sri Lankan | 10 | 10 |
| **SAS** | BEB | Bengali | 10 | 10 |
| **SAS** | ITU | Indian | 10 | 10 |
| **SAS** | PJL | Punjabi | 10 | 10 |
| **SAS** | GIH | Gujarati | 10 | 10 |
| **EUR** | GBR | British | 10 | 10 |
| **EUR** | IBS | Spanish | 10 | 10 |
| **EUR** | TSI | Tuscan | 10 | 10 |
| **EUR** | FIN | Finnish | 10 | 10 |
| **EUR** | CEU | CEPH* | 11 | 11 |
| **EAS** | KHV | Kinh Vietnamese | 10 | 10 |
| **EAS** | CHS | Southern Han Chinese | 10 | 10 |
| **EAS** | JPT | Japanese | 10 | 10 |
| **EAS** | CDX | Dai Chinese | 10 | 10 |
| **EAS** | CHB | Han Chinese | 11 | 11 |

* Centre d'Étude du Polymorphisme Humain – Utah.

**Table S3.** Continental, super-population level summary of ntRoot GAI results on representative 1kGP whole-genome sequencing datasets (n=266, WGS validation dataset), compared to their Assigned Label Counts.

| Super-Population Code | Super-Population Name | Assigned Label Count | GAI Correct Prediction Count |
|---|---|---|---|
| **AFR** | African | 73 | 73 |
| **AMR** | American | 41 | 41 |
| **EAS** | East Asian | 51 | 51 |
| **EUR** | European | 51 | 51 |
| **SAS** | South Asian | 50 | 50 |
| **TOTAL** | - | 266 | 266 (100.0%) |

**Table S4.** ntRoot super-population level GAI summary on the 1kGP WGS validation dataset. The super-populations are EAS: East Asian, SAS: South Asian, AMR: Admixed American, AFR: African, EUR: European.

| Accession | Alternate ID | Population Label | Super-Population Label | GAI | SNV Count | Non-zero AF SNV Count | GAI score |
|---|---|---|---|---|---|---|---|
| **ERR3242189** | HG01882 | ACB | AFR | AFR | 3,186,519 | 3,167,755 | 0.4414 |
| **ERR3242193** | HG02014 | ACB | AFR | AFR | 3,137,786 | 3,111,573 | 0.4421 |
| **ERR3242218** | HG01914 | ACB | AFR | AFR | 3,172,303 | 3,143,270 | 0.4403 |
| **ERR3242297** | HG01988 | ACB | AFR | AFR | 2,988,030 | 2,930,917 | 0.4388 |
| **ERR3242299** | HG02051 | ACB | AFR | AFR | 3,206,909 | 3,189,368 | 0.4415 |
| **ERR3242306** | HG02143 | ACB | AFR | AFR | 3,168,705 | 3,136,300 | 0.4389 |
| **ERR3242308** | HG02307 | ACB | AFR | AFR | 3,170,854 | 3,150,587 | 0.4423 |
| **ERR3242318** | HG02330 | ACB | AFR | AFR | 3,127,777 | 3,102,525 | 0.4428 |
| **ERR3242321** | HG02419 | ACB | AFR | AFR | 3,124,724 | 3,103,323 | 0.4442 |
| **ERR3242322** | HG02420 | ACB | AFR | AFR | 3,016,590 | 2,963,682 | 0.4395 |
| **ERR3242326** | HG02442 | ACB | AFR | AFR | 3,160,348 | 3,142,986 | 0.4434 |
| **ERR3242740** | HG02541 | ACB | AFR | AFR | 3,128,220 | 3,094,466 | 0.4407 |
| **ERR3239918** | NA19711 | ASW | AFR | AFR | 3,149,808 | 3,124,561 | 0.4416 |
| **ERR3239961** | NA19834 | ASW | AFR | AFR | 3,137,894 | 3,100,899 | 0.4392 |
| **ERR3239963** | NA19900 | ASW | AFR | AFR | 3,129,227 | 3,098,140 | 0.4408 |
| **ERR3239965** | NA19904 | ASW | AFR | AFR | 3,159,895 | 3,129,145 | 0.4394 |
| **ERR3239973** | NA19982 | ASW | AFR | AFR | 3,133,911 | 3,107,279 | 0.4415 |
| **ERR3239974** | NA20126 | ASW | AFR | AFR | 3,060,072 | 3,018,048 | 0.4404 |
| **ERR3239982** | NA20291 | ASW | AFR | AFR | 3,141,807 | 3,124,530 | 0.4442 |
| **ERR3239990** | NA20340 | ASW | AFR | AFR | 3,146,409 | 3,117,144 | 0.4410 |
| **ERR3239993** | NA20348 | ASW | AFR | AFR | 3,158,165 | 3,127,319 | 0.4400 |
| **ERR3240095** | NA19922 | ASW | AFR | AFR | 3,081,920 | 3,040,542 | 0.4396 |
| **ERR3242481** | HG02923 | ESN | AFR | AFR | 3,158,237 | 3,143,080 | 0.4445 |
| **ERR3242487** | HG03100 | ESN | AFR | AFR | 3,197,023 | 3,181,912 | 0.4428 |
| **ERR3242587** | HG03268 | ESN | AFR | AFR | 3,167,022 | 3,151,949 | 0.4441 |
| **ERR3242589** | HG03271 | ESN | AFR | AFR | 3,185,636 | 3,170,682 | 0.4432 |
| **ERR3242612** | HG03160 | ESN | AFR | AFR | 3,211,873 | 3,196,857 | 0.4417 |
| **ERR3242623** | HG03343 | ESN | AFR | AFR | 3,231,466 | 3,216,418 | 0.4417 |
| **ERR3242982** | HG03127 | ESN | AFR | AFR | 3,202,430 | 3,187,118 | 0.4427 |
| **ERR3242986** | HG03175 | ESN | AFR | AFR | 3,212,686 | 3,197,683 | 0.4416 |
| **ERR3242991** | HG03265 | ESN | AFR | AFR | 3,132,528 | 3,117,465 | 0.4460 |
| **ERR3242993** | HG03298 | ESN | AFR | AFR | 3,211,047 | 3,195,779 | 0.4419 |
| **ERR3242357** | HG02461 | GWD | AFR | AFR | 3,172,040 | 3,155,906 | 0.4448 |
| **ERR3242365** | HG02573 | GWD | AFR | AFR | 3,192,014 | 3,174,718 | 0.4425 |
| **ERR3242373** | HG02594 | GWD | AFR | AFR | 3,156,410 | 3,140,070 | 0.4449 |
| **ERR3242379** | HG02623 | GWD | AFR | AFR | 3,155,106 | 3,139,185 | 0.4447 |
| **ERR3242385** | HG02645 | GWD | AFR | AFR | 3,169,900 | 3,153,671 | 0.4438 |
| **ERR3242391** | HG02678 | GWD | AFR | AFR | 3,118,185 | 3,101,143 | 0.4460 |
| **ERR3242395** | HG02721 | GWD | AFR | AFR | 3,159,119 | 3,142,210 | 0.4441 |
| **ERR3242423** | HG02768 | GWD | AFR | AFR | 3,196,630 | 3,180,432 | 0.4427 |
| **ERR3242429** | HG02804 | GWD | AFR | AFR | 3,206,479 | 3,190,618 | 0.4419 |
| **ERR3242457** | HG02816 | GWD | AFR | AFR | 3,190,090 | 3,173,346 | 0.4438 |
| **ERR3239689** | NA19026 | LWK | AFR | AFR | 3,110,637 | 3,092,710 | 0.4452 |
| **ERR3239698** | NA19041 | LWK | AFR | AFR | 3,174,494 | 3,156,605 | 0.4435 |
| **ERR3239706** | NA19309 | LWK | AFR | AFR | 3,202,156 | 3,184,246 | 0.4421 |
| **ERR3239708** | NA19312 | LWK | AFR | AFR | 3,163,420 | 3,145,640 | 0.4431 |
| **ERR3239714** | NA19319 | LWK | AFR | AFR | 3,192,714 | 3,174,618 | 0.4424 |
| **ERR3239726** | NA19347 | LWK | AFR | AFR | 3,163,453 | 3,145,926 | 0.4435 |
| **ERR3239727** | NA19350 | LWK | AFR | AFR | 3,130,396 | 3,112,541 | 0.4443 |
| **ERR3239738** | NA19380 | LWK | AFR | AFR | 3,123,986 | 3,105,442 | 0.4452 |
| **ERR3239744** | NA19393 | LWK | AFR | AFR | 3,185,803 | 3,167,871 | 0.4418 |
| **ERR3239752** | NA19428 | LWK | AFR | AFR | 3,155,303 | 3,137,504 | 0.4435 |
| **ERR3239768** | NA19451 | LWK | AFR | AFR | 3,177,078 | 3,159,053 | 0.4425 |
| **ERR3242471** | HG03084 | MSL | AFR | AFR | 3,199,546 | 3,183,933 | 0.4404 |
| **ERR3242476** | HG03451 | MSL | AFR | AFR | 3,190,982 | 3,175,496 | 0.4420 |
| **ERR3242551** | HG03439 | MSL | AFR | AFR | 3,219,858 | 3,204,118 | 0.4396 |
| **ERR3242559** | HG03066 | MSL | AFR | AFR | 3,225,280 | 3,209,755 | 0.4396 |
| **ERR3242568** | HG03484 | MSL | AFR | AFR | 3,243,027 | 3,227,586 | 0.4393 |

| | | | | | | | |
|---|---|---|---|---|---|---|---|
| **ERR3242570** | HG03547 | MSL | AFR | AFR | 3,205,453 | 3,189,830 | 0.4408 |
| **ERR3242578** | HG03565 | MSL | AFR | AFR | 3,228,611 | 3,212,922 | 0.4395 |
| **ERR3242952** | HG03060 | MSL | AFR | AFR | 3,212,658 | 3,197,009 | 0.4403 |
| **ERR3242966** | HG03382 | MSL | AFR | AFR | 3,166,825 | 3,150,852 | 0.4417 |
| **ERR3242968** | HG03432 | MSL | AFR | AFR | 3,227,029 | 3,211,516 | 0.4399 |
| **ERR3239341** | NA18504 | YRI | AFR | AFR | 3,220,233 | 3,205,057 | 0.4418 |
| **ERR3239349** | NA18519 | YRI | AFR | AFR | 3,210,002 | 3,194,891 | 0.4428 |
| **ERR3239387** | NA18871 | YRI | AFR | AFR | 3,239,814 | 3,224,800 | 0.4413 |
| **ERR3239429** | NA19119 | YRI | AFR | AFR | 3,222,033 | 3,207,023 | 0.4423 |
| **ERR3239551** | NA18910 | YRI | AFR | AFR | 3,202,713 | 3,187,530 | 0.4429 |
| **ERR3239556** | NA18934 | YRI | AFR | AFR | 3,226,763 | 3,211,770 | 0.4419 |
| **ERR3239619** | NA19096 | YRI | AFR | AFR | 3,203,540 | 3,188,547 | 0.4422 |
| **ERR3239622** | NA19117 | YRI | AFR | AFR | 3,154,229 | 3,139,147 | 0.4441 |
| **ERR3239626** | NA19146 | YRI | AFR | AFR | 3,187,343 | 3,172,239 | 0.4430 |
| **ERR3239629** | NA19184 | YRI | AFR | AFR | 3,217,574 | 3,202,556 | 0.4426 |
| **ERR3241834** | HG01136 | CLM | AMR | AMR | 2,752,323 | 2,701,315 | 0.4759 |
| **ERR3241862** | HG01250 | CLM | AMR | AMR | 2,749,878 | 2,700,209 | 0.4783 |
| **ERR3241875** | HG01344 | CLM | AMR | AMR | 2,681,692 | 2,641,536 | 0.4895 |
| **ERR3241878** | HG01356 | CLM | AMR | AMR | 2,716,041 | 2,672,917 | 0.4816 |
| **ERR3241888** | HG01353 | CLM | AMR | AMR | 2,723,381 | 2,683,396 | 0.4834 |
| **ERR3241894** | HG01374 | CLM | AMR | AMR | 2,682,698 | 2,646,860 | 0.4892 |
| **ERR3241898** | HG01437 | CLM | AMR | AMR | 2,662,534 | 2,625,832 | 0.4863 |
| **ERR3241904** | HG01461 | CLM | AMR | AMR | 2,781,384 | 2,709,874 | 0.4648 |
| **ERR3242787** | HG01130 | CLM | AMR | AMR | 2,707,307 | 2,661,499 | 0.4807 |
| **ERR3242791** | HG01280 | CLM | AMR | AMR | 2,689,021 | 2,652,649 | 0.4884 |
| **ERR3239897** | NA19652 | MXL | AMR | AMR | 2,684,554 | 2,641,504 | 0.4852 |
| **ERR3239902** | NA19661 | MXL | AMR | AMR | 2,677,412 | 2,635,126 | 0.4885 |
| **ERR3239904** | NA19664 | MXL | AMR | AMR | 2,666,127 | 2,622,038 | 0.4884 |
| **ERR3239911** | NA19682 | MXL | AMR | AMR | 2,653,571 | 2,616,722 | 0.4934 |
| **ERR3239928** | NA19726 | MXL | AMR | AMR | 2,658,275 | 2,623,563 | 0.4974 |
| **ERR3239940** | NA19759 | MXL | AMR | AMR | 2,618,380 | 2,582,644 | 0.4998 |
| **ERR3239946** | NA19774 | MXL | AMR | AMR | 2,678,609 | 2,634,280 | 0.4873 |
| **ERR3239950** | NA19780 | MXL | AMR | AMR | 2,661,645 | 2,621,548 | 0.4883 |
| **ERR3239956** | NA19789 | MXL | AMR | AMR | 2,699,620 | 2,651,139 | 0.4829 |
| **ERR3240091** | NA19741 | MXL | AMR | AMR | 2,598,423 | 2,566,358 | 0.5047 |
| **ERR3240094** | NA19792 | MXL | AMR | AMR | 2,706,649 | 2,661,706 | 0.4837 |
| **ERR3241988** | HG01577 | PEL | AMR | AMR | 2,684,977 | 2,639,000 | 0.4869 |
| **ERR3241996** | HG01923 | PEL | AMR | AMR | 2,546,247 | 2,517,362 | 0.5129 |
| **ERR3241998** | HG01926 | PEL | AMR | AMR | 2,540,590 | 2,512,437 | 0.5154 |
| **ERR3242004** | HG01938 | PEL | AMR | AMR | 2,575,381 | 2,547,140 | 0.5125 |
| **ERR3242010** | HG01947 | PEL | AMR | AMR | 2,633,025 | 2,594,955 | 0.4970 |
| **ERR3242018** | HG01970 | PEL | AMR | AMR | 2,533,693 | 2,502,472 | 0.5081 |
| **ERR3242024** | HG01979 | PEL | AMR | AMR | 2,628,838 | 2,596,064 | 0.5021 |
| **ERR3242028** | HG02008 | PEL | AMR | AMR | 2,640,034 | 2,608,356 | 0.5048 |
| **ERR3242241** | HG02271 | PEL | AMR | AMR | 2,569,248 | 2,537,914 | 0.5113 |
| **ERR3242811** | HG01961 | PEL | AMR | AMR | 2,533,822 | 2,506,141 | 0.5160 |
| **ERR3241726** | HG00637 | PUR | AMR | AMR | 2,759,816 | 2,699,019 | 0.4655 |
| **ERR3241728** | HG00640 | PUR | AMR | AMR | 2,668,638 | 2,624,708 | 0.4794 |
| **ERR3241763** | HG01054 | PUR | AMR | AMR | 2,664,162 | 2,623,429 | 0.4811 |
| **ERR3241811** | HG01072 | PUR | AMR | AMR | 2,718,191 | 2,671,825 | 0.4755 |
| **ERR3241816** | HG01085 | PUR | AMR | AMR | 2,722,139 | 2,665,212 | 0.4698 |
| **ERR3241822** | HG01104 | PUR | AMR | AMR | 2,694,699 | 2,649,246 | 0.4775 |
| **ERR3241844** | HG01173 | PUR | AMR | AMR | 2,676,271 | 2,631,773 | 0.4780 |
| **ERR3241854** | HG01197 | PUR | AMR | AMR | 2,685,646 | 2,643,155 | 0.4803 |
| **ERR3242774** | HG01161 | PUR | AMR | AMR | 2,776,409 | 2,708,994 | 0.4611 |
| **ERR3242776** | HG01164 | PUR | AMR | AMR | 2,700,734 | 2,652,064 | 0.4753 |
| **ERR3242164** | HG01816 | CDX | EAS | EAS | 2,575,551 | 2,532,627 | 0.5351 |
| **ERR3242214** | HG02373 | CDX | EAS | EAS | 2,668,456 | 2,625,211 | 0.5298 |
| **ERR3242216** | HG02396 | CDX | EAS | EAS | 2,631,855 | 2,588,417 | 0.5328 |
| **ERR3242217** | HG02397 | CDX | EAS | EAS | 2,613,135 | 2,569,460 | 0.5329 |
| **ERR3242262** | HG02353 | CDX | EAS | EAS | 2,627,644 | 2,583,837 | 0.5315 |
| **ERR3242265** | HG02360 | CDX | EAS | EAS | 2,608,205 | 2,564,862 | 0.5335 |
| **ERR3242275** | HG02384 | CDX | EAS | EAS | 2,604,758 | 2,560,807 | 0.5337 |
| **ERR3242279** | HG02390 | CDX | EAS | EAS | 2,590,254 | 2,546,518 | 0.5342 |
| **ERR3242285** | HG02401 | CDX | EAS | EAS | 2,623,556 | 2,579,880 | 0.5316 |
| **ERR3242289** | HG02409 | CDX | EAS | EAS | 2,637,355 | 2,593,908 | 0.5327 |

| | | | | | | | |
|---|---|---|---|---|---|---|---|
| **ERR3239362** | NA18558 | CHB | EAS | EAS | 2,604,389 | 2,559,175 | 0.5323 |
| **ERR3239363** | NA18561 | CHB | EAS | EAS | 2,612,695 | 2,565,500 | 0.5314 |
| **ERR3239370** | NA18572 | CHB | EAS | EAS | 2,679,439 | 2,635,041 | 0.5302 |
| **ERR3239378** | NA18605 | CHB | EAS | EAS | 2,642,200 | 2,596,043 | 0.5311 |
| **ERR3239379** | NA18608 | CHB | EAS | EAS | 2,621,978 | 2,575,231 | 0.5311 |
| **ERR3239496** | NA18544 | CHB | EAS | EAS | 2,631,366 | 2,585,755 | 0.5312 |
| **ERR3239501** | NA18559 | CHB | EAS | EAS | 2,659,175 | 2,614,869 | 0.5315 |
| **ERR3239510** | NA18612 | CHB | EAS | EAS | 2,676,866 | 2,632,454 | 0.5287 |
| **ERR3239511** | NA18613 | CHB | EAS | EAS | 2,632,870 | 2,587,786 | 0.5316 |
| **ERR3239534** | NA18639 | CHB | EAS | EAS | 2,608,848 | 2,566,038 | 0.5344 |
| **ERR3239660** | NA18629 | CHB | EAS | EAS | 2,662,410 | 2,619,065 | 0.5311 |
| **ERR3240183** | HG00442 | CHS | EAS | EAS | 2,643,529 | 2,601,142 | 0.5326 |
| **ERR3241665** | HG00403 | CHS | EAS | EAS | 2,637,645 | 2,595,071 | 0.5316 |
| **ERR3241673** | HG00436 | CHS | EAS | EAS | 2,618,115 | 2,575,429 | 0.5339 |
| **ERR3241675** | HG00457 | CHS | EAS | EAS | 2,651,857 | 2,609,443 | 0.5321 |
| **ERR3241681** | HG00478 | CHS | EAS | EAS | 2,617,084 | 2,573,850 | 0.5347 |
| **ERR3241685** | HG00524 | CHS | EAS | EAS | 2,637,335 | 2,594,462 | 0.5321 |
| **ERR3241697** | HG00556 | CHS | EAS | EAS | 2,643,691 | 2,600,925 | 0.5328 |
| **ERR3241707** | HG00589 | CHS | EAS | EAS | 2,633,437 | 2,590,648 | 0.5336 |
| **ERR3241715** | HG00610 | CHS | EAS | EAS | 2,646,040 | 2,602,961 | 0.5317 |
| **ERR3241723** | HG00628 | CHS | EAS | EAS | 2,655,588 | 2,612,379 | 0.5309 |
| **ERR3239401** | NA18952 | JPT | EAS | EAS | 2,639,995 | 2,597,141 | 0.5314 |
| **ERR3239564** | NA18966 | JPT | EAS | EAS | 2,662,444 | 2,619,348 | 0.5287 |
| **ERR3239574** | NA18995 | JPT | EAS | EAS | 2,644,569 | 2,601,671 | 0.5312 |
| **ERR3239578** | NA19000 | JPT | EAS | EAS | 2,658,942 | 2,616,126 | 0.5306 |
| **ERR3239583** | NA19009 | JPT | EAS | EAS | 2,663,461 | 2,620,483 | 0.5298 |
| **ERR3239590** | NA19058 | JPT | EAS | EAS | 2,624,383 | 2,581,149 | 0.5315 |
| **ERR3239673** | NA18982 | JPT | EAS | EAS | 2,609,073 | 2,566,064 | 0.5318 |
| **ERR3239675** | NA18984 | JPT | EAS | EAS | 2,613,772 | 2,571,434 | 0.5319 |
| **ERR3239678** | NA18988 | JPT | EAS | EAS | 2,628,364 | 2,585,633 | 0.5314 |
| **ERR3239681** | NA19006 | JPT | EAS | EAS | 2,623,314 | 2,580,367 | 0.5321 |
| **ERR3242036** | HG02017 | KHV | EAS | EAS | 2,660,111 | 2,611,411 | 0.5281 |
| **ERR3242038** | HG02020 | KHV | EAS | EAS | 2,619,942 | 2,573,659 | 0.5306 |
| **ERR3242043** | HG02029 | KHV | EAS | EAS | 2,640,424 | 2,597,479 | 0.5320 |
| **ERR3242047** | HG02070 | KHV | EAS | EAS | 2,617,710 | 2,572,571 | 0.5324 |
| **ERR3242049** | HG02073 | KHV | EAS | EAS | 2,631,017 | 2,586,134 | 0.5322 |
| **ERR3242069** | HG01842 | KHV | EAS | EAS | 2,645,543 | 2,599,017 | 0.5301 |
| **ERR3242079** | HG01852 | KHV | EAS | EAS | 2,660,322 | 2,613,546 | 0.5296 |
| **ERR3242092** | HG01867 | KHV | EAS | EAS | 2,647,088 | 2,599,620 | 0.5293 |
| **ERR3242210** | HG02122 | KHV | EAS | EAS | 2,646,683 | 2,601,392 | 0.5297 |
| **ERR3242343** | HG02047 | KHV | EAS | EAS | 2,651,613 | 2,605,275 | 0.5298 |
| **ERR3239281** | NA07051 | CEU | EUR | EUR | 2,610,640 | 2,570,725 | 0.4967 |
| **ERR3239285** | NA10851 | CEU | EUR | EUR | 2,629,778 | 2,590,367 | 0.4958 |
| **ERR3239286** | NA11829 | CEU | EUR | EUR | 2,656,801 | 2,616,970 | 0.4951 |
| **ERR3239288** | NA11831 | CEU | EUR | EUR | 2,636,146 | 2,595,221 | 0.4950 |
| **ERR3239294** | NA11919 | CEU | EUR | EUR | 2,625,154 | 2,585,104 | 0.4961 |
| **ERR3239302** | NA12005 | CEU | EUR | EUR | 2,587,088 | 2,547,189 | 0.4985 |
| **ERR3239309** | NA12155 | CEU | EUR | EUR | 2,629,238 | 2,588,866 | 0.4961 |
| **ERR3239334** | NA12878 | CEU | EUR | EUR | 2,630,986 | 2,590,319 | 0.4951 |
| **ERR3239462** | NA11893 | CEU | EUR | EUR | 2,628,750 | 2,589,168 | 0.4973 |
| **ERR3239473** | NA12342 | CEU | EUR | EUR | 2,644,497 | 2,604,098 | 0.4961 |
| **ERR3239643** | NA11932 | CEU | EUR | EUR | 2,636,597 | 2,595,220 | 0.4950 |
| **ERR3240170** | HG00183 | FIN | EUR | EUR | 2,634,112 | 2,592,974 | 0.4944 |
| **ERR3240176** | HG00190 | FIN | EUR | EUR | 2,640,560 | 2,599,453 | 0.4944 |
| **ERR3240231** | HG00308 | FIN | EUR | EUR | 2,602,565 | 2,559,963 | 0.4953 |
| **ERR3240255** | HG00351 | FIN | EUR | EUR | 2,657,287 | 2,615,144 | 0.4928 |
| **ERR3240260** | HG00358 | FIN | EUR | EUR | 2,655,974 | 2,614,280 | 0.4921 |
| **ERR3240266** | HG00366 | FIN | EUR | EUR | 2,649,661 | 2,607,979 | 0.4937 |
| **ERR3241786** | HG00277 | FIN | EUR | EUR | 2,622,037 | 2,581,315 | 0.4957 |
| **ERR3241793** | HG00310 | FIN | EUR | EUR | 2,624,698 | 2,584,007 | 0.4952 |
| **ERR3241795** | HG00325 | FIN | EUR | EUR | 2,641,685 | 2,600,695 | 0.4945 |
| **ERR3241802** | HG00341 | FIN | EUR | EUR | 2,630,414 | 2,583,433 | 0.4923 |
| **ERR3240123** | HG00107 | GBR | EUR | EUR | 2,616,699 | 2,577,380 | 0.4974 |
| **ERR3240130** | HG00114 | GBR | EUR | EUR | 2,627,487 | 2,586,688 | 0.4957 |
| **ERR3240142** | HG00136 | GBR | EUR | EUR | 2,623,098 | 2,582,788 | 0.4963 |
| **ERR3240144** | HG00138 | GBR | EUR | EUR | 2,650,144 | 2,609,268 | 0.4941 |

| | | | | | | | |
|---|---|---|---|---|---|---|---|
| **ERR3240147** | HG00141 | GBR | EUR | EUR | 2,615,804 | 2,575,567 | 0.4964 |
| **ERR3240150** | HG00145 | GBR | EUR | EUR | 2,605,787 | 2,564,751 | 0.4971 |
| **ERR3240152** | HG00148 | GBR | EUR | EUR | 2,608,382 | 2,568,049 | 0.4966 |
| **ERR3240156** | HG00155 | GBR | EUR | EUR | 2,603,938 | 2,563,680 | 0.4978 |
| **ERR3240157** | HG00157 | GBR | EUR | EUR | 2,662,884 | 2,621,612 | 0.4947 |
| **ERR3240199** | HG00126 | GBR | EUR | EUR | 2,642,676 | 2,602,318 | 0.4951 |
| **ERR3241920** | HG01506 | IBS | EUR | EUR | 2,625,902 | 2,572,914 | 0.4914 |
| **ERR3241930** | HG01521 | IBS | EUR | EUR | 2,603,334 | 2,561,611 | 0.4977 |
| **ERR3241938** | HG01536 | IBS | EUR | EUR | 2,647,249 | 2,597,681 | 0.4905 |
| **ERR3241947** | HG01608 | IBS | EUR | EUR | 2,619,911 | 2,567,829 | 0.4914 |
| **ERR3241952** | HG01617 | IBS | EUR | EUR | 2,628,237 | 2,576,407 | 0.4910 |
| **ERR3241961** | HG01630 | IBS | EUR | EUR | 2,641,893 | 2,576,645 | 0.4841 |
| **ERR3241962** | HG01631 | IBS | EUR | EUR | 2,605,974 | 2,561,397 | 0.4955 |
| **ERR3241967** | HG01672 | IBS | EUR | EUR | 2,625,316 | 2,577,958 | 0.4925 |
| **ERR3241971** | HG01678 | IBS | EUR | EUR | 2,606,650 | 2,559,642 | 0.4945 |
| **ERR3241978** | HG01700 | IBS | EUR | EUR | 2,593,540 | 2,551,064 | 0.4968 |
| **ERR3239795** | NA20512 | TSI | EUR | EUR | 2,645,142 | 2,600,933 | 0.4934 |
| **ERR3239799** | NA20516 | TSI | EUR | EUR | 2,636,281 | 2,592,586 | 0.4938 |
| **ERR3239806** | NA20524 | TSI | EUR | EUR | 2,645,857 | 2,602,317 | 0.4936 |
| **ERR3239809** | NA20528 | TSI | EUR | EUR | 2,603,839 | 2,557,156 | 0.4942 |
| **ERR3239813** | NA20532 | TSI | EUR | EUR | 2,600,291 | 2,555,513 | 0.4951 |
| **ERR3239824** | NA20544 | TSI | EUR | EUR | 2,635,784 | 2,591,811 | 0.4930 |
| **ERR3239835** | NA20755 | TSI | EUR | EUR | 2,653,922 | 2,609,649 | 0.4919 |
| **ERR3239843** | NA20763 | TSI | EUR | EUR | 2,662,212 | 2,616,493 | 0.4924 |
| **ERR3239856** | NA20778 | TSI | EUR | EUR | 2,649,875 | 2,605,475 | 0.4929 |
| **ERR3239882** | NA20814 | TSI | EUR | EUR | 2,653,039 | 2,607,218 | 0.4914 |
| **ERR3242653** | HG03824 | BEB | SAS | SAS | 2,652,207 | 2,607,521 | 0.4928 |
| **ERR3242660** | HG03914 | BEB | SAS | SAS | 2,660,181 | 2,616,934 | 0.4929 |
| **ERR3242840** | HG03809 | BEB | SAS | SAS | 2,704,732 | 2,661,438 | 0.4918 |
| **ERR3242842** | HG03009 | BEB | SAS | SAS | 2,672,611 | 2,629,858 | 0.4928 |
| **ERR3242845** | HG03902 | BEB | SAS | SAS | 2,728,678 | 2,684,967 | 0.4881 |
| **ERR3242854** | HG03585 | BEB | SAS | SAS | 2,696,732 | 2,651,785 | 0.4904 |
| **ERR3242860** | HG03593 | BEB | SAS | SAS | 2,687,971 | 2,645,036 | 0.4918 |
| **ERR3243067** | HG03821 | BEB | SAS | SAS | 2,679,757 | 2,637,002 | 0.4917 |
| **ERR3243070** | HG03600 | BEB | SAS | SAS | 2,658,259 | 2,620,724 | 0.4957 |
| **ERR3243071** | HG03603 | BEB | SAS | SAS | 2,709,493 | 2,666,785 | 0.4910 |
| **ERR3240003** | NA20852 | GIH | SAS | SAS | 2,645,171 | 2,605,783 | 0.4943 |
| **ERR3240013** | NA20870 | GIH | SAS | SAS | 2,661,167 | 2,617,930 | 0.4905 |
| **ERR3240027** | NA20889 | GIH | SAS | SAS | 2,692,850 | 2,655,284 | 0.4931 |
| **ERR3240028** | NA20890 | GIH | SAS | SAS | 2,662,659 | 2,621,135 | 0.4916 |
| **ERR3240034** | NA20897 | GIH | SAS | SAS | 2,671,429 | 2,633,128 | 0.4945 |
| **ERR3240040** | NA20904 | GIH | SAS | SAS | 2,708,734 | 2,666,694 | 0.4891 |
| **ERR3240048** | NA21090 | GIH | SAS | SAS | 2,681,326 | 2,644,474 | 0.4924 |
| **ERR3240059** | NA21104 | GIH | SAS | SAS | 2,663,310 | 2,625,738 | 0.4939 |
| **ERR3240103** | NA20864 | GIH | SAS | SAS | 2,704,737 | 2,663,714 | 0.4903 |
| **ERR3240109** | NA21095 | GIH | SAS | SAS | 2,605,740 | 2,567,500 | 0.4968 |
| **ERR3242711** | HG03775 | ITU | SAS | SAS | 2,678,841 | 2,641,804 | 0.4933 |
| **ERR3242720** | HG03869 | ITU | SAS | SAS | 2,697,305 | 2,659,594 | 0.4920 |
| **ERR3242912** | HG03718 | ITU | SAS | SAS | 2,642,393 | 2,606,312 | 0.4956 |
| **ERR3242917** | HG03785 | ITU | SAS | SAS | 2,657,891 | 2,622,265 | 0.4951 |
| **ERR3242926** | HG03867 | ITU | SAS | SAS | 2,650,886 | 2,613,634 | 0.4949 |
| **ERR3242930** | HG03720 | ITU | SAS | SAS | 2,681,784 | 2,644,343 | 0.4930 |
| **ERR3242935** | HG04015 | ITU | SAS | SAS | 2,676,290 | 2,639,040 | 0.4936 |
| **ERR3243115** | HG03969 | ITU | SAS | SAS | 2,701,366 | 2,664,008 | 0.4929 |
| **ERR3243145** | HG03875 | ITU | SAS | SAS | 2,685,899 | 2,646,326 | 0.4909 |
| **ERR3243149** | HG03871 | ITU | SAS | SAS | 2,682,697 | 2,645,506 | 0.4933 |
| **ERR3242415** | HG02733 | PJL | SAS | SAS | 2,664,741 | 2,623,744 | 0.4918 |
| **ERR3242501** | HG02724 | PJL | SAS | SAS | 2,690,821 | 2,651,704 | 0.4924 |
| **ERR3242505** | HG02783 | PJL | SAS | SAS | 2,686,933 | 2,648,537 | 0.4926 |
| **ERR3242636** | HG03015 | PJL | SAS | SAS | 2,681,531 | 2,643,282 | 0.4928 |
| **ERR3243009** | HG02493 | PJL | SAS | SAS | 2,686,277 | 2,642,281 | 0.4897 |
| **ERR3243013** | HG02651 | PJL | SAS | SAS | 2,676,535 | 2,635,209 | 0.4915 |
| **ERR3243015** | HG02681 | PJL | SAS | SAS | 2,662,687 | 2,620,964 | 0.4913 |
| **ERR3243023** | HG02774 | PJL | SAS | SAS | 2,653,644 | 2,613,378 | 0.4926 |
| **ERR3243029** | HG03624 | PJL | SAS | SAS | 2,659,315 | 2,616,840 | 0.4910 |
| **ERR3243047** | HG01589 | PJL | SAS | SAS | 2,645,592 | 2,603,620 | 0.4923 |

| | | | | | | | |
|---|---|---|---|---|---|---|---|
| **ERR3242668** | HG03679 | STU | SAS | SAS | 2,668,802 | 2,630,742 | 0.4933 |
| **ERR3242675** | HG03693 | STU | SAS | SAS | 2,701,970 | 2,664,741 | 0.4930 |
| **ERR3242863** | HG03755 | STU | SAS | SAS | 2,706,007 | 2,667,816 | 0.4910 |
| **ERR3242864** | HG03850 | STU | SAS | SAS | 2,681,238 | 2,643,482 | 0.4932 |
| **ERR3242874** | HG03685 | STU | SAS | SAS | 2,662,519 | 2,623,703 | 0.4941 |
| **ERR3242880** | HG03694 | STU | SAS | SAS | 2,686,169 | 2,640,580 | 0.4905 |
| **ERR3242885** | HG03711 | STU | SAS | SAS | 2,656,401 | 2,618,904 | 0.4942 |
| **ERR3242889** | HG03744 | STU | SAS | SAS | 2,699,962 | 2,662,049 | 0.4925 |
| **ERR3242895** | HG03837 | STU | SAS | SAS | 2,604,773 | 2,568,134 | 0.4968 |
| **ERR3242901** | HG03890 | STU | SAS | SAS | 2,717,308 | 2,680,085 | 0.4910 |

**Table S5.** ntRoot super-population level LAI-based ancestry fraction estimates on the 1kGP WGS validation set using 5Mbp (5Mbt) or 2Mbp (2Mbt) tiles. The super-populations are EAS: East Asian, SAS: South Asian, AMR: Admixed American, AFR: African, EUR: European.

| Accession | Alternate ID | Population Code | LAI EAS fraction 5Mbt (%) | LAI SAS fraction 5Mbt (%) | LAI AMR fraction 5Mbt (%) | LAI AFR fraction 5Mbt (%) | LAI EUR fraction 5Mbt (%) | 1kGP Assigned Label | GAI Predicted from LAI, 5Mbt | GAI Predicted from LAI, 2Mbt |
|---|---|---|---|---|---|---|---|---|---|---|
| **ERR3242189** | HG01882 | ACB | 0.18 | 0.18 | 1.58 | 97.72 | 0.35 | AFR | AFR | AFR |
| **ERR3242193** | HG02014 | ACB | 0.18 | 0.00 | 4.20 | 95.45 | 0.18 | AFR | AFR | AFR |
| **ERR3242218** | HG01914 | ACB | 0.00 | 0.35 | 6.48 | 91.77 | 1.40 | AFR | AFR | AFR |
| **ERR3242297** | HG01988 | ACB | 0.18 | 1.58 | 12.61 | 74.78 | 10.86 | AFR | AFR | AFR |
| **ERR3242299** | HG02051 | ACB | 0.18 | 0.00 | 0.35 | 99.47 | 0.00 | AFR | AFR | AFR |
| **ERR3242306** | HG02143 | ACB | 0.35 | 0.35 | 5.95 | 92.12 | 1.23 | AFR | AFR | AFR |
| **ERR3242308** | HG02307 | ACB | 0.00 | 0.00 | 1.93 | 98.07 | 0.00 | AFR | AFR | AFR |
| **ERR3242318** | HG02330 | ACB | 0.00 | 0.00 | 5.60 | 93.87 | 0.53 | AFR | AFR | AFR |
| **ERR3242321** | HG02419 | ACB | 0.70 | 0.00 | 3.33 | 95.80 | 0.18 | AFR | AFR | AFR |
| **ERR3242322** | HG02420 | ACB | 0.00 | 1.40 | 12.78 | 80.56 | 5.25 | AFR | AFR | AFR |
| **ERR3242326** | HG02442 | ACB | 0.00 | 0.00 | 0.35 | 99.65 | 0.00 | AFR | AFR | AFR |
| **ERR3242740** | HG02541 | ACB | 0.00 | 0.35 | 4.38 | 92.82 | 2.45 | AFR | AFR | AFR |
| **ERR3239918** | NA19711 | ASW | 0.53 | 0.35 | 2.80 | 95.97 | 0.35 | AFR | AFR | AFR |
| **ERR3239961** | NA19834 | ASW | 0.18 | 0.18 | 10.51 | 88.44 | 0.70 | AFR | AFR | AFR |
| **ERR3239963** | NA19900 | ASW | 0.18 | 0.18 | 4.55 | 92.99 | 2.10 | AFR | AFR | AFR |
| **ERR3239965** | NA19904 | ASW | 0.00 | 0.70 | 6.83 | 91.24 | 1.23 | AFR | AFR | AFR |
| **ERR3239973** | NA19982 | ASW | 0.18 | 0.00 | 2.28 | 97.02 | 0.53 | AFR | AFR | AFR |
| **ERR3239974** | NA20126 | ASW | 0.70 | 1.23 | 7.88 | 85.11 | 5.08 | AFR | AFR | AFR |
| **ERR3239982** | NA20291 | ASW | 0.00 | 0.18 | 0.53 | 99.30 | 0.00 | AFR | AFR | AFR |
| **ERR3239990** | NA20340 | ASW | 0.18 | 0.18 | 5.95 | 91.94 | 1.75 | AFR | AFR | AFR |
| **ERR3239993** | NA20348 | ASW | 0.18 | 0.70 | 5.08 | 92.64 | 1.40 | AFR | AFR | AFR |
| **ERR3240095** | NA19922 | ASW | 0.18 | 1.23 | 8.41 | 85.46 | 4.73 | AFR | AFR | AFR |
| **ERR3242481** | HG02923 | ESN | 0.00 | 0.18 | 0.70 | 99.12 | 0.00 | AFR | AFR | AFR |
| **ERR3242487** | HG03100 | ESN | 0.00 | 0.18 | 0.53 | 99.30 | 0.00 | AFR | AFR | AFR |
| **ERR3242587** | HG03268 | ESN | 0.00 | 0.00 | 0.18 | 99.82 | 0.00 | AFR | AFR | AFR |
| **ERR3242589** | HG03271 | ESN | 0.00 | 0.00 | 0.18 | 99.82 | 0.00 | AFR | AFR | AFR |
| **ERR3242612** | HG03160 | ESN | 0.00 | 0.00 | 0.00 | 100.00 | 0.00 | AFR | AFR | AFR |
| **ERR3242623** | HG03343 | ESN | 0.00 | 0.00 | 0.35 | 99.65 | 0.00 | AFR | AFR | AFR |
| **ERR3242982** | HG03127 | ESN | 0.00 | 0.00 | 0.18 | 99.82 | 0.00 | AFR | AFR | AFR |
| **ERR3242986** | HG03175 | ESN | 0.00 | 0.00 | 0.00 | 100.00 | 0.00 | AFR | AFR | AFR |
| **ERR3242991** | HG03265 | ESN | 0.00 | 0.00 | 0.53 | 99.47 | 0.00 | AFR | AFR | AFR |
| **ERR3242993** | HG03298 | ESN | 0.00 | 0.00 | 0.18 | 99.82 | 0.00 | AFR | AFR | AFR |
| **ERR3242357** | HG02461 | GWD | 0.00 | 0.00 | 0.35 | 99.65 | 0.00 | AFR | AFR | AFR |
| **ERR3242365** | HG02573 | GWD | 0.18 | 0.00 | 0.18 | 99.65 | 0.00 | AFR | AFR | AFR |
| **ERR3242373** | HG02594 | GWD | 0.00 | 0.00 | 1.05 | 98.95 | 0.00 | AFR | AFR | AFR |
| **ERR3242379** | HG02623 | GWD | 0.00 | 0.00 | 0.35 | 99.65 | 0.00 | AFR | AFR | AFR |
| **ERR3242385** | HG02645 | GWD | 0.00 | 0.00 | 0.53 | 99.47 | 0.00 | AFR | AFR | AFR |
| **ERR3242391** | HG02678 | GWD | 0.00 | 0.18 | 1.05 | 98.77 | 0.00 | AFR | AFR | AFR |
| **ERR3242395** | HG02721 | GWD | 0.00 | 0.00 | 0.53 | 99.47 | 0.00 | AFR | AFR | AFR |
| **ERR3242423** | HG02768 | GWD | 0.00 | 0.00 | 0.70 | 99.30 | 0.00 | AFR | AFR | AFR |
| **ERR3242429** | HG02804 | GWD | 0.00 | 0.00 | 0.18 | 99.82 | 0.00 | AFR | AFR | AFR |
| **ERR3242457** | HG02816 | GWD | 0.00 | 0.00 | 1.23 | 98.60 | 0.18 | AFR | AFR | AFR |
| **ERR3239689** | NA19026 | LWK | 0.35 | 0.00 | 1.23 | 98.42 | 0.00 | AFR | AFR | AFR |
| **ERR3239698** | NA19041 | LWK | 0.00 | 0.00 | 0.70 | 99.30 | 0.00 | AFR | AFR | AFR |
| **ERR3239706** | NA19309 | LWK | 0.00 | 0.00 | 1.40 | 98.60 | 0.00 | AFR | AFR | AFR |
| **ERR3239708** | NA19312 | LWK | 0.00 | 0.00 | 0.88 | 99.12 | 0.00 | AFR | AFR | AFR |
| **ERR3239714** | NA19319 | LWK | 0.18 | 0.00 | 0.35 | 99.47 | 0.00 | AFR | AFR | AFR |
| **ERR3239726** | NA19347 | LWK | 0.18 | 0.00 | 0.53 | 99.30 | 0.00 | AFR | AFR | AFR |
| **ERR3239727** | NA19350 | LWK | 0.00 | 0.00 | 1.40 | 98.60 | 0.00 | AFR | AFR | AFR |
| **ERR3239738** | NA19380 | LWK | 0.00 | 0.00 | 1.40 | 98.60 | 0.00 | AFR | AFR | AFR |
| **ERR3239744** | NA19393 | LWK | 0.00 | 0.00 | 0.53 | 99.47 | 0.00 | AFR | AFR | AFR |
| **ERR3239752** | NA19428 | LWK | 0.00 | 0.00 | 0.88 | 99.12 | 0.00 | AFR | AFR | AFR |
| **ERR3239768** | NA19451 | LWK | 0.00 | 0.35 | 0.88 | 98.77 | 0.00 | AFR | AFR | AFR |
| **ERR3242471** | HG03084 | MSL | 0.00 | 0.00 | 0.35 | 99.47 | 0.18 | AFR | AFR | AFR |
| **ERR3242476** | HG03451 | MSL | 0.00 | 0.00 | 0.53 | 99.30 | 0.18 | AFR | AFR | AFR |
| **ERR3242551** | HG03439 | MSL | 0.00 | 0.00 | 0.70 | 99.30 | 0.00 | AFR | AFR | AFR |
| **ERR3242559** | HG03066 | MSL | 0.00 | 0.00 | 0.35 | 99.65 | 0.00 | AFR | AFR | AFR |

| | | | | | | | | | | |
|---|---|---|---|---|---|---|---|---|---|---|
| **ERR3242568** | HG03484 | MSL | 0.00 | 0.00 | 0.18 | 99.82 | 0.00 | AFR | AFR | AFR |
| **ERR3242570** | HG03547 | MSL | 0.00 | 0.18 | 0.70 | 99.12 | 0.00 | AFR | AFR | AFR |
| **ERR3242578** | HG03565 | MSL | 0.00 | 0.00 | 0.53 | 99.47 | 0.00 | AFR | AFR | AFR |
| **ERR3242952** | HG03060 | MSL | 0.00 | 0.00 | 0.53 | 99.47 | 0.00 | AFR | AFR | AFR |
| **ERR3242966** | HG03382 | MSL | 0.00 | 0.00 | 0.18 | 99.82 | 0.00 | AFR | AFR | AFR |
| **ERR3242968** | HG03432 | MSL | 0.00 | 0.18 | 0.35 | 99.47 | 0.00 | AFR | AFR | AFR |
| **ERR3239341** | NA18504 | YRI | 0.00 | 0.00 | 0.18 | 99.82 | 0.00 | AFR | AFR | AFR |
| **ERR3239349** | NA18519 | YRI | 0.00 | 0.00 | 0.53 | 99.47 | 0.00 | AFR | AFR | AFR |
| **ERR3239387** | NA18871 | YRI | 0.00 | 0.00 | 0.35 | 99.65 | 0.00 | AFR | AFR | AFR |
| **ERR3239429** | NA19119 | YRI | 0.00 | 0.00 | 0.70 | 99.30 | 0.00 | AFR | AFR | AFR |
| **ERR3239551** | NA18910 | YRI | 0.18 | 0.00 | 0.35 | 99.47 | 0.00 | AFR | AFR | AFR |
| **ERR3239556** | NA18934 | YRI | 0.00 | 0.00 | 0.53 | 99.47 | 0.00 | AFR | AFR | AFR |
| **ERR3239619** | NA19096 | YRI | 0.00 | 0.18 | 0.53 | 99.30 | 0.00 | AFR | AFR | AFR |
| **ERR3239622** | NA19117 | YRI | 0.00 | 0.00 | 0.70 | 99.30 | 0.00 | AFR | AFR | AFR |
| **ERR3239626** | NA19146 | YRI | 0.00 | 0.00 | 0.70 | 99.30 | 0.00 | AFR | AFR | AFR |
| **ERR3239629** | NA19184 | YRI | 0.00 | 0.00 | 0.70 | 99.30 | 0.00 | AFR | AFR | AFR |
| **ERR3241834** | HG01136 | CLM | 4.90 | 4.03 | 51.31 | 16.99 | 22.77 | AMR | AMR | AMR |
| **ERR3241862** | HG01250 | CLM | 7.36 | 4.20 | 54.29 | 14.19 | 19.96 | AMR | AMR | AMR |
| **ERR3241875** | HG01344 | CLM | 9.81 | 3.50 | 56.39 | 9.46 | 20.84 | AMR | AMR | AMR |
| **ERR3241878** | HG01356 | CLM | 6.48 | 4.20 | 53.59 | 12.61 | 23.12 | AMR | AMR | AMR |
| **ERR3241888** | HG01353 | CLM | 3.15 | 5.08 | 46.94 | 9.46 | 35.38 | AMR | AMR | AMR |
| **ERR3241894** | HG01374 | CLM | 2.98 | 7.01 | 53.06 | 6.65 | 30.30 | AMR | AMR | AMR |
| **ERR3241898** | HG01437 | CLM | 3.68 | 6.65 | 39.93 | 8.23 | 41.51 | AMR | EUR | EUR |
| **ERR3241904** | HG01461 | CLM | 10.16 | 2.98 | 50.96 | 26.27 | 9.63 | AMR | AMR | AMR |
| **ERR3242787** | HG01130 | CLM | 5.08 | 4.20 | 49.74 | 14.19 | 26.80 | AMR | AMR | AMR |
| **ERR3242791** | HG01280 | CLM | 4.38 | 4.90 | 56.92 | 7.71 | 26.09 | AMR | AMR | AMR |
| **ERR3239897** | NA19652 | MXL | 5.60 | 6.30 | 50.79 | 11.56 | 25.74 | AMR | AMR | AMR |
| **ERR3239902** | NA19661 | MXL | 14.54 | 3.85 | 57.79 | 11.73 | 12.08 | AMR | AMR | AMR |
| **ERR3239904** | NA19664 | MXL | 16.29 | 3.50 | 59.54 | 11.03 | 9.63 | AMR | AMR | AMR |
| **ERR3239911** | NA19682 | MXL | 14.89 | 5.43 | 59.72 | 5.95 | 14.01 | AMR | AMR | AMR |
| **ERR3239928** | NA19726 | MXL | 17.34 | 2.98 | 66.02 | 5.60 | 8.06 | AMR | AMR | AMR |
| **ERR3239940** | NA19759 | MXL | 22.07 | 1.75 | 66.02 | 6.13 | 4.03 | AMR | AMR | AMR |
| **ERR3239946** | NA19774 | MXL | 10.86 | 3.33 | 59.19 | 12.26 | 14.36 | AMR | AMR | AMR |
| **ERR3239950** | NA19780 | MXL | 8.76 | 3.15 | 59.72 | 8.41 | 19.96 | AMR | AMR | AMR |
| **ERR3239956** | NA19789 | MXL | 11.56 | 5.08 | 58.67 | 11.91 | 12.78 | AMR | AMR | AMR |
| **ERR3240091** | NA19741 | MXL | 28.55 | 1.93 | 64.10 | 3.68 | 1.75 | AMR | AMR | AMR |
| **ERR3240094** | NA19792 | MXL | 7.53 | 6.30 | 54.82 | 12.61 | 18.74 | AMR | AMR | AMR |
| **ERR3241988** | HG01577 | PEL | 9.46 | 4.03 | 60.95 | 13.31 | 12.26 | AMR | AMR | AMR |
| **ERR3241996** | HG01923 | PEL | 29.95 | 1.05 | 66.20 | 2.28 | 0.53 | AMR | AMR | AMR |
| **ERR3241998** | HG01926 | PEL | 32.92 | 0.53 | 64.97 | 1.40 | 0.18 | AMR | AMR | AMR |
| **ERR3242004** | HG01938 | PEL | 29.77 | 1.05 | 66.20 | 1.93 | 1.05 | AMR | AMR | AMR |
| **ERR3242010** | HG01947 | PEL | 15.41 | 2.28 | 68.48 | 7.71 | 6.13 | AMR | AMR | AMR |
| **ERR3242018** | HG01970 | PEL | 17.86 | 3.50 | 69.53 | 3.33 | 5.78 | AMR | AMR | AMR |
| **ERR3242024** | HG01979 | PEL | 19.61 | 4.38 | 66.20 | 4.73 | 5.08 | AMR | AMR | AMR |
| **ERR3242028** | HG02008 | PEL | 26.27 | 1.58 | 64.62 | 4.38 | 3.15 | AMR | AMR | AMR |
| **ERR3242241** | HG02271 | PEL | 32.57 | 1.05 | 64.80 | 1.23 | 0.35 | AMR | AMR | AMR |
| **ERR3242811** | HG01961 | PEL | 31.17 | 0.53 | 67.95 | 0.35 | 0.00 | AMR | AMR | AMR |
| **ERR3241726** | HG00637 | PUR | 2.28 | 5.43 | 44.13 | 23.82 | 24.34 | AMR | AMR | AMR |
| **ERR3241728** | HG00640 | PUR | 3.68 | 8.06 | 41.33 | 14.19 | 32.75 | AMR | AMR | EUR |
| **ERR3241763** | HG01054 | PUR | 3.50 | 4.73 | 47.46 | 14.36 | 29.95 | AMR | AMR | AMR |
| **ERR3241811** | HG01072 | PUR | 4.03 | 5.95 | 47.64 | 14.01 | 28.37 | AMR | AMR | AMR |
| **ERR3241816** | HG01085 | PUR | 2.63 | 5.08 | 42.73 | 21.02 | 28.55 | AMR | AMR | AMR |
| **ERR3241822** | HG01104 | PUR | 3.15 | 6.13 | 44.48 | 14.01 | 32.22 | AMR | AMR | AMR |
| **ERR3241844** | HG01173 | PUR | 3.15 | 7.71 | 39.93 | 13.49 | 35.73 | AMR | AMR | EUR |
| **ERR3241854** | HG01197 | PUR | 3.68 | 5.60 | 44.31 | 15.24 | 31.17 | AMR | AMR | AMR |
| **ERR3242774** | HG01161 | PUR | 3.68 | 3.50 | 45.01 | 27.67 | 20.14 | AMR | AMR | AMR |
| **ERR3242776** | HG01164 | PUR | 2.10 | 5.78 | 44.13 | 17.86 | 30.12 | AMR | AMR | AMR |
| **ERR3242164** | HG01816 | CDX | 98.25 | 0.18 | 0.70 | 0.70 | 0.18 | EAS | EAS | EAS |
| **ERR3242214** | HG02373 | CDX | 97.72 | 0.70 | 0.35 | 1.23 | 0.00 | EAS | EAS | EAS |
| **ERR3242216** | HG02396 | CDX | 97.72 | 0.53 | 0.88 | 0.53 | 0.35 | EAS | EAS | EAS |
| **ERR3242217** | HG02397 | CDX | 97.55 | 0.88 | 0.70 | 0.88 | 0.00 | EAS | EAS | EAS |
| **ERR3242262** | HG02353 | CDX | 97.02 | 0.70 | 0.88 | 1.23 | 0.18 | EAS | EAS | EAS |
| **ERR3242265** | HG02360 | CDX | 97.90 | 0.35 | 0.88 | 0.53 | 0.35 | EAS | EAS | EAS |
| **ERR3242275** | HG02384 | CDX | 97.02 | 0.70 | 1.05 | 0.88 | 0.35 | EAS | EAS | EAS |
| **ERR3242279** | HG02390 | CDX | 97.37 | 0.88 | 0.18 | 1.40 | 0.18 | EAS | EAS | EAS |
| **ERR3242285** | HG02401 | CDX | 97.20 | 0.70 | 0.88 | 1.23 | 0.00 | EAS | EAS | EAS |

| | | | | | | | | | | |
|---|---|---|---|---|---|---|---|---|---|---|
| **ERR3242289** | HG02409 | CDX | 97.02 | 1.05 | 1.23 | 0.35 | 0.35 | EAS | EAS | EAS |
| **ERR3239362** | NA18558 | CHB | 97.02 | 1.05 | 0.88 | 0.88 | 0.18 | EAS | EAS | EAS |
| **ERR3239363** | NA18561 | CHB | 95.45 | 0.88 | 1.93 | 1.58 | 0.18 | EAS | EAS | EAS |
| **ERR3239370** | NA18572 | CHB | 98.07 | 0.18 | 1.05 | 0.53 | 0.18 | EAS | EAS | EAS |
| **ERR3239378** | NA18605 | CHB | 97.55 | 0.70 | 0.35 | 1.05 | 0.35 | EAS | EAS | EAS |
| **ERR3239379** | NA18608 | CHB | 96.15 | 1.23 | 1.05 | 1.40 | 0.18 | EAS | EAS | EAS |
| **ERR3239496** | NA18544 | CHB | 96.67 | 1.23 | 1.05 | 0.53 | 0.53 | EAS | EAS | EAS |
| **ERR3239501** | NA18559 | CHB | 97.55 | 0.35 | 0.53 | 0.88 | 0.70 | EAS | EAS | EAS |
| **ERR3239510** | NA18612 | CHB | 97.55 | 0.70 | 1.23 | 0.35 | 0.18 | EAS | EAS | EAS |
| **ERR3239511** | NA18613 | CHB | 98.25 | 0.53 | 0.70 | 0.53 | 0.00 | EAS | EAS | EAS |
| **ERR3239534** | NA18639 | CHB | 98.42 | 0.35 | 0.35 | 0.88 | 0.00 | EAS | EAS | EAS |
| **ERR3239660** | NA18629 | CHB | 97.37 | 0.70 | 1.05 | 0.70 | 0.18 | EAS | EAS | EAS |
| **ERR3240183** | HG00442 | CHS | 97.72 | 0.35 | 0.70 | 0.88 | 0.35 | EAS | EAS | EAS |
| **ERR3241665** | HG00403 | CHS | 97.37 | 0.53 | 0.88 | 1.05 | 0.18 | EAS | EAS | EAS |
| **ERR3241673** | HG00436 | CHS | 97.90 | 0.88 | 0.53 | 0.70 | 0.00 | EAS | EAS | EAS |
| **ERR3241675** | HG00457 | CHS | 97.72 | 0.53 | 0.53 | 0.88 | 0.35 | EAS | EAS | EAS |
| **ERR3241681** | HG00478 | CHS | 97.20 | 0.70 | 0.88 | 1.23 | 0.00 | EAS | EAS | EAS |
| **ERR3241685** | HG00524 | CHS | 97.37 | 0.53 | 1.40 | 0.70 | 0.00 | EAS | EAS | EAS |
| **ERR3241697** | HG00556 | CHS | 97.20 | 0.70 | 0.53 | 1.40 | 0.18 | EAS | EAS | EAS |
| **ERR3241707** | HG00589 | CHS | 98.07 | 0.35 | 0.88 | 0.70 | 0.00 | EAS | EAS | EAS |
| **ERR3241715** | HG00610 | CHS | 98.07 | 0.70 | 0.35 | 0.88 | 0.00 | EAS | EAS | EAS |
| **ERR3241723** | HG00628 | CHS | 97.20 | 1.75 | 0.70 | 0.35 | 0.00 | EAS | EAS | EAS |
| **ERR3239401** | NA18952 | JPT | 97.37 | 0.70 | 0.88 | 1.05 | 0.00 | EAS | EAS | EAS |
| **ERR3239564** | NA18966 | JPT | 97.90 | 0.88 | 0.88 | 0.18 | 0.18 | EAS | EAS | EAS |
| **ERR3239574** | NA18995 | JPT | 96.85 | 0.88 | 0.88 | 1.05 | 0.35 | EAS | EAS | EAS |
| **ERR3239578** | NA19000 | JPT | 97.55 | 0.35 | 1.05 | 0.88 | 0.18 | EAS | EAS | EAS |
| **ERR3239583** | NA19009 | JPT | 97.20 | 0.53 | 0.88 | 1.05 | 0.35 | EAS | EAS | EAS |
| **ERR3239590** | NA19058 | JPT | 96.67 | 0.53 | 1.40 | 1.40 | 0.00 | EAS | EAS | EAS |
| **ERR3239673** | NA18982 | JPT | 97.37 | 0.70 | 0.35 | 1.23 | 0.35 | EAS | EAS | EAS |
| **ERR3239675** | NA18984 | JPT | 97.55 | 0.35 | 1.40 | 0.70 | 0.00 | EAS | EAS | EAS |
| **ERR3239678** | NA18988 | JPT | 97.90 | 0.18 | 0.88 | 1.05 | 0.00 | EAS | EAS | EAS |
| **ERR3239681** | NA19006 | JPT | 97.72 | 0.53 | 1.05 | 0.70 | 0.00 | EAS | EAS | EAS |
| **ERR3242036** | HG02017 | KHV | 97.02 | 0.88 | 0.70 | 1.05 | 0.35 | EAS | EAS | EAS |
| **ERR3242038** | HG02020 | KHV | 97.20 | 1.40 | 0.70 | 0.70 | 0.00 | EAS | EAS | EAS |
| **ERR3242043** | HG02029 | KHV | 96.67 | 0.70 | 0.88 | 1.58 | 0.18 | EAS | EAS | EAS |
| **ERR3242047** | HG02070 | KHV | 97.37 | 0.88 | 0.53 | 0.53 | 0.70 | EAS | EAS | EAS |
| **ERR3242049** | HG02073 | KHV | 97.55 | 0.53 | 0.70 | 1.23 | 0.00 | EAS | EAS | EAS |
| **ERR3242069** | HG01842 | KHV | 97.37 | 0.35 | 1.40 | 0.88 | 0.00 | EAS | EAS | EAS |
| **ERR3242079** | HG01852 | KHV | 97.20 | 0.53 | 1.40 | 0.70 | 0.18 | EAS | EAS | EAS |
| **ERR3242092** | HG01867 | KHV | 96.15 | 0.88 | 0.70 | 1.93 | 0.35 | EAS | EAS | EAS |
| **ERR3242210** | HG02122 | KHV | 96.32 | 1.05 | 1.23 | 1.40 | 0.00 | EAS | EAS | EAS |
| **ERR3242343** | HG02047 | KHV | 97.20 | 0.53 | 0.70 | 1.40 | 0.18 | EAS | EAS | EAS |
| **ERR3239281** | NA07051 | CEU | 1.40 | 11.03 | 11.38 | 6.30 | 69.88 | EUR | EUR | EUR |
| **ERR3239285** | NA10851 | CEU | 0.88 | 9.28 | 11.73 | 4.73 | 73.38 | EUR | EUR | EUR |
| **ERR3239286** | NA11829 | CEU | 1.05 | 10.16 | 12.08 | 5.43 | 71.28 | EUR | EUR | EUR |
| **ERR3239288** | NA11831 | CEU | 1.58 | 10.68 | 14.89 | 4.03 | 68.83 | EUR | EUR | EUR |
| **ERR3239294** | NA11919 | CEU | 1.05 | 10.33 | 13.13 | 5.78 | 69.70 | EUR | EUR | EUR |
| **ERR3239302** | NA12005 | CEU | 1.75 | 10.51 | 12.61 | 4.38 | 70.75 | EUR | EUR | EUR |
| **ERR3239309** | NA12155 | CEU | 1.40 | 9.81 | 13.31 | 4.73 | 70.75 | EUR | EUR | EUR |
| **ERR3239334** | NA12878 | CEU | 1.05 | 8.93 | 14.01 | 5.95 | 70.05 | EUR | EUR | EUR |
| **ERR3239462** | NA11893 | CEU | 1.75 | 9.46 | 12.78 | 5.08 | 70.93 | EUR | EUR | EUR |
| **ERR3239473** | NA12342 | CEU | 0.88 | 10.86 | 13.31 | 6.13 | 68.83 | EUR | EUR | EUR |
| **ERR3239643** | NA11932 | CEU | 1.05 | 9.11 | 14.54 | 5.95 | 69.35 | EUR | EUR | EUR |
| **ERR3240170** | HG00183 | FIN | 3.68 | 9.11 | 13.84 | 6.13 | 67.25 | EUR | EUR | EUR |
| **ERR3240176** | HG00190 | FIN | 2.98 | 10.68 | 12.08 | 5.78 | 68.48 | EUR | EUR | EUR |
| **ERR3240231** | HG00308 | FIN | 5.60 | 9.28 | 12.26 | 6.83 | 66.02 | EUR | EUR | EUR |
| **ERR3240255** | HG00351 | FIN | 4.90 | 10.33 | 12.43 | 5.08 | 67.25 | EUR | EUR | EUR |
| **ERR3240260** | HG00358 | FIN | 5.95 | 8.93 | 12.08 | 4.03 | 69.00 | EUR | EUR | EUR |
| **ERR3240266** | HG00366 | FIN | 4.73 | 9.46 | 12.61 | 4.90 | 68.30 | EUR | EUR | EUR |
| **ERR3241786** | HG00277 | FIN | 3.50 | 11.91 | 15.94 | 5.60 | 63.05 | EUR | EUR | EUR |
| **ERR3241793** | HG00310 | FIN | 4.90 | 9.11 | 12.26 | 4.55 | 69.18 | EUR | EUR | EUR |
| **ERR3241795** | HG00325 | FIN | 5.43 | 10.16 | 12.08 | 6.13 | 66.20 | EUR | EUR | EUR |
| **ERR3241802** | HG00341 | FIN | 2.80 | 12.61 | 14.19 | 6.65 | 63.75 | EUR | EUR | EUR |
| **ERR3240123** | HG00107 | GBR | 2.63 | 8.93 | 13.31 | 6.13 | 69.00 | EUR | EUR | EUR |
| **ERR3240130** | HG00114 | GBR | 1.40 | 8.76 | 14.36 | 5.60 | 69.88 | EUR | EUR | EUR |
| **ERR3240142** | HG00136 | GBR | 1.58 | 9.28 | 13.13 | 5.25 | 70.75 | EUR | EUR | EUR |

| | | | | | | | | | | |
|---|---|---|---|---|---|---|---|---|---|---|
| **ERR3240144** | HG00138 | GBR | 1.40 | 8.58 | 13.31 | 5.60 | 71.10 | EUR | EUR | EUR |
| **ERR3240147** | HG00141 | GBR | 1.75 | 8.58 | 13.31 | 4.38 | 71.98 | EUR | EUR | EUR |
| **ERR3240150** | HG00145 | GBR | 1.23 | 8.58 | 13.31 | 7.71 | 69.18 | EUR | EUR | EUR |
| **ERR3240152** | HG00148 | GBR | 1.75 | 7.53 | 11.38 | 6.65 | 72.68 | EUR | EUR | EUR |
| **ERR3240156** | HG00155 | GBR | 1.40 | 9.81 | 14.89 | 5.78 | 68.13 | EUR | EUR | EUR |
| **ERR3240157** | HG00157 | GBR | 1.75 | 9.98 | 14.54 | 5.25 | 68.48 | EUR | EUR | EUR |
| **ERR3240199** | HG00126 | GBR | 1.93 | 8.41 | 13.13 | 6.65 | 69.88 | EUR | EUR | EUR |
| **ERR3241920** | HG01506 | IBS | 1.75 | 8.93 | 14.89 | 9.81 | 64.62 | EUR | EUR | EUR |
| **ERR3241930** | HG01521 | IBS | 1.05 | 7.71 | 17.16 | 5.43 | 68.65 | EUR | EUR | EUR |
| **ERR3241938** | HG01536 | IBS | 1.40 | 8.06 | 16.99 | 9.63 | 63.92 | EUR | EUR | EUR |
| **ERR3241947** | HG01608 | IBS | 0.88 | 8.41 | 17.34 | 9.28 | 64.10 | EUR | EUR | EUR |
| **ERR3241952** | HG01617 | IBS | 1.58 | 9.28 | 16.99 | 8.76 | 63.40 | EUR | EUR | EUR |
| **ERR3241961** | HG01630 | IBS | 1.23 | 7.88 | 22.77 | 10.51 | 57.62 | EUR | EUR | EUR |
| **ERR3241962** | HG01631 | IBS | 1.40 | 8.93 | 15.76 | 5.43 | 68.48 | EUR | EUR | EUR |
| **ERR3241967** | HG01672 | IBS | 1.23 | 8.58 | 16.81 | 8.23 | 65.15 | EUR | EUR | EUR |
| **ERR3241971** | HG01678 | IBS | 2.10 | 8.23 | 16.11 | 7.88 | 65.67 | EUR | EUR | EUR |
| **ERR3241978** | HG01700 | IBS | 1.40 | 5.95 | 18.21 | 4.90 | 69.53 | EUR | EUR | EUR |
| **ERR3239795** | NA20512 | TSI | 1.23 | 11.38 | 13.49 | 7.36 | 66.55 | EUR | EUR | EUR |
| **ERR3239799** | NA20516 | TSI | 1.75 | 10.51 | 14.71 | 6.30 | 66.73 | EUR | EUR | EUR |
| **ERR3239806** | NA20524 | TSI | 1.75 | 10.86 | 14.89 | 6.48 | 66.02 | EUR | EUR | EUR |
| **ERR3239809** | NA20528 | TSI | 1.23 | 9.81 | 16.81 | 6.83 | 65.32 | EUR | EUR | EUR |
| **ERR3239813** | NA20532 | TSI | 1.58 | 10.51 | 15.41 | 7.18 | 65.32 | EUR | EUR | EUR |
| **ERR3239824** | NA20544 | TSI | 1.75 | 11.91 | 16.29 | 6.65 | 63.40 | EUR | EUR | EUR |
| **ERR3239835** | NA20755 | TSI | 0.88 | 12.61 | 14.36 | 6.65 | 65.50 | EUR | EUR | EUR |
| **ERR3239843** | NA20763 | TSI | 1.23 | 11.21 | 14.54 | 7.71 | 65.32 | EUR | EUR | EUR |
| **ERR3239856** | NA20778 | TSI | 1.40 | 14.36 | 13.84 | 7.53 | 62.87 | EUR | EUR | EUR |
| **ERR3239882** | NA20814 | TSI | 2.10 | 9.46 | 13.31 | 9.28 | 65.85 | EUR | EUR | EUR |
| **ERR3242653** | HG03824 | BEB | 21.37 | 65.67 | 4.03 | 5.25 | 3.68 | SAS | SAS | SAS |
| **ERR3242660** | HG03914 | BEB | 17.51 | 73.38 | 2.98 | 3.68 | 2.45 | SAS | SAS | SAS |
| **ERR3242840** | HG03809 | BEB | 17.16 | 72.50 | 3.68 | 3.68 | 2.98 | SAS | SAS | SAS |
| **ERR3242842** | HG03009 | BEB | 15.06 | 72.50 | 3.50 | 4.38 | 4.55 | SAS | SAS | SAS |
| **ERR3242845** | HG03902 | BEB | 16.29 | 72.33 | 3.85 | 3.68 | 3.85 | SAS | SAS | SAS |
| **ERR3242854** | HG03585 | BEB | 17.69 | 68.65 | 4.20 | 5.78 | 3.68 | SAS | SAS | SAS |
| **ERR3242860** | HG03593 | BEB | 17.51 | 72.50 | 3.50 | 3.68 | 2.80 | SAS | SAS | SAS |
| **ERR3243067** | HG03821 | BEB | 16.11 | 74.61 | 2.10 | 4.38 | 2.80 | SAS | SAS | SAS |
| **ERR3243070** | HG03600 | BEB | 8.93 | 83.19 | 1.75 | 4.38 | 1.75 | SAS | SAS | SAS |
| **ERR3243071** | HG03603 | BEB | 17.16 | 73.38 | 2.98 | 4.38 | 2.10 | SAS | SAS | SAS |
| **ERR3240003** | NA20852 | GIH | 7.18 | 80.21 | 3.33 | 3.85 | 5.43 | SAS | SAS | SAS |
| **ERR3240013** | NA20870 | GIH | 4.03 | 70.40 | 5.95 | 5.95 | 13.66 | SAS | SAS | SAS |
| **ERR3240027** | NA20889 | GIH | 5.08 | 86.51 | 2.28 | 3.50 | 2.63 | SAS | SAS | SAS |
| **ERR3240028** | NA20890 | GIH | 5.25 | 76.01 | 3.85 | 6.48 | 8.41 | SAS | SAS | SAS |
| **ERR3240034** | NA20897 | GIH | 9.46 | 82.31 | 3.15 | 4.03 | 1.05 | SAS | SAS | SAS |
| **ERR3240040** | NA20904 | GIH | 6.65 | 71.80 | 7.01 | 4.38 | 10.16 | SAS | SAS | SAS |
| **ERR3240048** | NA21090 | GIH | 5.25 | 87.57 | 1.40 | 3.33 | 2.45 | SAS | SAS | SAS |
| **ERR3240059** | NA21104 | GIH | 7.18 | 83.54 | 2.28 | 4.03 | 2.98 | SAS | SAS | SAS |
| **ERR3240103** | NA20864 | GIH | 4.90 | 77.76 | 3.50 | 5.43 | 8.41 | SAS | SAS | SAS |
| **ERR3240109** | NA21095 | GIH | 5.25 | 83.19 | 3.15 | 3.85 | 4.55 | SAS | SAS | SAS |
| **ERR3242711** | HG03775 | ITU | 7.53 | 83.01 | 1.40 | 5.25 | 2.80 | SAS | SAS | SAS |
| **ERR3242720** | HG03869 | ITU | 8.23 | 81.79 | 2.28 | 5.25 | 2.45 | SAS | SAS | SAS |
| **ERR3242912** | HG03718 | ITU | 7.18 | 88.09 | 1.23 | 2.63 | 0.88 | SAS | SAS | SAS |
| **ERR3242917** | HG03785 | ITU | 8.93 | 85.11 | 1.05 | 3.15 | 1.75 | SAS | SAS | SAS |
| **ERR3242926** | HG03867 | ITU | 13.13 | 78.11 | 1.58 | 4.20 | 2.98 | SAS | SAS | SAS |
| **ERR3242930** | HG03720 | ITU | 8.58 | 82.66 | 2.98 | 4.38 | 1.40 | SAS | SAS | SAS |
| **ERR3242935** | HG04015 | ITU | 7.53 | 85.46 | 1.93 | 3.33 | 1.75 | SAS | SAS | SAS |
| **ERR3243115** | HG03969 | ITU | 6.30 | 83.54 | 2.45 | 4.90 | 2.80 | SAS | SAS | SAS |
| **ERR3243145** | HG03875 | ITU | 6.13 | 78.81 | 3.15 | 5.25 | 6.65 | SAS | SAS | SAS |
| **ERR3243149** | HG03871 | ITU | 6.30 | 85.64 | 2.28 | 3.85 | 1.93 | SAS | SAS | SAS |
| **ERR3242415** | HG02733 | PJL | 5.08 | 76.88 | 4.03 | 4.90 | 9.11 | SAS | SAS | SAS |
| **ERR3242501** | HG02724 | PJL | 8.58 | 80.04 | 3.68 | 4.38 | 3.33 | SAS | SAS | SAS |
| **ERR3242505** | HG02783 | PJL | 8.41 | 81.44 | 2.10 | 4.38 | 3.68 | SAS | SAS | SAS |
| **ERR3242636** | HG03015 | PJL | 8.93 | 78.98 | 3.15 | 4.90 | 4.03 | SAS | SAS | SAS |
| **ERR3243009** | HG02493 | PJL | 4.73 | 76.53 | 3.33 | 7.36 | 8.06 | SAS | SAS | SAS |
| **ERR3243013** | HG02651 | PJL | 5.60 | 75.48 | 4.90 | 4.38 | 9.63 | SAS | SAS | SAS |
| **ERR3243015** | HG02681 | PJL | 5.25 | 71.28 | 5.95 | 5.43 | 12.08 | SAS | SAS | SAS |
| **ERR3243023** | HG02774 | PJL | 6.30 | 78.11 | 4.20 | 5.08 | 6.30 | SAS | SAS | SAS |
| **ERR3243029** | HG03624 | PJL | 2.98 | 73.03 | 6.83 | 5.60 | 11.56 | SAS | SAS | SAS |

| | | | | | | | | | | |
|---|---|---|---|---|---|---|---|---|---|---|
| **ERR3243047** | HG01589 | PJL | 5.78 | 73.20 | 5.25 | 4.20 | 11.56 | SAS | SAS | SAS |
| **ERR3242668** | HG03679 | STU | 6.83 | 84.41 | 3.33 | 3.50 | 1.93 | SAS | SAS | SAS |
| **ERR3242675** | HG03693 | STU | 9.11 | 82.49 | 2.28 | 3.50 | 2.63 | SAS | SAS | SAS |
| **ERR3242863** | HG03755 | STU | 8.23 | 82.31 | 2.45 | 4.90 | 2.10 | SAS | SAS | SAS |
| **ERR3242864** | HG03850 | STU | 8.06 | 83.19 | 2.28 | 4.38 | 2.10 | SAS | SAS | SAS |
| **ERR3242874** | HG03685 | STU | 6.83 | 83.36 | 2.98 | 5.25 | 1.58 | SAS | SAS | SAS |
| **ERR3242880** | HG03694 | STU | 12.96 | 68.65 | 4.73 | 6.83 | 6.83 | SAS | SAS | SAS |
| **ERR3242885** | HG03711 | STU | 7.71 | 83.19 | 2.10 | 4.90 | 2.10 | SAS | SAS | SAS |
| **ERR3242889** | HG03744 | STU | 9.63 | 81.09 | 1.93 | 5.78 | 1.58 | SAS | SAS | SAS |
| **ERR3242895** | HG03837 | STU | 8.41 | 81.09 | 3.33 | 4.90 | 2.28 | SAS | SAS | SAS |
| **ERR3242901** | HG03890 | STU | 8.93 | 81.44 | 2.28 | 4.38 | 2.98 | SAS | SAS | SAS |

**Table S6.** ntRoot super-population level LAI-based ancestry fraction estimates on the SGDP WGS discovery set. The LAI fraction estimates were generated using 5Mbp tiles. The super-populations are EAS: East Asian, SAS: South Asian, AMR: Admixed American, AFR: African, EUR: European.

| Accession | Dataset Fold Coverage | Library | Population | Region | LAI EAS Fraction (%) | LAI SAS Fraction (%) | LAI AMR Fraction (%) | LAI AFR Fraction (%) | LAI EUR Fraction (%) |
|---|---|---|---|---|---|---|---|---|---|
| **ERR1019043^** | 41.0 | LP6005441-DNA_F01 | BantuHerero | Africa | 0.00 | 0.18 | 0.18 | 99.65 | 0.00 |
| **ERR1347714** | 45.2 | LP6005443-DNA_E02 | BantuHerero | Africa | 0.00 | 0.00 | 0.18 | 99.82 | 0.00 |
| **ERR1019044^** | 37.3 | LP6005441-DNA_B02 | BantuKenya | Africa | 0.00 | 0.00 | 0.53 | 99.47 | 0.00 |
| **ERR1347680** | 50.7 | LP6005443-DNA_A01 | BantuKenya | Africa | 0.00 | 0.00 | 0.53 | 99.47 | 0.00 |
| **ERR1347723^** | 39.9 | LP6005443-DNA_F02 | BantuTswana | Africa | 0.00 | 0.00 | 0.35 | 99.65 | 0.00 |
| **ERR1347732** | 44.3 | LP6005443-DNA_G02 | BantuTswana | Africa | 0.00 | 0.00 | 0.18 | 99.82 | 0.00 |
| **ERR1025612^** | 44.3 | LP6005441-DNA_G02 | Biaka | Africa | 0.00 | 0.00 | 0.70 | 99.30 | 0.00 |
| **ERR1419161** | 43.7 | LP6005441-DNA_H02 | Biaka | Africa | 0.00 | 0.00 | 0.18 | 99.82 | 0.00 |
| **ERR1347694^** | 46.3 | LP6005443-DNA_B09 | Dinka | Africa | 0.18 | 0.00 | 0.35 | 99.47 | 0.00 |
| **ERR1395603** | 48.4 | SS6004480 | Dinka | Africa | 0.00 | 0.00 | 0.70 | 99.30 | 0.00 |
| **ERR1395606** | 46.7 | LP6005443-DNA_H08 | Dinka | Africa | 0.00 | 0.00 | 1.23 | 98.77 | 0.00 |
| **ERR1025621^** | 35.6 | LP6005442-DNA_A10 | Esan | Africa | 0.00 | 0.18 | 0.53 | 99.30 | 0.00 |
| **ERR1025622** | 41.9 | LP6005442-DNA_B10 | Esan | Africa | 0.00 | 0.00 | 0.35 | 99.65 | 0.00 |
| **ERR1347670^** | 40.8 | LP6005442-DNA_G10 | Gambian | Africa | 0.35 | 0.35 | 0.35 | 98.95 | 0.00 |
| **ERR1347677** | 39.0 | LP6005442-DNA_H10 | Gambian | Africa | 0.00 | 0.18 | 0.70 | 99.12 | 0.00 |
| **ERR1347737^** | 47.2 | LP6005443-DNA_G08 | Ju hoan North | Africa | 0.00 | 0.00 | 0.70 | 99.30 | 0.00 |
| **ERR1395596** | 42.7 | SS6004473 | Ju hoan North | Africa | 0.00 | 0.00 | 1.05 | 98.95 | 0.00 |
| **ERR1419096** | 38.7 | LP6005441-DNA_A11 | Ju hoan North | Africa | 0.00 | 0.00 | 1.23 | 98.77 | 0.00 |
| **ERR1419108** | 34.8 | LP6005441-DNA_B11 | Ju hoan North | Africa | 0.18 | 0.18 | 0.88 | 98.77 | 0.00 |
| **ERR1395572^** | 53.9 | LP6005592-DNA_C05 | Khomani San | Africa | 0.00 | 0.18 | 0.18 | 99.65 | 0.00 |
| **ERR1395588** | 48.7 | LP6005677-DNA_D03 | Khomani San | Africa | 0.00 | 0.00 | 1.05 | 98.77 | 0.18 |
| **ERR1347655^** | 36.3 | LP6005442-DNA_E11 | Luhya | Africa | 0.00 | 0.00 | 1.40 | 98.60 | 0.00 |
| **ERR1347662** | 39.0 | LP6005442-DNA_F11 | Luhya | Africa | 0.00 | 0.00 | 1.05 | 98.95 | 0.00 |
| **ERR1347660^** | 37.7 | LP6005442-DNA_F09 | Luo | Africa | 0.00 | 0.00 | 0.35 | 99.65 | 0.00 |
| **ERR1395591** | 38.9 | LP6005677-DNA_G01 | Luo | Africa | 0.00 | 0.00 | 1.40 | 98.60 | 0.00 |
| **ERR1025640^** | 35.1 | LP6005441-DNA_E07 | Mandenka | Africa | 0.00 | 0.00 | 0.53 | 99.47 | 0.00 |
| **ERR1395593** | 41.4 | SS6004470 | Mandenka | Africa | 0.00 | 0.00 | 0.53 | 99.12 | 0.35 |
| **ERR1419145** | 39.0 | LP6005441-DNA_F07 | Mandenka | Africa | 0.00 | 0.00 | 0.35 | 99.65 | 0.00 |
| **ERR1347716^** | 39.6 | LP6005443-DNA_E06 | Masai | Africa | 0.00 | 0.18 | 7.01 | 92.29 | 0.53 |
| **ERR1347726** | 48.7 | LP6005443-DNA_F06 | Masai | Africa | 0.18 | 0.35 | 6.48 | 92.47 | 0.53 |

| | | | | | | | | | |
|---|---|---|---|---|---|---|---|---|---|
| **ERR1019077^** | 38.1 | LP6005441-DNA_B08 | Mbuti | Africa | 0.00 | 0.00 | 0.35 | 99.65 | 0.00 |
| **ERR1395571** | 42.7 | LP6005592-DNA_C03 | Mbuti | Africa | 0.00 | 0.00 | 0.18 | 99.82 | 0.00 |
| **ERR1395594** | 41.5 | SS6004471 | Mbuti | Africa | 0.00 | 0.00 | 0.53 | 99.47 | 0.00 |
| **ERR1419093** | 39.5 | LP6005441-DNA_A08 | Mbuti | Africa | 0.00 | 0.00 | 0.53 | 99.47 | 0.00 |
| **ERR1347671^** | 40.8 | LP6005442-DNA_G11 | Mende | Africa | 0.18 | 0.00 | 0.35 | 99.47 | 0.00 |
| **ERR1347678** | 37.6 | LP6005442-DNA_H11 | Mende | Africa | 0.00 | 0.18 | 0.35 | 99.47 | 0.00 |
| **ERR1025601^** | 38.0 | LP6005441-DNA_G08 | Mozabite | Africa | 0.18 | 4.03 | 27.15 | 41.86 | 26.80 |
| **ERR1025602** | 40.4 | LP6005441-DNA_H08 | Mozabite | Africa | 0.53 | 3.33 | 27.67 | 50.44 | 18.04 |
| **ERR1395581^** | 47.9 | LP6005619-DNA_B01 | Saharawi | Africa | 0.35 | 7.01 | 30.65 | 40.81 | 21.19 |
| **ERR1395582** | 51.3 | LP6005619-DNA_C01 | Saharawi | Africa | 0.35 | 4.03 | 26.80 | 47.81 | 21.02 |
| **ERR1025600^** | 42.0 | LP6005442-DNA_D09 | Somali | Africa | 0.18 | 1.23 | 13.49 | 82.84 | 2.28 |
| **ERR1395598^** | 44.2 | SS6004475 | Yoruba | Africa | 0.00 | 0.00 | 0.18 | 99.82 | 0.00 |
| **ERR1419171** | 34.5 | LP6005442-DNA_A02 | Yoruba | Africa | 0.00 | 0.00 | 0.88 | 99.12 | 0.00 |
| **ERR1419179** | 36.2 | LP6005442-DNA_B02 | Yoruba | Africa | 0.00 | 0.18 | 0.35 | 99.47 | 0.00 |
| **ERR1395547^** | 59.3 | LP6005519-DNA_D01 | Chane | America | 34.85 | 0.35 | 63.57 | 0.70 | 0.53 |
| **ERR1025635^** | 46.6 | LP6005441-DNA_G06 | Karitiana | America | 36.25 | 0.70 | 61.47 | 1.58 | 0.00 |
| **ERR1025636** | 39.1 | LP6005441-DNA_H06 | Karitiana | America | 38.18 | 0.53 | 60.07 | 1.23 | 0.00 |
| **ERR1395599** | 39.6 | SS6004476 | Karitiana | America | 36.43 | 0.18 | 61.82 | 1.05 | 0.53 |
| **ERR1025642^** | 38.0 | LP6005441-DNA_G07 | Mayan | America | 36.60 | 0.53 | 60.95 | 1.75 | 0.18 |
| **ERR1419165** | 45.6 | LP6005441-DNA_H07 | Mayan | America | 35.20 | 0.53 | 62.35 | 1.75 | 0.18 |
| **ERR1347721^** | 47.8 | LP6005443-DNA_E11 | Mixe | America | 39.05 | 0.35 | 59.02 | 1.40 | 0.18 |
| **ERR1347730** | 47.4 | LP6005443-DNA_F11 | Mixe | America | 35.20 | 0.53 | 62.00 | 2.28 | 0.00 |
| **ERR1395602** | 45.6 | SS6004479 | Mixe | America | 38.53 | 0.18 | 59.72 | 1.23 | 0.35 |
| **ERR1347738^** | 45.1 | LP6005443-DNA_G11 | Mixtec | America | 32.40 | 1.75 | 61.12 | 3.68 | 1.05 |
| **ERR1395607** | 47.5 | LP6005443-DNA_H11 | Mixtec | America | 21.54 | 2.63 | 65.85 | 5.43 | 4.55 |
| **ERR1019067^** | 42.5 | LP6005441-DNA_B04 | Piapoco | America | 36.95 | 0.53 | 61.12 | 1.40 | 0.00 |
| **ERR1419090** | 41.4 | LP6005441-DNA_A04 | Piapoco | America | 35.03 | 0.70 | 63.75 | 0.53 | 0.00 |
| **ERR1025604^** | 38.4 | LP6005441-DNA_F10 | Pima | America | 41.86 | 0.70 | 56.04 | 1.23 | 0.18 |
| **ERR1419139‡** | 21.8 | LP6005441-DNA_E10 | Pima | America | 39.58 | 0.53 | 58.14 | 1.75 | 0.00 |
| **ERR1395558^** | 60.9 | LP6005519-DNA_G02 | Quechua | America | 34.85 | 0.88 | 63.22 | 0.70 | 0.35 |
| **ERR1395589** | 45.8 | LP6005677-DNA_E01 | Quechua | America | 33.80 | 0.53 | 64.80 | 0.70 | 0.18 |
| **ERR1395590** | 42.4 | LP6005677-DNA_F01 | Quechua | America | 31.35 | 1.05 | 66.02 | 1.58 | 0.00 |
| **ERR1019070^** | 41.9 | LP6005441-DNA_A12 | Surui | America | 39.40 | 0.53 | 58.32 | 1.75 | 0.00 |
| **ERR1019071** | 36.1 | LP6005441-DNA_B12 | Surui | America | 38.53 | 0.70 | 59.02 | 1.75 | 0.00 |
| **ERR1347686^** | 46.3 | LP6005443-DNA_A12 | Zapotec | America | 38.88 | 0.70 | 59.37 | 1.05 | 0.00 |
| **ERR1395622** | 39.8 | LP6005677-DNA_D01 | Zapotec | America | 32.40 | 0.53 | 64.45 | 2.28 | 0.35 |

| | | | | | | | | | |
|---|---|---|---|---|---|---|---|---|---|
| **ERR1347682^** | 47.0 | LP6005443-DNA_A03 | Aleut | CentralAsiaSiberia | 44.13 | 7.53 | 33.63 | 3.33 | 11.38 |
| **ERR1347740** | 38.8 | LP6005443-DNA_H02 | Aleut | CentralAsiaSiberia | 41.68 | 8.06 | 34.50 | 4.03 | 11.73 |
| **ERR1419200^** | 40.8 | LP6005442-DNA_F02 | Altaian | CentralAsiaSiberia | 74.78 | 8.93 | 8.23 | 3.50 | 4.55 |
| **ERR1347698^** | 47.3 | LP6005443-DNA_C03 | Chukchi | CentralAsiaSiberia | 39.23 | 11.03 | 29.95 | 6.13 | 13.66 |
| **ERR1347706^** | 46.1 | LP6005443-DNA_D03 | Eskimo Chaplin | CentralAsiaSiberia | 79.16 | 1.23 | 17.16 | 1.75 | 0.70 |
| **ERR1347724^** | 49.2 | LP6005443-DNA_F03 | Eskimo Naukan | CentralAsiaSiberia | 76.88 | 2.28 | 18.04 | 2.63 | 0.18 |
| **ERR1347733** | 46.8 | LP6005443-DNA_G03 | Eskimo Naukan | CentralAsiaSiberia | 77.76 | 2.45 | 16.99 | 2.28 | 0.53 |
| **ERR1347689^** | 44.0 | LP6005443-DNA_B03 | Eskimo Sireniki | CentralAsiaSiberia | 78.98 | 1.58 | 17.34 | 1.75 | 0.35 |
| **ERR1347741** | 49.4 | LP6005443-DNA_H03 | Eskimo Sireniki | CentralAsiaSiberia | 75.31 | 3.15 | 19.26 | 1.93 | 0.35 |
| **ERR1347690^** | 50.8 | LP6005443-DNA_B04 | Even | CentralAsiaSiberia | 90.37 | 1.75 | 4.38 | 2.98 | 0.53 |
| **ERR1347699** | 40.7 | LP6005443-DNA_C04 | Even | CentralAsiaSiberia | 91.77 | 2.28 | 3.33 | 1.75 | 0.88 |
| **ERR1395577** | 50.3 | LP6005592-DNA_F03 | Even | CentralAsiaSiberia | 75.66 | 6.30 | 8.76 | 3.68 | 5.60 |
| **ERR1347707^** | 48.3 | LP6005443-DNA_D04 | Itelman | CentralAsiaSiberia | 84.76 | 3.68 | 8.06 | 2.63 | 0.88 |
| **ERR1395583^** | 39.6 | LP6005677-DNA_A02 | Kyrgyz | CentralAsiaSiberia | 68.48 | 11.38 | 9.98 | 2.98 | 7.18 |
| **ERR1395586** | 48.9 | LP6005677-DNA_B02 | Kyrgyz | CentralAsiaSiberia | 69.18 | 11.91 | 8.41 | 3.15 | 7.36 |
| **ERR1347725^** | 39.3 | LP6005443-DNA_F04 | Mansi | CentralAsiaSiberia | 47.64 | 14.54 | 18.74 | 5.25 | 13.84 |
| **ERR1347734** | 46.7 | LP6005443-DNA_G04 | Mansi | CentralAsiaSiberia | 38.00 | 13.84 | 21.72 | 4.73 | 21.72 |
| **ERR1025645^** | 45.1 | LP6005441-DNA_F08 | Mongola | CentralAsiaSiberia | 93.35 | 2.45 | 2.28 | 1.75 | 0.18 |
| **ERR1419137** | 39.5 | LP6005441-DNA_E08 | Mongola | CentralAsiaSiberia | 93.52 | 1.93 | 2.28 | 1.40 | 0.88 |
| **ERR1347708^** | 48.5 | LP6005443-DNA_D05 | Tlingit | CentralAsiaSiberia | 34.33 | 8.06 | 34.50 | 3.50 | 19.61 |
| **ERR1347715** | 45.8 | LP6005443-DNA_E05 | Tlingit | CentralAsiaSiberia | 17.34 | 12.26 | 36.60 | 5.60 | 28.20 |
| **ERR1347657^** | 42.1 | LP6005442-DNA_E12 | Tubalar | CentralAsiaSiberia | 60.25 | 12.78 | 12.08 | 4.20 | 10.68 |
| **ERR1347663** | 39.1 | LP6005442-DNA_F12 | Tubalar | CentralAsiaSiberia | 56.04 | 14.54 | 13.84 | 4.90 | 10.68 |
| **ERR1347672^** | 40.8 | LP6005442-DNA_G12 | Ulchi | CentralAsiaSiberia | 93.87 | 1.75 | 2.28 | 1.75 | 0.35 |
| **ERR1347679** | 43.6 | LP6005442-DNA_H12 | Ulchi | CentralAsiaSiberia | 93.70 | 1.40 | 2.63 | 2.10 | 0.18 |
| **ERR1347658^** | 42.1 | LP6005442-DNA_F01 | Yakut | CentralAsiaSiberia | 85.29 | 5.08 | 5.25 | 2.45 | 1.93 |
| **ERR1347705** | 48.2 | LP6005443-DNA_D02 | Yakut | CentralAsiaSiberia | 87.04 | 4.38 | 4.38 | 1.93 | 2.28 |
| **ERR1347735^** | 47.0 | LP6005443-DNA_G05 | Ami | EastAsia | 97.72 | 1.05 | 0.70 | 0.53 | 0.00 |
| **ERR1419188** | 36.9 | LP6005442-DNA_C07 | Ami | EastAsia | 97.37 | 0.88 | 0.53 | 1.05 | 0.18 |
| **ERR1419198^** | 37.3 | LP6005442-DNA_E07 | Atayal | EastAsia | 97.37 | 0.35 | 0.70 | 1.58 | 0.00 |
| **ERR1395610^** | 55.0 | LP6005519-DNA_A06 | Burmese | EastAsia | 87.22 | 9.46 | 0.88 | 1.93 | 0.53 |
| **ERR1395613** | 60.0 | LP6005519-DNA_B06 | Burmese | EastAsia | 93.17 | 2.80 | 2.10 | 1.58 | 0.35 |
| **ERR1025617^** | 43.0 | LP6005441-DNA_G03 | Cambodian | EastAsia | 91.24 | 4.03 | 1.58 | 2.80 | 0.35 |
| **ERR1419162** | 39.1 | LP6005441-DNA_H03 | Cambodian | EastAsia | 89.49 | 5.78 | 1.05 | 2.98 | 0.70 |

| | | | | | | | | | |
|---|---|---|---|---|---|---|---|---|---|
| **ERR1347687^** | 40.7 | LP6005443-DNA_B01 | Dai | EastAsia | 97.72 | 0.53 | 0.70 | 0.70 | 0.35 |
| **ERR1395573** | 50.4 | LP6005592-DNA_D03 | Dai | EastAsia | 97.02 | 0.88 | 0.88 | 0.88 | 0.35 |
| **ERR1395618** | 41.3 | SS6004467 | Dai | EastAsia | 97.20 | 0.88 | 1.23 | 0.70 | 0.00 |
| **ERR1419123** | 39.3 | LP6005441-DNA_D04 | Dai | EastAsia | 96.67 | 1.40 | 1.05 | 0.53 | 0.35 |
| **ERR1425294^** | 46.9 | LP6005441-DNA_F04 | Daur | EastAsia | 93.52 | 2.28 | 1.75 | 1.58 | 0.88 |
| **ERR1019056^** | 43.0 | LP6005441-DNA_C05 | Han | EastAsia | 97.90 | 0.88 | 0.88 | 0.18 | 0.18 |
| **ERR1395592** | 39.9 | SS6004469 | Han | EastAsia | 97.20 | 0.18 | 0.88 | 1.40 | 0.35 |
| **ERR1419124** | 70.5 | LP6005441-DNA_D05 | Han | EastAsia | 97.20 | 1.05 | 1.40 | 0.35 | 0.00 |
| **ERR1025628^** | 41.1 | LP6005441-DNA_H05 | Hezhen | EastAsia | 94.57 | 1.23 | 2.28 | 1.40 | 0.53 |
| **ERR1419153** | 43.4 | LP6005441-DNA_G05 | Hezhen | EastAsia | 94.92 | 1.58 | 1.58 | 1.40 | 0.53 |
| **ERR1019038^** | 87.0 | LP6005441-DNA_C06 | Japanese | EastAsia | 98.07 | 0.70 | 0.53 | 0.53 | 0.18 |
| **ERR1395570** | 48.7 | LP6005592-DNA_C02 | Japanese | EastAsia | 97.20 | 1.05 | 0.88 | 0.53 | 0.35 |
| **ERR1419125** | 43.5 | LP6005441-DNA_D06 | Japanese | EastAsia | 98.07 | 0.35 | 1.05 | 0.53 | 0.00 |
| **ERR1025638^** | 37.3 | LP6005442-DNA_C11 | Kinh | EastAsia | 96.15 | 0.88 | 1.40 | 1.23 | 0.35 |
| **ERR1419196** | 38.3 | LP6005442-DNA_D11 | Kinh | EastAsia | 97.72 | 0.70 | 0.70 | 0.70 | 0.18 |
| **ERR1347700^** | 53.8 | LP6005443-DNA_C06 | Korean | EastAsia | 97.20 | 1.05 | 0.88 | 0.88 | 0.00 |
| **ERR1347709** | 44.7 | LP6005443-DNA_D06 | Korean | EastAsia | 96.32 | 0.70 | 1.23 | 1.40 | 0.35 |
| **ERR1019057^** | 38.4 | LP6005441-DNA_B07 | Lahu | EastAsia | 97.55 | 1.05 | 0.70 | 0.70 | 0.00 |
| **ERR1347713** | 59.4 | LP6005443-DNA_E01 | Lahu | EastAsia | 95.80 | 1.93 | 0.88 | 1.40 | 0.00 |
| **ERR1019060^** | 35.4 | LP6005441-DNA_C08 | Miao | EastAsia | 97.37 | 0.53 | 1.23 | 0.88 | 0.00 |
| **ERR1419127** | 37.7 | LP6005441-DNA_D08 | Miao | EastAsia | 97.02 | 0.88 | 0.53 | 1.23 | 0.35 |
| **ERR1019039^** | 41.5 | LP6005441-DNA_A09 | Naxi | EastAsia | 97.20 | 0.70 | 0.88 | 1.05 | 0.18 |
| **ERR1019079** | 43.5 | LP6005441-DNA_B09 | Naxi | EastAsia | 96.32 | 0.70 | 1.05 | 1.58 | 0.35 |
| **ERR1347719** | 42.2 | LP6005443-DNA_E09 | Naxi | EastAsia | 97.37 | 0.70 | 0.88 | 0.70 | 0.35 |
| **ERR1025647^** | 39.8 | LP6005441-DNA_F09 | Oroqen | EastAsia | 93.70 | 1.75 | 1.93 | 1.40 | 1.23 |
| **ERR1419138** | 39.5 | LP6005441-DNA_E09 | Oroqen | EastAsia | 94.92 | 0.88 | 1.93 | 2.10 | 0.18 |
| **ERR1347722^** | 49.1 | LP6005443-DNA_F01 | She | EastAsia | 96.67 | 0.35 | 1.05 | 1.58 | 0.35 |
| **ERR1347731** | 49.7 | LP6005443-DNA_G01 | She | EastAsia | 97.72 | 0.53 | 0.53 | 1.05 | 0.18 |
| **ERR1347684^** | 51.9 | LP6005443-DNA_A07 | Thai | EastAsia | 87.39 | 7.71 | 1.75 | 2.10 | 1.05 |
| **ERR1347692** | 46.5 | LP6005443-DNA_B07 | Thai | EastAsia | 95.62 | 2.10 | 1.23 | 1.05 | 0.00 |
| **ERR1347739^** | 50.8 | LP6005443-DNA_H01 | Tu | EastAsia | 90.37 | 3.85 | 1.58 | 2.80 | 1.40 |
| **ERR1419131** | 39.2 | LP6005441-DNA_D12 | Tu | EastAsia | 93.87 | 2.10 | 1.75 | 1.75 | 0.53 |
| **ERR1025652^** | 40.9 | LP6005441-DNA_F12 | Tujia | EastAsia | 97.20 | 0.70 | 1.23 | 0.70 | 0.18 |
| **ERR1347681** | 50.2 | LP6005443-DNA_A02 | Tujia | EastAsia | 97.90 | 0.35 | 0.70 | 0.70 | 0.35 |
| **ERR1025655^** | 40.0 | LP6005442-DNA_B01 | Uygur | EastAsia | 54.12 | 18.21 | 10.86 | 4.03 | 12.78 |

| | | | | | | | | | |
|---|---|---|---|---|---|---|---|---|---|
| **ERR1347688** | 51.1 | LP6005443-DNA_B02 | Uygur | EastAsia | 47.46 | 23.64 | 9.63 | 5.08 | 14.19 |
| **ERR1025656^** | 38.0 | LP6005442-DNA_D01 | Xibo | EastAsia | 92.82 | 2.98 | 1.58 | 1.93 | 0.70 |
| **ERR1347697** | 52.6 | LP6005443-DNA_C02 | Xibo | EastAsia | 94.75 | 1.23 | 2.28 | 1.05 | 0.70 |
| **ERR1347664^** | 44.2 | LP6005442-DNA_G01 | Yi | EastAsia | 96.85 | 0.53 | 1.58 | 1.05 | 0.00 |
| **ERR1347673** | 42.1 | LP6005442-DNA_H01 | Yi | EastAsia | 95.97 | 0.70 | 1.05 | 1.58 | 0.70 |
| **ERR1395600^** | 44.4 | SS6004477 | Australian | Oceania | 55.52 | 26.97 | 4.73 | 11.38 | 1.40 |
| **ERR1395601** | 46.2 | SS6004478 | Australian | Oceania | 54.12 | 27.15 | 4.38 | 12.78 | 1.58 |
| **ERR1019048^** | 42.2 | LP6005441-DNA_B03 | Bougainville | Oceania | 64.80 | 18.39 | 2.80 | 11.56 | 2.45 |
| **ERR1019049** | 38.5 | LP6005441-DNA_A03 | Bougainville | Oceania | 67.08 | 16.99 | 4.03 | 9.98 | 1.93 |
| **ERR1395554^** | 51.0 | LP6005519-DNA_E06 | Dusun | Oceania | 97.02 | 0.88 | 1.05 | 0.88 | 0.18 |
| **ERR1395557** | 53.0 | LP6005519-DNA_F06 | Dusun | Oceania | 96.67 | 1.05 | 0.53 | 1.40 | 0.35 |
| **ERR1395580^** | 46.8 | LP6005592-DNA_H03 | Hawaiian | Oceania | 92.12 | 1.23 | 0.88 | 5.25 | 0.53 |
| **ERR1395546^** | 40.6 | LP6005519-DNA_C06 | Igorot | Oceania | 97.02 | 1.23 | 0.53 | 0.88 | 0.35 |
| **ERR1395551** | 48.9 | LP6005519-DNA_D06 | Igorot | Oceania | 96.15 | 0.88 | 0.53 | 2.28 | 0.18 |
| **ERR1395566^** | 40.6 | LP6005592-DNA_B02 | Maori | Oceania | 42.73 | 15.24 | 15.06 | 10.51 | 16.46 |
| **ERR1019080^** | 46.4 | LP6005441-DNA_B10 | Papuan | Oceania | 56.92 | 24.87 | 3.68 | 11.91 | 2.63 |
| **ERR1347685** | 46.0 | LP6005443-DNA_A08 | Papuan | Oceania | 52.19 | 26.27 | 5.25 | 13.66 | 2.63 |
| **ERR1347693** | 42.7 | LP6005443-DNA_B08 | Papuan | Oceania | 53.24 | 26.80 | 4.38 | 12.78 | 2.80 |
| **ERR1347701** | 43.3 | LP6005443-DNA_C07 | Papuan | Oceania | 55.69 | 26.80 | 3.68 | 11.91 | 1.93 |
| **ERR1347702** | 45.0 | LP6005443-DNA_C08 | Papuan | Oceania | 54.99 | 26.44 | 3.85 | 12.61 | 2.10 |
| **ERR1347710** | 52.3 | LP6005443-DNA_D07 | Papuan | Oceania | 53.94 | 26.27 | 4.03 | 14.01 | 1.75 |
| **ERR1347711** | 44.1 | LP6005443-DNA_D08 | Papuan | Oceania | 53.77 | 26.44 | 5.25 | 12.78 | 1.75 |
| **ERR1347717** | 49.9 | LP6005443-DNA_E07 | Papuan | Oceania | 53.77 | 24.34 | 4.55 | 15.41 | 1.93 |
| **ERR1347718** | 35.9 | LP6005443-DNA_E08 | Papuan | Oceania | 52.54 | 26.97 | 4.38 | 13.84 | 2.28 |
| **ERR1347727** | 49.8 | LP6005443-DNA_F07 | Papuan | Oceania | 53.42 | 27.50 | 4.55 | 13.31 | 1.23 |
| **ERR1347728** | 44.1 | LP6005443-DNA_F08 | Papuan | Oceania | 54.47 | 27.15 | 3.15 | 12.61 | 2.63 |
| **ERR1347736** | 45.4 | LP6005443-DNA_G07 | Papuan | Oceania | 55.17 | 25.92 | 3.85 | 13.31 | 1.75 |
| **ERR1395595** | 47.3 | SS6004472 | Papuan | Oceania | 50.61 | 28.20 | 3.68 | 15.06 | 2.45 |
| **ERR1395605** | 48.0 | LP6005443-DNA_H07 | Papuan | Oceania | 54.29 | 26.97 | 3.50 | 13.66 | 1.58 |
| **ERR1419095** | 35.1 | LP6005441-DNA_A10 | Papuan | Oceania | 56.57 | 23.82 | 4.73 | 12.26 | 2.63 |
| **ERR1019035^** | 43.0 | LP6005441-DNA_D01 | Balochi | SouthAsia | 3.50 | 62.87 | 7.36 | 11.21 | 15.06 |
| **ERR1419110** | 43.3 | LP6005441-DNA_C01 | Balochi | SouthAsia | 3.68 | 63.22 | 7.88 | 7.36 | 17.86 |
| **ERR1347669^** | 44.5 | LP6005442-DNA_G09 | Bengali | SouthAsia | 15.41 | 73.56 | 3.50 | 3.85 | 3.68 |
| **ERR1347676** | 34.0 | LP6005442-DNA_H09 | Bengali | SouthAsia | 14.36 | 73.91 | 4.03 | 4.38 | 3.33 |
| **ERR1395559^** | 52.9 | LP6005519-DNA_G03 | Brahmin | SouthAsia | 8.93 | 75.31 | 2.80 | 5.78 | 7.18 |

| | | | | | | | | | |
|---|---|---|---|---|---|---|---|---|---|
| **ERR1395561** | 36.6 | LP6005519-DNA_H03 | Brahmin | SouthAsia | 7.53 | 76.01 | 4.03 | 7.36 | 5.08 |
| **ERR1419112^** | 35.7 | LP6005441-DNA_C03 | Brahui | SouthAsia | 6.65 | 54.29 | 11.03 | 8.06 | 19.96 |
| **ERR1419122** | 38.3 | LP6005441-DNA_D03 | Brahui | SouthAsia | 4.20 | 55.87 | 9.28 | 10.51 | 20.14 |
| **ERR1025616^** | 42.3 | LP6005441-DNA_E03 | Burusho | SouthAsia | 12.96 | 60.07 | 7.53 | 5.95 | 13.49 |
| **ERR1419142** | 41.6 | LP6005441-DNA_F03 | Burusho | SouthAsia | 9.98 | 60.77 | 7.36 | 6.65 | 15.24 |
| **ERR1025625^** | 64.8 | LP6005441-DNA_F05 | Hazara | SouthAsia | 49.39 | 22.59 | 10.16 | 6.13 | 11.73 |
| **ERR1419134** | 67.7 | LP6005441-DNA_E05 | Hazara | SouthAsia | 50.96 | 23.82 | 9.98 | 5.60 | 9.63 |
| **ERR1395545^** | 50.2 | LP6005519-DNA_C05 | Irula | SouthAsia | 13.13 | 77.58 | 1.93 | 5.60 | 1.75 |
| **ERR1395550** | 53.1 | LP6005519-DNA_D05 | Irula | SouthAsia | 12.78 | 76.71 | 2.63 | 5.25 | 2.63 |
| **ERR1025633^** | 37.3 | LP6005441-DNA_E06 | Kalash | SouthAsia | 4.90 | 60.60 | 11.03 | 5.25 | 18.21 |
| **ERR1419144** | 34.9 | LP6005441-DNA_F06 | Kalash | SouthAsia | 6.30 | 57.27 | 10.16 | 8.06 | 18.21 |
| **ERR1395608^** | 52.1 | LP6005519-DNA_A04 | Kapu | SouthAsia | 10.33 | 79.33 | 2.45 | 4.20 | 3.68 |
| **ERR1395611** | 52.8 | LP6005519-DNA_B04 | Kapu | SouthAsia | 16.11 | 74.96 | 2.98 | 4.03 | 1.93 |
| **ERR1395553^** | 50.3 | LP6005519-DNA_E05 | Khonda Dora | SouthAsia | 50.09 | 43.78 | 1.23 | 4.55 | 0.35 |
| **ERR1347703^** | 45.4 | LP6005443-DNA_C09 | Kusunda | SouthAsia | 71.80 | 20.84 | 2.98 | 3.68 | 0.70 |
| **ERR1347712** | 44.4 | LP6005443-DNA_D09 | Kusunda | SouthAsia | 75.83 | 17.86 | 2.10 | 3.50 | 0.70 |
| **ERR1395560^** | 47.5 | LP6005519-DNA_G04 | Madiga | SouthAsia | 10.16 | 80.21 | 2.80 | 4.03 | 2.80 |
| **ERR1395562** | 49.5 | LP6005519-DNA_H04 | Madiga | SouthAsia | 11.56 | 79.33 | 1.75 | 5.25 | 2.10 |
| **ERR1019059^** | 40.3 | LP6005441-DNA_D07 | Makrani | SouthAsia | 2.80 | 52.19 | 9.98 | 14.01 | 21.02 |
| **ERR1419115** | 43.2 | LP6005441-DNA_C07 | Makrani | SouthAsia | 1.93 | 56.74 | 11.38 | 8.93 | 21.02 |
| **ERR1395552^** | 49.6 | LP6005519-DNA_E04 | Mala | SouthAsia | 11.03 | 80.39 | 1.58 | 4.55 | 2.45 |
| **ERR1395556** | 48.4 | LP6005519-DNA_F04 | Mala | SouthAsia | 11.03 | 81.26 | 2.10 | 3.85 | 1.75 |
| **ERR1019065^** | 42.4 | LP6005441-DNA_D10 | Pathan | SouthAsia | 4.73 | 70.23 | 6.65 | 4.90 | 13.49 |
| **ERR1419118** | 41.4 | LP6005441-DNA_C10 | Pathan | SouthAsia | 3.68 | 65.15 | 7.53 | 6.30 | 17.34 |
| **ERR1025664^** | 34.0 | LP6005442-DNA_B12 | Punjabi | SouthAsia | 9.98 | 80.21 | 2.45 | 3.68 | 3.68 |
| **ERR1395564** | 35.3 | LP6005592-DNA_A04 | Punjabi | SouthAsia | 8.41 | 67.60 | 6.48 | 5.08 | 12.43 |
| **ERR1395568** | 35.0 | LP6005592-DNA_B04 | Punjabi | SouthAsia | 8.93 | 80.04 | 3.68 | 4.20 | 3.15 |
| **ERR1419177** | 39.0 | LP6005442-DNA_A12 | Punjabi | SouthAsia | 8.58 | 81.09 | 2.28 | 4.38 | 3.68 |
| **ERR1395609^** | 46.8 | LP6005519-DNA_A05 | Relli | SouthAsia | 19.79 | 71.98 | 2.28 | 4.38 | 1.58 |
| **ERR1395612** | 49.4 | LP6005519-DNA_B05 | Relli | SouthAsia | 10.86 | 77.58 | 3.50 | 4.73 | 3.33 |
| **ERR1025648^** | 40.0 | LP6005441-DNA_G11 | Sindhi | SouthAsia | 5.25 | 67.78 | 7.88 | 5.78 | 13.31 |
| **ERR1025649** | 39.4 | LP6005441-DNA_H11 | Sindhi | SouthAsia | 4.55 | 74.26 | 5.60 | 7.18 | 8.41 |
| **ERR1395549^** | 50.2 | LP6005519-DNA_D04 | Yadava | SouthAsia | 8.76 | 81.26 | 2.80 | 4.90 | 2.28 |
| **ERR1395615** | 55.5 | LP6005519-DNA_C04 | Yadava | SouthAsia | 9.98 | 80.21 | 1.75 | 5.08 | 2.98 |

| | | | | | | | | | |
|---|---|---|---|---|---|---|---|---|---|
| **ERR1025608^** | 37.9 | LP6005442-DNA_C02 | Abkhasian | WestEurasia | 5.08 | 28.20 | 13.49 | 6.48 | 46.76 |
| **ERR1419191** | 39.6 | LP6005442-DNA_D02 | Abkhasian | WestEurasia | 4.20 | 27.85 | 12.08 | 7.71 | 48.16 |
| **ERR1419088^** | 41.7 | LP6005441-DNA_A01 | Adygei | WestEurasia | 2.63 | 26.44 | 13.66 | 6.30 | 50.96 |
| **ERR1419098** | 35.4 | LP6005441-DNA_B01 | Adygei | WestEurasia | 4.38 | 24.87 | 12.96 | 7.88 | 49.91 |
| **ERR1395585^** | 38.2 | LP6005677-DNA_B01 | Albanian | WestEurasia | 1.40 | 14.19 | 15.06 | 7.53 | 61.82 |
| **ERR1347665^** | 40.4 | LP6005442-DNA_G02 | Armenian | WestEurasia | 1.75 | 24.69 | 16.64 | 7.18 | 49.74 |
| **ERR1395555** | 46.2 | LP6005519-DNA_F03 | Armenian | WestEurasia | 1.40 | 26.80 | 16.46 | 8.58 | 46.76 |
| **ERR1019045^** | 42.5 | LP6005441-DNA_C02 | Basque | WestEurasia | 1.58 | 8.58 | 15.41 | 5.08 | 69.35 |
| **ERR1019046** | 38.8 | LP6005441-DNA_D02 | Basque | WestEurasia | 1.23 | 7.71 | 17.86 | 6.30 | 66.90 |
| **ERR1019047^** | 42.3 | LP6005441-DNA_F02 | BedouinB | WestEurasia | 2.28 | 15.24 | 23.12 | 19.09 | 40.28 |
| **ERR1419132** | 44.7 | LP6005441-DNA_E02 | BedouinB | WestEurasia | 2.45 | 16.99 | 21.37 | 18.21 | 40.98 |
| **ERR1019036^** | 80.3 | LP6005441-DNA_B06 | Bergamo | WestEurasia | 2.45 | 12.43 | 15.59 | 7.01 | 62.52 |
| **ERR1419092** | 70.9 | LP6005441-DNA_A06 | Bergamo | WestEurasia | 1.75 | 10.68 | 15.06 | 8.58 | 63.92 |
| **ERR1025614^** | 42.4 | LP6005442-DNA_A03 | Bulgarian | WestEurasia | 1.93 | 13.84 | 14.36 | 6.83 | 63.05 |
| **ERR1025615** | 37.9 | LP6005442-DNA_B03 | Bulgarian | WestEurasia | 2.10 | 12.26 | 14.71 | 5.60 | 65.32 |
| **ERR1025599^** | 46.6 | LP6005442-DNA_D03 | Chechen | WestEurasia | 3.15 | 26.44 | 16.81 | 7.36 | 46.23 |
| **ERR1395617^** | 44.1 | LP6007068-DNA_A01 | Crete | WestEurasia | 2.63 | 17.16 | 15.59 | 9.63 | 54.99 |
| **ERR1425293** | 43.7 | LP6007069-DNA_A01 | Crete | WestEurasia | 2.63 | 15.59 | 15.94 | 8.93 | 56.92 |
| **ERR1395604^** | 50.9 | LP6005443-DNA_H05 | Czech | WestEurasia | 1.75 | 10.33 | 17.34 | 6.83 | 63.75 |
| **ERR1347704^** | 51.2 | LP6005443-DNA_D01 | Druze | WestEurasia | 1.23 | 20.32 | 14.71 | 15.41 | 48.34 |
| **ERR1419152** | 45.0 | LP6005441-DNA_G04 | Druze | WestEurasia | 2.10 | 21.54 | 17.51 | 12.08 | 46.76 |
| **ERR1347661^** | 46.0 | LP6005442-DNA_F10 | English | WestEurasia | 1.75 | 8.76 | 13.66 | 5.08 | 70.75 |
| **ERR1419199** | 41.7 | LP6005442-DNA_E10 | English | WestEurasia | 1.93 | 8.58 | 13.31 | 6.48 | 69.70 |
| **ERR1347666^** | 35.6 | LP6005442-DNA_G03 | Estonian | WestEurasia | 2.80 | 8.76 | 14.89 | 5.60 | 67.95 |
| **ERR1347674** | 38.6 | LP6005442-DNA_H03 | Estonian | WestEurasia | 3.33 | 10.86 | 13.84 | 5.43 | 66.55 |
| **ERR1025659^** | 38.2 | LP6005442-DNA_D10 | Finnish | WestEurasia | 4.90 | 7.88 | 15.59 | 4.38 | 67.25 |
| **ERR1346534** | 35.2 | LP6005442-DNA_C10 | Finnish | WestEurasia | 3.50 | 10.86 | 12.26 | 6.13 | 67.25 |
| **ERR1395563** | 47.9 | LP6005592-DNA_A02 | Finnish | WestEurasia | 3.68 | 11.56 | 12.96 | 5.43 | 66.37 |
| **ERR1019053^** | 44.0 | LP6005441-DNA_A05 | French | WestEurasia | 1.58 | 9.98 | 16.46 | 6.30 | 65.67 |
| **ERR1019054** | 39.9 | LP6005441-DNA_B05 | French | WestEurasia | 0.88 | 9.98 | 17.51 | 5.60 | 66.02 |
| **ERR1395619** | 45.9 | SS6004468 | French | WestEurasia | 1.23 | 10.16 | 16.11 | 5.78 | 66.73 |
| **ERR1025624^** | 33.9 | LP6005442-DNA_A04 | Georgian | WestEurasia | 1.93 | 27.15 | 13.49 | 7.71 | 49.74 |
| **ERR1419181** | 43.0 | LP6005442-DNA_B04 | Georgian | WestEurasia | 1.93 | 26.44 | 15.06 | 8.41 | 48.16 |
| **ERR1347668^** | 37.9 | LP6005442-DNA_G07 | Greek | WestEurasia | 1.93 | 14.19 | 13.66 | 7.88 | 62.35 |

| | | | | | | | | | |
|---|---|---|---|---|---|---|---|---|---|
| **ERR1347683** | 46.4 | LP6005443-DNA_A06 | Greek | WestEurasia | 1.40 | 26.09 | 12.43 | 6.65 | 53.42 |
| **ERR1025630^** | 41.0 | LP6005442-DNA_A08 | Hungarian | WestEurasia | 2.10 | 11.21 | 14.19 | 6.13 | 66.37 |
| **ERR1419182** | 40.1 | LP6005442-DNA_B08 | Hungarian | WestEurasia | 3.15 | 11.56 | 15.41 | 5.43 | 64.45 |
| **ERR1025631^** | 38.7 | LP6005442-DNA_D08 | Icelandic | WestEurasia | 2.28 | 11.38 | 14.36 | 4.73 | 67.25 |
| **ERR1347691** | 49.9 | LP6005443-DNA_B06 | Icelandic | WestEurasia | 1.93 | 9.63 | 14.36 | 5.60 | 68.48 |
| **ERR1025632^** | 38.0 | LP6005442-DNA_C04 | Iranian | WestEurasia | 3.50 | 34.15 | 13.66 | 11.56 | 37.13 |
| **ERR1347695** | 46.6 | LP6005443-DNA_B10 | Iranian | WestEurasia | 3.15 | 35.38 | 13.49 | 9.98 | 38.00 |
| **ERR1395576^** | 43.1 | LP6005592-DNA_F01 | Iraqi Jew | WestEurasia | 1.40 | 24.34 | 17.86 | 9.46 | 46.94 |
| **ERR1395620** | 48.4 | LP6005592-DNA_E01 | Iraqi Jew | WestEurasia | 1.40 | 22.77 | 15.06 | 12.26 | 48.51 |
| **ERR1025661^** | 39.7 | LP6005442-DNA_E04 | Jordanian | WestEurasia | 1.93 | 18.91 | 18.74 | 19.61 | 40.81 |
| **ERR1347659** | 36.6 | LP6005442-DNA_F04 | Jordanian | WestEurasia | 2.10 | 17.16 | 23.12 | 18.74 | 38.88 |
| **ERR1395578** | 48.4 | LP6005592-DNA_G03 | Jordanian | WestEurasia | 1.93 | 18.74 | 20.49 | 18.21 | 40.63 |
| **ERR1347667^** | 40.2 | LP6005442-DNA_G04 | Lezgin | WestEurasia | 1.93 | 26.97 | 13.66 | 8.06 | 49.39 |
| **ERR1347675** | 40.4 | LP6005442-DNA_H04 | Lezgin | WestEurasia | 1.75 | 28.20 | 15.24 | 7.18 | 47.64 |
| **ERR1347720^** | 44.0 | LP6005443-DNA_E10 | North Ossetian | WestEurasia | 6.83 | 27.85 | 16.64 | 7.53 | 41.16 |
| **ERR1347729** | 47.3 | LP6005443-DNA_F10 | North Ossetian | WestEurasia | 5.60 | 26.27 | 15.06 | 6.83 | 46.23 |
| **ERR1395565^** | 50.1 | LP6005592-DNA_B01 | Norwegian | WestEurasia | 2.10 | 8.41 | 15.59 | 6.13 | 67.78 |
| **ERR1019062^** | 39.8 | LP6005441-DNA_C09 | Orcadian | WestEurasia | 1.58 | 9.81 | 16.99 | 6.65 | 64.97 |
| **ERR1419128** | 39.8 | LP6005441-DNA_D09 | Orcadian | WestEurasia | 0.88 | 10.68 | 14.36 | 5.78 | 68.30 |
| **ERR1025662^** | 42.1 | LP6005441-DNA_G09 | Palestinian | WestEurasia | 1.93 | 19.44 | 20.84 | 17.69 | 40.11 |
| **ERR1395567** | 46.7 | LP6005592-DNA_B03 | Palestinian | WestEurasia | 1.58 | 17.34 | 20.32 | 21.72 | 39.05 |
| **ERR1419167** | 41.2 | LP6005441-DNA_H09 | Palestinian | WestEurasia | 1.05 | 15.59 | 21.72 | 18.91 | 42.73 |
| **ERR1395575^** | 50.3 | LP6005592-DNA_E02 | Polish | WestEurasia | 2.45 | 12.43 | 13.49 | 7.36 | 64.27 |
| **ERR1025605^** | 39.0 | LP6005441-DNA_G10 | Russian | WestEurasia | 6.48 | 14.19 | 16.99 | 5.95 | 56.39 |
| **ERR1025606** | 39.1 | LP6005441-DNA_H10 | Russian | WestEurasia | 5.95 | 13.13 | 16.11 | 4.73 | 60.07 |
| **ERR1395569^** | 50.2 | LP6005592-DNA_C01 | Saami | WestEurasia | 14.36 | 12.78 | 20.14 | 4.55 | 48.16 |
| **ERR1395616** | 45.3 | LP6005592-DNA_D01 | Saami | WestEurasia | 19.26 | 11.38 | 19.79 | 5.95 | 43.61 |
| **ERR1395574^** | 43.0 | LP6005592-DNA_D04 | Samaritan | WestEurasia | 1.05 | 19.44 | 19.61 | 13.49 | 46.41 |
| **ERR1019068^** | 37.2 | LP6005441-DNA_C11 | Sardinian | WestEurasia | 0.53 | 8.06 | 14.89 | 8.23 | 68.30 |
| **ERR1395597** | 41.7 | SS6004474 | Sardinian | WestEurasia | 1.40 | 8.23 | 15.94 | 8.93 | 65.50 |
| **ERR1419130** | 34.8 | LP6005441-DNA_D11 | Sardinian | WestEurasia | 0.70 | 7.01 | 19.44 | 7.88 | 64.97 |
| **ERR1419176^** | 35.5 | LP6005442-DNA_A11 | Spanish | WestEurasia | 1.40 | 8.76 | 16.29 | 8.58 | 64.97 |
| **ERR1419184** | 43.0 | LP6005442-DNA_B11 | Spanish | WestEurasia | 0.53 | 9.28 | 16.64 | 9.46 | 64.10 |
| **ERR1395548^** | 48.1 | LP6005519-DNA_D03 | Tajik | WestEurasia | 5.43 | 37.65 | 13.84 | 7.88 | 35.20 |

| | | | | | | | | | |
|---|---|---|---|---|---|---|---|---|---|
| **ERR1395614** | 54.2 | LP6005519-DNA_C03 | Tajik | WestEurasia | 7.36 | 41.16 | 13.13 | 5.25 | 33.10 |
| **ERR1395584^** | 44.6 | LP6005677-DNA_A03 | Turkish | WestEurasia | 6.30 | 25.57 | 18.04 | 7.53 | 42.56 |
| **ERR1395587** | 45.5 | LP6005677-DNA_C03 | Turkish | WestEurasia | 7.18 | 23.47 | 14.71 | 9.46 | 45.18 |
| **ERR1025653^** | 40.2 | LP6005441-DNA_H12 | Tuscan | WestEurasia | 1.75 | 10.68 | 14.01 | 8.58 | 64.97 |
| **ERR1419160** | 43.6 | LP6005441-DNA_G12 | Tuscan | WestEurasia | 1.93 | 12.43 | 16.29 | 8.23 | 61.12 |
| **ERR1395579^** | 51.7 | LP6005592-DNA_H01 | Yemenite Jew | WestEurasia | 2.28 | 17.16 | 21.02 | 17.69 | 41.86 |
| **ERR1395621** | 56.2 | LP6005592-DNA_G01 | Yemenite Jew | WestEurasia | 2.28 | 15.94 | 21.02 | 18.91 | 41.86 |

***We selected a unique download Accession for each record (Library) to avoid duplicates. ^Population representatives (129 individuals) are plotted in Figure 1d when multiple individuals from the same population are identified, to minimize crowding on the display. ‡Half the WGS was downloaded and processed for this individual of American ancestry and ntRoot predictions, consistent with the region of origin (population Pima from America), are achievable on WGS data as low as 22-fold coverage.**

**Table S7.** ntRoot super-population level GAI summary on complete and draft genome sequences, with matching raw whole-genome read sequencing data (where applicable). The super-populations are EAS: East Asian, SAS: South Asian, AMR: Admixed American, AFR: African, EUR: European.

| Individual | Type | Assigned Label | ntRoot GAI | SNV Count | Non-zero AF SNV Count | GAI score |
|---|---|---|---|---|---|---|
| **HuRef** | WGS | CEU(EUR) | EUR | 2,636,477 | 2,592,289 | 0.4942 |
| | WGA HuRef‡ | CEU(EUR) | EUR | 1,943,423 | 1,903,006 | 0.5361 |
| | WGA HuRefPrime‡ | CEU(EUR) | EUR | 1,945,709 | 1,905,081 | 0.5355 |
| **CN1** | WGA maternal | EAS | EAS | 2,002,956 | 1,960,790 | 0.5756 |
| | WGA paternal | EAS | EAS | 2,011,544 | 1,969,732 | 0.5754 |
| **HuRef/CN1 simulated diploid** | WGA | mixed | EAS | 2,776,922 | 2,640,999 | 0.4840 |
| **KOREF** | WGA* | EAS | EAS | 1,996,558 | 1,953,649 | 0.5785 |
| | WGS(Illumina) | EAS | EAS | 2,622,427 | 2,578,255 | 0.5314 |
| | WGS(PacBio SII) | EAS | EAS | 2,672,832 | 2,629,079 | 0.5292 |
| **NA01243** | WGS(Illumina) | AMR | AFR | 3,266,892 | 3,180,440 | 0.4190 |
| **NA24385** | WGA maternal | EUR | EUR | 2,098,033 | 1,978,829 | 0.4842 |
| | WGA paternal | EUR | EUR | 2,096,296 | 1,980,081 | 0.4882 |
| | WGS(Illumina) | EUR | EUR | 2,904,614 | 2,773,521 | 0.4528 |
| | WGS(ONT V14) | EUR | EUR | 2,859,212 | 2,729,585 | 0.4541 |
| | WGS(PacBio SII) | EUR | EUR | 2,886,074 | 2,756,569 | 0.4539 |
| **HG02055** | WGS(Illumina) | AFR | AFR | 3,187,354 | 3,161,469 | 0.4402 |
| | WGA(GoldRush*,^) | AFR | AFR | 1,168,009 | 1,144,993 | 0.5060 |
| | WGA(Shasta*,^) | AFR | AFR | 2,055,892 | 2,035,138 | 0.4968 |
| | WGA(Flye*,^) | AFR | AFR | 2,123,506 | 2,102,196 | 0.4859 |
| | WGA maternal | AFR | AFR | 2,249,827 | 2,226,829 | 0.4892 |
| | WGA paternal | AFR | AFR | 2,271,615 | 2,253,601 | 0.4910 |

***Pseudohaploid assembly, assembled with WGS Oxford Nanopore datasets (Wong et al., 2022, Table 6). The mat/pat suffix indicates maternal and paternal haplotypes, respectively. ‡HuRef and HuRefPrime haploid WGAs each refer to the principal and alternate haplotypes, respectively.**

**Table S8.** ntRoot (unless specified) super-population level LAI-based ancestry fraction estimates on complete and draft genome sequences, with matching raw whole-genome read sequencing data (where applicable). The LAI fraction estimates were generated using 5Mbp tiles. The super-populations are EAS: East Asian, SAS: South Asian, AMR: Admixed American, AFR: African, EUR: European.

| Individual | Type | Assigned Label | Predicted from LAI | LAI EAS fraction (%) | LAI SAS fraction (%) | LAI AMR fraction (%) | LAI AFR fraction (%) | LAI EUR fraction (%) |
|---|---|---|---|---|---|---|---|---|
| **HuRef** | WGS | CEU(EUR) | EUR | 1.75 | 11.38 | 14.01 | 7.01 | 65.85 |
| | WGA HuRef‡ | CEU(EUR) | EUR | 2.28 | 11.03 | 16.11 | 8.93 | 61.65 |
| | WGA HuRefPrime‡ | CEU(EUR) | EUR | 2.45 | 11.91 | 15.76 | 9.11 | 60.77 |
| **CN1** | WGA maternal | EAS | EAS | 95.27 | 0.88 | 0.70 | 2.10 | 1.05 |
| | WGA paternal | EAS | EAS | 95.27 | 0.88 | 2.10 | 1.58 | 0.18 |
| **HuRef/CN1 simulated diploid** | WGA | mixed | EAS | 54.63 | 11.29 | 12.57 | 7.77 | 13.74 |
| **KOREF** | WGA* | EAS | EAS | 94.57 | 0.70 | 1.23 | 2.98 | 0.53 |
| | WGS(Illumina) | EAS | EAS | 97.72 | 0.53 | 0.53 | 0.88 | 0.35 |
| | WGS(PacBio) | EAS | EAS | 97.90 | 0.35 | 0.53 | 0.88 | 0.35 |
| **NA24385** | WGA maternal | EUR | EUR | 4.73 | 14.36 | 21.89 | 18.91 | 40.11 |
| | WGA paternal | EUR | EUR | 3.33 | 15.59 | 23.29 | 17.69 | 40.11 |
| | WGS(Illumina) | EUR | EUR | 1.58 | 14.01 | 22.94 | 16.64 | 44.83 |
| | WGS(ONT V14) | EUR | EUR | 1.23 | 13.84 | 23.47 | 17.34 | 44.13 |
| | WGS(PacBio SII) | EUR | EUR | 1.40 | 14.19 | 22.94 | 16.99 | 44.48 |
| **NA01243** | WGS(Illumina) | AMR | | | | | | |
| ntRoot | | | AFR | 0.70 | 0.70 | 15.76 | 77.58 | 5.25 |
| ADMIXTURE | | | AFR | 0.00 | 0.00 | 29.12 | 70.87 | 0.00 |
| **HG02055** | WGS(Illumina) | AFR | AFR | 0.00 | 0.00 | 5.43 | 93.35 | 1.23 |
| | WGA(GoldRush*,^) | AFR | AFR | 0.18 | 1.05 | 4.90 | 93.17 | 0.70 |
| | WGA(Shasta*,^) | AFR | AFR | 0.18 | 0.35 | 5.78 | 92.47 | 1.23 |
| | WGA(Flye*,^) | AFR | AFR | 0.18 | 0.35 | 5.60 | 92.82 | 1.05 |
| | WGA maternal | AFR | AFR | 1.05 | 1.58 | 3.68 | 83.89 | 9.81 |
| | WGA paternal | AFR | AFR | 0.35 | 1.23 | 2.28 | 93.87 | 2.28 |

***Pseudohaploid assembly. ^Assembled with WGS Oxford Nanopore datasets (Wong et al., 2022, Table 6). The mat/pat suffix indicates maternal and paternal haplotypes, respectively. ‡HuRef and HuRefPrime haploid WGAs each refer to the principal and alternate haplotypes, respectively.**

**Table S9.** Resource usage for ntRoot, SNVstory and ADMIXTURE.

| Individual | Type | Input sequence file size# (GB) | Wall-clock run time (h) | Peak RAM (GB) |
|---|---|---|---|---|
| **HuRef** | WGS~47X | 75.0 | 1.30 | 82.05 |
| | WGA HuRef‡ | 2.7 | 0.47 | 12.77 |
| | WGA HuRefPrime‡ | 2.7 | 0.48 | 12.72 |
| **CN1** | WGA maternal | 2.9 | 0.53 | 12.99 |
| | WGA paternal | 2.8 | 0.55 | 12.79 |
| **HuRef/CN1 simulated diploid+** | WGA | 5.4 | 0.50 | 14.84 |
| **KOREF** | WGS~42X (Illumina) | 72.0 | 1.27 | 84.77 |
| | WGS~38X (PacBio SII) | 79.0 | 1.27 | 77.30 |
| | WGA* | 2.7 | 0.47 | 12.86 |
| **NA24385 (HG002)** | WGA maternal | 2.9 | 0.41 | 14.54 |
| | WGA paternal | 2.8 | 0.39 | 14.30 |
| | WGS~60X(Illumina) | 42.0 | 1.26 | 77.67 |
| | WGS~72X(ONT V14) | 177.0 | 2.15 | 148.19 |
| | WGS~34X(PacBio SII) | 39.0 | 1.19 | 75.90 |
| **HG02055** | WGS~38X | 26.0 | 0.98 | 66.48 |
| | WGA(GoldRush*,^) | 2.9 | 0.42 | 13.08 |
| | WGA(Shasta*,^) | 2.8 | 0.47 | 12.99 |
| | WGA(Flye*,^) | 2.8 | 0.47 | 12.90 |
| | WGA maternal | 2.9 | 0.48 | 12.99 |
| | WGA paternal | 2.9 | 0.49 | 12.82 |
| **Validation set, n=266 1kGP** | WGS~30X | ~26.6 | 1.14 +/- 0.11 | 67.29 +/- 1.68 |
| **Benchmark set, n=100 1kGP** | WGS~30X | ~27.2 | | |
| **ntRoot** | | | 1.14 +/- 0.18 | 67.08 +/- 1.73 |
| **SNVstory** | | | 4.42 +/- 0.21 | 87.35 +/- 6.71 |
| **ADMIXTURE** | | | 273.70 | 449.15 |
| **Discovery set, n=279 SGDP** | WGS~44X [min:22X-max:87X] | ~116.1 | 1.60 +/- 0.22 | 68.19 +/- 7.32 |

***Pseudohaploid assembly. ^Assembled with WGS Oxford Nanopore datasets (Wong et al., 2022, Table 6). The mat/pat suffix indicates maternal and paternal haplotypes, respectively. ‡HuRef and HuRefPrime haploid WGAs each refer to the principal and alternate haplotypes, respectively. #The WGS datasets input file size is gzip compressed. +Unlike with other WGA input, we used a 2 x 2.7 Gbp (~5.4 Gbp) genome input for this ntRoot analysis.**

**Table S10.** Effect of HG002 (NA24385, Illumina) WGS coverage on ntRoot ancestry predictions. The largest ancestry fraction is shown with bold face.

| Fold coverage | Read pairs sampled (M) | LAI EAS fraction (%) | LAI SAS fraction (%) | LAI AMR fraction (%) | LAI AFR fraction (%) | LAI EUR fraction (%) |
|---|---|---|---|---|---|---|
| 59.9 (full) | 179.9 | 1.58 | 14.01 | 22.94 | 16.64 | **44.83** |
| 55.0 | 165.0 | 1.40 | 14.01 | 23.12 | 17.16 | **44.31** |
| 50.0 | 150.0 | 1.58 | 14.01 | 23.29 | 17.16 | **43.96** |
| 45.0 | 135.0 | 1.40 | 14.01 | 23.12 | 17.34 | **44.13** |
| 40.0 | 120.0 | 1.23 | 13.84 | 23.29 | 17.69 | **43.96** |
| 35.0 | 105.0 | 1.23 | 14.54 | 23.12 | 17.51 | **43.61** |
| 30.0 | 90.0 | 1.23 | 13.66 | 23.82 | 18.21 | **43.08** |
| 25.0 | 75.0 | 1.05 | 13.84 | 23.12 | 18.56 | **43.43** |
| 20.0 | 60.0 | 1.05 | 14.36 | 23.64 | 18.21 | **42.73** |
| 17.5 | 52.5 | 1.23 | 14.71 | 23.29 | 18.21 | **42.56** |
| 15.0 | 45.0 | 1.23 | 14.54 | 23.47 | 18.39 | **42.38** |
| 12.5 | 37.5 | 1.23 | 14.36 | 23.82 | 18.39 | **42.21** |
| 10.0 | 30.0 | 0.70 | 0.88 | 0.70 | **97.55** | 0.18 |
| 7.5 | 22.5 | 0.35 | 1.05 | 1.40 | **97.19** | 0.00 |
| 5.0 | 15.0 | 0.35 | 1.05 | 1.05 | **97.36** | 0.18 |
| 2.5 | 7.5 | 0.70 | 1.23 | 1.23 | **96.66** | 0.18 |

**Table S11.** Rationale for selecting the parameter k in ntRoot. Performance (Sensitivity, Precision, F1) of ntRoot in identifying ancestry-discriminant SNVs against a benchmark set of known HG002 variants[12] as a function of parameter k (word length) and its effect on ancestry fraction predictions using the HG002 (NA24385, Illumina) WGS dataset. The largest ancestry fraction is shown in bold face.

| k | Sensitivity | Precision | F1 | LAI EAS fraction (%) | LAI SAS fraction (%) | LAI AMR fraction (%) | LAI AFR fraction (%) | LAI EUR fraction (%) |
|---|---|---|---|---|---|---|---|---|
| 70 | 0.7739 | 0.8651 | 0.8170 | 1.75 | 14.71 | 18.74 | 11.73 | **53.06** |
| 65 | 0.7830 | 0.8577 | 0.8186 | 1.75 | 15.24 | 18.39 | 12.43 | **52.19** |
| 60 | 0.7956 | 0.8447 | 0.8194 | 1.93 | 14.71 | 20.14 | 13.84 | **49.39** |
| 55 | 0.8051 | 0.8262 | 0.8155 | 1.58 | 14.01 | 22.94 | 16.64 | **44.83** |
| 50 | 0.8093 | 0.8089 | 0.8091 | 1.23 | 12.96 | 25.22 | 20.49 | **40.11** |
| 45 | 0.8189 | 0.7813 | 0.7997 | 1.05 | 12.08 | 28.02 | 23.64 | **35.20** |
| 40 | 0.8287 | 0.7354 | 0.7793 | 0.88 | 9.98 | **31.87** | 31.17 | 26.09 |
| 35 | 0.8299 | 0.6599 | 0.7352 | 0.70 | 9.28 | 31.52 | **37.48** | 21.02 |
| 30 | 0.8302 | 0.7034 | 0.7616 | 0.53 | 7.18 | 31.00 | **47.81** | 13.49 |

**Table S12.** ADMIXTURE (AdM), SNVstory (SnvS) and ntRoot (ntR) ancestry prediction results on the balanced 1kGP benchmark set of 100 samples (20 per super-population). The ancestry inference probability distribution of each sample's ancestry across a population group is shown for SNVstory. For ntRoot, the highest predicted GAI along with its score, and LAI-based super-population fraction estimates are reported. The super-populations are EAS: East Asian, SAS: South Asian, AMR: Admixed American, AFR: African, EUR: European.

| ACCESSION | ID | 1kGP assigned Label | AdM GAI | ntR GAI | SnvS GAI | ntR GAI score | SnvS GAI score | AdM (ancestry fractions) | | | | | ntR (LAI-based ancestry fractions) | | | | | SnvS (GAI probabilities) | | | | |
|---|---|---|---|---|---|---|---|---|---|---|---|---|---|---|---|---|---|---|---|---|---|---|
| | | | | | | | | EAS | SAS | AMR | AFR | EUR | EAS | SAS | AMR | AFR | EUR | EAS | SAS | AMR | AFR | EUR |
| **ERR3242231** | HG02282 | AFR | AFR | AFR | AFR | 0.4410 | 0.9930 | 0.00 | 0.00 | 0.00 | 0.92 | 0.08 | 0.00 | 0.00 | 0.06 | 0.91 | 0.03 | 0.00 | 0.00 | 0.00 | 0.99 | 0.00 |
| **ERR3989003** | HG02257 | AFR | AFR | AFR | AFR | 0.4419 | 0.9914 | 0.00 | 0.00 | 0.01 | 0.95 | 0.04 | 0.00 | 0.00 | 0.04 | 0.95 | 0.01 | 0.00 | 0.00 | 0.01 | 0.99 | 0.00 |
| **ERR3242485** | HG02974 | AFR | AFR | AFR | AFR | 0.4441 | 1.0000 | 0.00 | 0.00 | 0.00 | 1.00 | 0.00 | 0.00 | 0.00 | 0.00 | 1.00 | 0.00 | 0.00 | 0.00 | 0.00 | 1.00 | 0.00 |
| **ERR3989126** | HG03125 | AFR | AFR | AFR | AFR | 0.4419 | 0.9983 | 0.00 | 0.00 | 0.00 | 1.00 | 0.00 | 0.00 | 0.00 | 0.00 | 1.00 | 0.00 | 0.00 | 0.00 | 0.00 | 1.00 | 0.00 |
| **ERR3989162** | HG03371 | AFR | AFR | AFR | AFR | 0.4421 | 0.9983 | 0.00 | 0.00 | 0.00 | 1.00 | 0.00 | 0.00 | 0.00 | 0.00 | 1.00 | 0.00 | 0.00 | 0.00 | 0.00 | 1.00 | 0.00 |
| **ERR3989174** | HG03516 | AFR | AFR | AFR | AFR | 0.4431 | 1.0000 | 0.00 | 0.00 | 0.00 | 1.00 | 0.00 | 0.00 | 0.00 | 0.00 | 1.00 | 0.00 | 0.00 | 0.00 | 0.00 | 1.00 | 0.00 |
| **ERR3242376** | HG02614 | AFR | AFR | AFR | AFR | 0.4450 | 0.9978 | 0.00 | 0.00 | 0.00 | 1.00 | 0.00 | 0.00 | 0.00 | 0.01 | 0.99 | 0.00 | 0.00 | 0.00 | 0.00 | 1.00 | 0.00 |
| **ERR3242458** | HG02817 | AFR | AFR | AFR | AFR | 0.4444 | 0.9972 | 0.00 | 0.00 | 0.00 | 1.00 | 0.00 | 0.00 | 0.00 | 0.01 | 0.99 | 0.00 | 0.00 | 0.00 | 0.00 | 1.00 | 0.00 |
| **ERR3989031** | HG02587 | AFR | AFR | AFR | AFR | 0.4443 | 0.9973 | 0.00 | 0.00 | 0.00 | 1.00 | 0.00 | 0.00 | 0.00 | 0.01 | 1.00 | 0.00 | 0.00 | 0.00 | 0.00 | 1.00 | 0.00 |
| **ERR3989060** | HG02723 | AFR | AFR | AFR | AFR | 0.4416 | 0.9974 | 0.00 | 0.00 | 0.00 | 1.00 | 0.00 | 0.00 | 0.00 | 0.01 | 0.99 | 0.00 | 0.00 | 0.00 | 0.00 | 1.00 | 0.00 |
| **ERR3989080** | HG02818 | AFR | AFR | AFR | AFR | 0.4436 | 0.9973 | 0.00 | 0.00 | 0.00 | 1.00 | 0.00 | 0.00 | 0.00 | 0.01 | 0.99 | 0.00 | 0.00 | 0.00 | 0.00 | 1.00 | 0.00 |
| **ERR3989091** | HG02886 | AFR | AFR | AFR | AFR | 0.4419 | 0.9970 | 0.00 | 0.00 | 0.00 | 1.00 | 0.00 | 0.00 | 0.00 | 0.00 | 1.00 | 0.00 | 0.00 | 0.00 | 0.00 | 1.00 | 0.00 |
| **ERR3239692** | NA19030 | AFR | AFR | AFR | AFR | 0.4441 | 0.9958 | 0.00 | 0.00 | 0.00 | 1.00 | 0.00 | 0.00 | 0.00 | 0.01 | 0.99 | 0.00 | 0.00 | 0.00 | 0.00 | 1.00 | 0.00 |
| **ERR3239700** | NA19043 | AFR | AFR | AFR | AFR | 0.4421 | 0.9963 | 0.00 | 0.00 | 0.00 | 1.00 | 0.00 | 0.00 | 0.00 | 0.01 | 0.99 | 0.00 | 0.00 | 0.00 | 0.00 | 1.00 | 0.00 |
| **ERR3242560** | HG03069 | AFR | AFR | AFR | AFR | 0.4377 | 0.9984 | 0.00 | 0.00 | 0.00 | 1.00 | 0.00 | 0.00 | 0.00 | 0.00 | 1.00 | 0.00 | 0.00 | 0.00 | 0.00 | 1.00 | 0.00 |
| **ERR3242571** | HG03548 | AFR | AFR | AFR | AFR | 0.4390 | 0.9979 | 0.00 | 0.00 | 0.00 | 1.00 | 0.00 | 0.00 | 0.00 | 0.00 | 1.00 | 0.00 | 0.00 | 0.00 | 0.00 | 1.00 | 0.00 |
| **ERR3989118** | HG03065 | AFR | AFR | AFR | AFR | 0.4409 | 0.9980 | 0.00 | 0.00 | 0.00 | 1.00 | 0.00 | 0.00 | 0.00 | 0.01 | 1.00 | 0.00 | 0.00 | 0.00 | 0.00 | 1.00 | 0.00 |
| **ERR3989180** | HG03579 | AFR | AFR | AFR | AFR | 0.4412 | 0.9982 | 0.00 | 0.00 | 0.00 | 1.00 | 0.00 | 0.00 | 0.00 | 0.01 | 1.00 | 0.00 | 0.00 | 0.00 | 0.00 | 1.00 | 0.00 |
| **ERR3239351** | NA18522 | AFR | AFR | AFR | AFR | 0.4444 | 0.9985 | 0.00 | 0.00 | 0.00 | 1.00 | 0.00 | 0.00 | 0.00 | 0.01 | 0.99 | 0.00 | 0.00 | 0.00 | 0.00 | 1.00 | 0.00 |
| **ERR3239453** | NA19238 | AFR | AFR | AFR | AFR | 0.4420 | 0.9985 | 0.00 | 0.00 | 0.00 | 1.00 | 0.00 | 0.00 | 0.00 | 0.00 | 1.00 | 0.00 | 0.00 | 0.00 | 0.00 | 1.00 | 0.00 |
| **ERR3242786** | HG01122 | AMR | AMR | AMR | AMR | 0.4800 | 0.9976 | 0.00 | 0.00 | 0.55 | 0.16 | 0.29 | 0.04 | 0.05 | 0.48 | 0.11 | 0.32 | 0.00 | 0.00 | 1.00 | 0.00 | 0.00 |
| **ERR3242795** | HG01363 | AMR | AMR | AMR | AMR | 0.4642 | 0.9954 | 0.00 | 0.00 | 0.64 | 0.30 | 0.06 | 0.07 | 0.04 | 0.48 | 0.27 | 0.14 | 0.00 | 0.00 | 1.00 | 0.00 | 0.00 |
| **ERR3242796** | HG01369 | AMR | AMR | AMR | AMR | 0.4764 | 0.9951 | 0.00 | 0.00 | 0.79 | 0.21 | 0.00 | 0.10 | 0.03 | 0.56 | 0.21 | 0.10 | 0.00 | 0.00 | 1.00 | 0.00 | 0.00 |
| **ERR3242797** | HG01372 | AMR | AMR | AMR | AMR | 0.4878 | 0.9978 | 0.00 | 0.00 | 0.77 | 0.08 | 0.15 | 0.06 | 0.04 | 0.60 | 0.11 | 0.20 | 0.00 | 0.00 | 1.00 | 0.00 | 0.00 |
| **ERR3242805** | HG01468 | AMR | AMR | AMR | AMR | 0.4902 | 0.9975 | 0.00 | 0.00 | 0.56 | 0.06 | 0.38 | 0.03 | 0.07 | 0.46 | 0.09 | 0.35 | 0.00 | 0.00 | 1.00 | 0.00 | 0.00 |
| **ERR3988843** | HG01114 | AMR | AMR | AMR | AMR | 0.4892 | 0.9984 | 0.00 | 0.00 | 0.59 | 0.02 | 0.39 | 0.03 | 0.06 | 0.47 | 0.06 | 0.39 | 0.00 | 0.00 | 1.00 | 0.00 | 0.00 |
| **ERR3988876** | HG01361 | AMR | AMR | AMR | AMR | 0.4833 | 0.9975 | 0.00 | 0.00 | 0.71 | 0.10 | 0.19 | 0.07 | 0.06 | 0.50 | 0.11 | 0.26 | 0.00 | 0.00 | 1.00 | 0.00 | 0.00 |
| **ERR3988882** | HG01433 | AMR | AMR | AMR | AMR | 0.4784 | 0.9971 | 0.00 | 0.00 | 0.69 | 0.15 | 0.16 | 0.05 | 0.04 | 0.48 | 0.16 | 0.27 | 0.00 | 0.00 | 1.00 | 0.00 | 0.00 |
| **ERR3988889** | HG01496 | AMR | AMR | AMR | AMR | 0.4763 | 0.9953 | 0.00 | 0.00 | 0.80 | 0.20 | 0.00 | 0.10 | 0.04 | 0.56 | 0.19 | 0.11 | 0.00 | 0.00 | 1.00 | 0.00 | 0.00 |
| **ERR3989413** | NA19650 | AMR | AMR | AMR | AMR | 0.4825 | 0.9971 | 0.00 | 0.00 | 0.47 | 0.08 | 0.45 | 0.05 | 0.06 | 0.42 | 0.08 | 0.39 | 0.00 | 0.00 | 1.00 | 0.00 | 0.00 |
| **ERR3242814** | HG02102 | AMR | AMR | AMR | AMR | 0.5069 | 0.9970 | 0.05 | 0.00 | 0.95 | 0.00 | 0.00 | 0.24 | 0.01 | 0.70 | 0.03 | 0.02 | 0.00 | 0.00 | 1.00 | 0.00 | 0.00 |
| **ERR3988951** | HG01928 | AMR | AMR | AMR | AMR | 0.5134 | 0.9970 | 0.07 | 0.00 | 0.93 | 0.00 | 0.00 | 0.31 | 0.01 | 0.67 | 0.01 | 0.00 | 0.00 | 0.00 | 1.00 | 0.00 | 0.00 |
| **ERR3988952** | HG01934 | AMR | AMR | AMR | AMR | 0.4951 | 0.9965 | 0.04 | 0.00 | 0.94 | 0.02 | 0.00 | 0.22 | 0.02 | 0.64 | 0.07 | 0.05 | 0.00 | 0.00 | 1.00 | 0.00 | 0.00 |
| **ERR3988955** | HG01943 | AMR | AMR | AMR | AMR | 0.5068 | 0.9972 | 0.05 | 0.00 | 0.95 | 0.00 | 0.00 | 0.27 | 0.01 | 0.67 | 0.04 | 0.01 | 0.00 | 0.00 | 1.00 | 0.00 | 0.00 |
| **ERR3988958** | HG01952 | AMR | AMR | AMR | AMR | 0.5044 | 0.9963 | 0.04 | 0.00 | 0.96 | 0.00 | 0.00 | 0.26 | 0.01 | 0.67 | 0.06 | 0.01 | 0.00 | 0.00 | 1.00 | 0.00 | 0.00 |
| **ERR3988964** | HG01975 | AMR | AMR | AMR | AMR | 0.5031 | 0.9971 | 0.05 | 0.00 | 0.95 | 0.00 | 0.00 | 0.26 | 0.01 | 0.69 | 0.04 | 0.00 | 0.00 | 0.00 | 1.00 | 0.00 | 0.00 |
| **ERR3241754** | HG00731 | AMR | EUR | AMR | AMR | 0.4852 | 0.9979 | 0.00 | 0.00 | 0.27 | 0.10 | 0.63 | 0.02 | 0.05 | 0.43 | 0.09 | 0.41 | 0.00 | 0.00 | 1.00 | 0.00 | 0.00 |
| **ERR3241755** | HG00732 | AMR | AMR | AMR | AMR | 0.4815 | 0.9969 | 0.00 | 0.00 | 0.45 | 0.13 | 0.42 | 0.03 | 0.06 | 0.42 | 0.15 | 0.34 | 0.00 | 0.00 | 1.00 | 0.00 | 0.00 |
| **ERR3242761** | HG01395 | AMR | AMR | AMR | AMR | 0.4776 | 0.9970 | 0.00 | 0.00 | 0.41 | 0.18 | 0.41 | 0.02 | 0.06 | 0.46 | 0.15 | 0.31 | 0.00 | 0.00 | 1.00 | 0.00 | 0.00 |
| **ERR3242766** | HG01414 | AMR | AMR | AMR | AMR | 0.4674 | 0.9966 | 0.00 | 0.00 | 0.47 | 0.26 | 0.27 | 0.03 | 0.04 | 0.49 | 0.20 | 0.24 | 0.00 | 0.00 | 1.00 | 0.00 | 0.00 |
| **ERR3242132** | HG00864 | EAS | EAS | EAS | EAS | 0.5315 | 0.9949 | 0.97 | 0.02 | 0.00 | 0.02 | 0.00 | 0.97 | 0.01 | 0.01 | 0.01 | 0.00 | 0.99 | 0.00 | 0.00 | 0.00 | 0.00 |

| | | | | | | | | | | | | | | | | | | | | | | |
|---|---|---|---|---|---|---|---|---|---|---|---|---|---|---|---|---|---|---|---|---|---|---|
| **ERR3242142** | HG01046 | EAS | EAS | EAS | EAS | 0.5328 | 0.9952 | 0.99 | 0.00 | 0.00 | 0.01 | 0.00 | 0.98 | 0.00 | 0.01 | 0.00 | 0.00 | 1.00 | 0.00 | 0.00 | 0.00 | 0.00 |
| **ERR3242150** | HG01801 | EAS | EAS | EAS | EAS | 0.5303 | 0.9952 | 0.98 | 0.00 | 0.00 | 0.02 | 0.00 | 0.97 | 0.01 | 0.01 | 0.01 | 0.00 | 1.00 | 0.00 | 0.00 | 0.00 | 0.00 |
| **ERR3242160** | HG01812 | EAS | EAS | EAS | EAS | 0.5333 | 0.9967 | 0.99 | 0.00 | 0.00 | 0.01 | 0.00 | 0.98 | 0.01 | 0.01 | 0.01 | 0.00 | 1.00 | 0.00 | 0.00 | 0.00 | 0.00 |
| **ERR3241678** | HG00473 | EAS | EAS | EAS | EAS | 0.5303 | 0.9957 | 0.99 | 0.00 | 0.00 | 0.01 | 0.00 | 0.98 | 0.01 | 0.01 | 0.01 | 0.01 | 1.00 | 0.00 | 0.00 | 0.00 | 0.00 |
| **ERR3241684** | HG00513 | EAS | EAS | EAS | EAS | 0.5309 | 0.9964 | 0.99 | 0.00 | 0.00 | 0.01 | 0.00 | 0.98 | 0.01 | 0.00 | 0.01 | 0.00 | 1.00 | 0.00 | 0.00 | 0.00 | 0.00 |
| **ERR3241686** | HG00525 | EAS | EAS | EAS | EAS | 0.5353 | 0.9948 | 0.99 | 0.00 | 0.00 | 0.01 | 0.00 | 0.97 | 0.01 | 0.01 | 0.01 | 0.00 | 0.99 | 0.00 | 0.00 | 0.00 | 0.00 |
| **ERR3241750** | HG00705 | EAS | EAS | EAS | EAS | 0.5356 | 0.9973 | 1.00 | 0.00 | 0.00 | 0.00 | 0.00 | 0.98 | 0.00 | 0.00 | 0.01 | 0.00 | 1.00 | 0.00 | 0.00 | 0.00 | 0.00 |
| **ERR3242512** | HG00410 | EAS | EAS | EAS | EAS | 0.5308 | 0.9958 | 0.99 | 0.00 | 0.00 | 0.01 | 0.00 | 0.98 | 0.00 | 0.01 | 0.01 | 0.00 | 1.00 | 0.00 | 0.00 | 0.00 | 0.00 |
| **ERR3242514** | HG00599 | EAS | EAS | EAS | EAS | 0.5318 | 0.9953 | 0.99 | 0.00 | 0.00 | 0.01 | 0.00 | 0.98 | 0.00 | 0.01 | 0.01 | 0.00 | 1.00 | 0.00 | 0.00 | 0.00 | 0.00 |
| **ERR3242515** | HG00728 | EAS | EAS | EAS | EAS | 0.5334 | 0.9958 | 0.99 | 0.00 | 0.00 | 0.01 | 0.00 | 0.98 | 0.01 | 0.01 | 0.01 | 0.00 | 1.00 | 0.00 | 0.00 | 0.00 | 0.00 |
| **ERR3242519** | HG00631 | EAS | EAS | EAS | EAS | 0.5332 | 0.9957 | 1.00 | 0.00 | 0.00 | 0.00 | 0.00 | 0.98 | 0.01 | 0.01 | 0.01 | 0.00 | 1.00 | 0.00 | 0.00 | 0.00 | 0.00 |
| **ERR3242522** | HG00675 | EAS | EAS | EAS | EAS | 0.5332 | 0.9974 | 0.99 | 0.00 | 0.00 | 0.01 | 0.00 | 0.97 | 0.01 | 0.01 | 0.02 | 0.00 | 1.00 | 0.00 | 0.00 | 0.00 | 0.00 |
| **ERR3988762** | HG00408 | EAS | EAS | EAS | EAS | 0.5320 | 0.9961 | 1.00 | 0.00 | 0.00 | 0.00 | 0.00 | 0.98 | 0.01 | 0.00 | 0.01 | 0.00 | 1.00 | 0.00 | 0.00 | 0.00 | 0.00 |
| **ERR3988765** | HG00423 | EAS | EAS | EAS | EAS | 0.5323 | 0.9963 | 1.00 | 0.00 | 0.00 | 0.00 | 0.00 | 0.98 | 0.00 | 0.01 | 0.01 | 0.00 | 1.00 | 0.00 | 0.00 | 0.00 | 0.00 |
| **ERR3988768** | HG00438 | EAS | EAS | EAS | EAS | 0.5324 | 0.9956 | 1.00 | 0.00 | 0.00 | 0.00 | 0.00 | 0.97 | 0.01 | 0.01 | 0.01 | 0.00 | 1.00 | 0.00 | 0.00 | 0.00 | 0.00 |
| **ERR3988780** | HG00512 | EAS | EAS | EAS | EAS | 0.5329 | 0.9954 | 1.00 | 0.00 | 0.00 | 0.00 | 0.00 | 0.98 | 0.00 | 0.01 | 0.01 | 0.00 | 1.00 | 0.00 | 0.00 | 0.00 | 0.00 |
| **ERR3242087** | HG01862 | EAS | EAS | EAS | EAS | 0.5303 | 0.9950 | 0.95 | 0.04 | 0.00 | 0.01 | 0.00 | 0.97 | 0.01 | 0.01 | 0.01 | 0.00 | 1.00 | 0.00 | 0.00 | 0.00 | 0.00 |
| **ERR3242101** | HG02048 | EAS | EAS | EAS | EAS | 0.5309 | 0.9949 | 0.94 | 0.04 | 0.00 | 0.02 | 0.00 | 0.97 | 0.01 | 0.00 | 0.01 | 0.01 | 0.99 | 0.00 | 0.00 | 0.00 | 0.00 |
| **ERR3242342** | HG02040 | EAS | EAS | EAS | EAS | 0.5297 | 0.9948 | 0.95 | 0.04 | 0.00 | 0.02 | 0.00 | 0.97 | 0.01 | 0.01 | 0.01 | 0.00 | 0.99 | 0.00 | 0.00 | 0.00 | 0.00 |
| **ERR3239334** | NA12878 | EUR | EUR | EUR | EUR | 0.4951 | 0.9952 | 0.00 | 0.00 | 0.00 | 0.02 | 0.98 | 0.01 | 0.09 | 0.14 | 0.06 | 0.70 | 0.00 | 0.00 | 0.00 | 0.00 | 1.00 |
| **ERR3240160** | HG00171 | EUR | EUR | EUR | EUR | 0.4950 | 0.9957 | 0.09 | 0.00 | 0.01 | 0.00 | 0.89 | 0.05 | 0.09 | 0.14 | 0.05 | 0.67 | 0.00 | 0.00 | 0.00 | 0.00 | 1.00 |
| **ERR3240240** | HG00323 | EUR | EUR | EUR | EUR | 0.4950 | 0.9958 | 0.08 | 0.00 | 0.00 | 0.02 | 0.91 | 0.04 | 0.11 | 0.12 | 0.06 | 0.68 | 0.00 | 0.00 | 0.00 | 0.00 | 1.00 |
| **ERR3241788** | HG00280 | EUR | EUR | EUR | EUR | 0.4932 | 0.9956 | 0.07 | 0.00 | 0.00 | 0.01 | 0.91 | 0.04 | 0.10 | 0.12 | 0.05 | 0.69 | 0.00 | 0.00 | 0.00 | 0.00 | 1.00 |
| **ERR3241799** | HG00331 | EUR | EUR | EUR | EUR | 0.4934 | 0.9957 | 0.07 | 0.00 | 0.00 | 0.01 | 0.91 | 0.05 | 0.08 | 0.12 | 0.05 | 0.70 | 0.00 | 0.00 | 0.00 | 0.00 | 1.00 |
| **ERR3240114** | HG00096 | EUR | EUR | EUR | EUR | 0.4950 | 0.9966 | 0.00 | 0.00 | 0.00 | 0.02 | 0.98 | 0.01 | 0.10 | 0.13 | 0.05 | 0.71 | 0.00 | 0.00 | 0.00 | 0.00 | 1.00 |
| **ERR3240116** | HG00099 | EUR | EUR | EUR | EUR | 0.4947 | 0.9956 | 0.00 | 0.01 | 0.00 | 0.02 | 0.97 | 0.00 | 0.10 | 0.13 | 0.06 | 0.71 | 0.00 | 0.00 | 0.00 | 0.00 | 1.00 |
| **ERR3240121** | HG00105 | EUR | EUR | EUR | EUR | 0.4949 | 0.9963 | 0.00 | 0.00 | 0.00 | 0.03 | 0.96 | 0.02 | 0.10 | 0.15 | 0.05 | 0.68 | 0.00 | 0.00 | 0.00 | 0.00 | 1.00 |
| **ERR3240126** | HG00110 | EUR | EUR | EUR | EUR | 0.4981 | 0.9960 | 0.00 | 0.01 | 0.00 | 0.02 | 0.97 | 0.02 | 0.08 | 0.14 | 0.07 | 0.70 | 0.00 | 0.00 | 0.00 | 0.00 | 1.00 |
| **ERR3240135** | HG00127 | EUR | EUR | EUR | EUR | 0.4965 | 0.9955 | 0.00 | 0.01 | 0.00 | 0.02 | 0.97 | 0.01 | 0.12 | 0.12 | 0.05 | 0.71 | 0.00 | 0.00 | 0.00 | 0.00 | 1.00 |
| **ERR3240146** | HG00140 | EUR | EUR | EUR | EUR | 0.4961 | 0.9963 | 0.00 | 0.03 | 0.00 | 0.03 | 0.95 | 0.01 | 0.09 | 0.15 | 0.07 | 0.69 | 0.00 | 0.00 | 0.00 | 0.00 | 1.00 |
| **ERR3240155** | HG00151 | EUR | EUR | EUR | EUR | 0.4941 | 0.9960 | 0.00 | 0.01 | 0.00 | 0.03 | 0.96 | 0.01 | 0.10 | 0.13 | 0.07 | 0.69 | 0.00 | 0.00 | 0.00 | 0.00 | 1.00 |
| **ERR3240195** | HG00121 | EUR | EUR | EUR | EUR | 0.4976 | 0.9969 | 0.00 | 0.02 | 0.00 | 0.02 | 0.97 | 0.02 | 0.10 | 0.11 | 0.06 | 0.71 | 0.00 | 0.00 | 0.00 | 0.00 | 1.00 |
| **ERR3242186** | HG01790 | EUR | EUR | EUR | EUR | 0.4960 | 0.9960 | 0.00 | 0.01 | 0.00 | 0.02 | 0.97 | 0.02 | 0.11 | 0.13 | 0.06 | 0.69 | 0.00 | 0.00 | 0.00 | 0.00 | 1.00 |
| **ERR3241917** | HG01501 | EUR | EUR | EUR | EUR | 0.4875 | 0.9956 | 0.00 | 0.00 | 0.00 | 0.12 | 0.88 | 0.01 | 0.08 | 0.18 | 0.10 | 0.62 | 0.00 | 0.00 | 0.00 | 0.00 | 1.00 |
| **ERR3241951** | HG01615 | EUR | EUR | EUR | EUR | 0.4903 | 0.9959 | 0.00 | 0.00 | 0.00 | 0.07 | 0.93 | 0.01 | 0.09 | 0.15 | 0.08 | 0.67 | 0.00 | 0.00 | 0.00 | 0.00 | 1.00 |
| **ERR3242111** | HG01695 | EUR | EUR | EUR | EUR | 0.4832 | 0.9959 | 0.00 | 0.00 | 0.00 | 0.15 | 0.85 | 0.03 | 0.07 | 0.19 | 0.16 | 0.56 | 0.00 | 0.00 | 0.00 | 0.00 | 1.00 |
| **ERR3988892** | HG01505 | EUR | EUR | EUR | EUR | 0.4943 | 0.9952 | 0.00 | 0.00 | 0.00 | 0.07 | 0.93 | 0.02 | 0.09 | 0.16 | 0.09 | 0.65 | 0.00 | 0.00 | 0.00 | 0.00 | 1.00 |
| **ERR3239792** | NA20509 | EUR | EUR | EUR | EUR | 0.4920 | 0.9958 | 0.00 | 0.04 | 0.00 | 0.06 | 0.90 | 0.01 | 0.15 | 0.13 | 0.07 | 0.64 | 0.00 | 0.00 | 0.00 | 0.00 | 1.00 |
| **ERR3239873** | NA20805 | EUR | EUR | EUR | EUR | 0.4933 | 0.9957 | 0.00 | 0.02 | 0.00 | 0.05 | 0.93 | 0.01 | 0.09 | 0.16 | 0.06 | 0.68 | 0.00 | 0.00 | 0.00 | 0.00 | 1.00 |
| **ERR3989224** | HG03831 | SAS | SAS | SAS | SAS | 0.4910 | 0.9959 | 0.08 | 0.90 | 0.00 | 0.02 | 0.00 | 0.18 | 0.73 | 0.03 | 0.05 | 0.02 | 0.00 | 1.00 | 0.00 | 0.00 | 0.00 |
| **ERR3989225** | HG03834 | SAS | SAS | SAS | SAS | 0.4906 | 0.9953 | 0.07 | 0.91 | 0.00 | 0.02 | 0.00 | 0.18 | 0.73 | 0.03 | 0.04 | 0.02 | 0.00 | 1.00 | 0.00 | 0.00 | 0.00 |
| **ERR3240106** | NA20905 | SAS | SAS | SAS | SAS | 0.4901 | 0.9971 | 0.00 | 0.96 | 0.00 | 0.02 | 0.01 | 0.06 | 0.82 | 0.03 | 0.04 | 0.04 | 0.00 | 1.00 | 0.00 | 0.00 | 0.00 |
| **ERR3243148** | HG03868 | SAS | SAS | SAS | SAS | 0.4933 | 0.9963 | 0.00 | 0.98 | 0.00 | 0.02 | 0.00 | 0.07 | 0.84 | 0.01 | 0.05 | 0.04 | 0.00 | 1.00 | 0.00 | 0.00 | 0.00 |
| **ERR3989210** | HG03732 | SAS | SAS | SAS | SAS | 0.4929 | 0.9990 | 0.00 | 0.97 | 0.00 | 0.03 | 0.00 | 0.05 | 0.84 | 0.03 | 0.06 | 0.02 | 0.00 | 1.00 | 0.00 | 0.00 | 0.00 |
| **ERR3242635** | HG03668 | SAS | SAS | SAS | SAS | 0.4919 | 0.9964 | 0.00 | 0.95 | 0.00 | 0.03 | 0.02 | 0.08 | 0.76 | 0.04 | 0.05 | 0.06 | 0.00 | 1.00 | 0.00 | 0.00 | 0.00 |
| **ERR3243049** | HG02731 | SAS | SAS | SAS | SAS | 0.4909 | 0.9963 | 0.00 | 0.91 | 0.00 | 0.03 | 0.06 | 0.07 | 0.77 | 0.04 | 0.05 | 0.08 | 0.00 | 1.00 | 0.00 | 0.00 | 0.00 |
| **ERR3243053** | HG03022 | SAS | SAS | SAS | SAS | 0.4942 | 0.9963 | 0.00 | 0.98 | 0.00 | 0.02 | 0.00 | 0.06 | 0.82 | 0.03 | 0.04 | 0.05 | 0.00 | 1.00 | 0.00 | 0.00 | 0.00 |
| **ERR3989034** | HG02602 | SAS | SAS | SAS | SAS | 0.4916 | 0.9962 | 0.00 | 0.96 | 0.00 | 0.02 | 0.01 | 0.07 | 0.81 | 0.04 | 0.04 | 0.04 | 0.00 | 1.00 | 0.00 | 0.00 | 0.00 |
| **ERR3989056** | HG02698 | SAS | SAS | SAS | SAS | 0.4913 | 0.9960 | 0.00 | 0.91 | 0.00 | 0.02 | 0.07 | 0.06 | 0.79 | 0.05 | 0.04 | 0.06 | 0.00 | 1.00 | 0.00 | 0.00 | 0.00 |
| **ERR3989064** | HG02738 | SAS | SAS | SAS | SAS | 0.4911 | 0.9956 | 0.00 | 0.88 | 0.00 | 0.02 | 0.10 | 0.06 | 0.72 | 0.05 | 0.06 | 0.11 | 0.00 | 1.00 | 0.00 | 0.00 | 0.00 |
| **ERR3989139** | HG03239 | SAS | SAS | SAS | SAS | 0.4902 | 0.9946 | 0.00 | 0.81 | 0.00 | 0.03 | 0.16 | 0.05 | 0.73 | 0.06 | 0.05 | 0.11 | 0.00 | 0.99 | 0.00 | 0.00 | 0.00 |

| | | | | | | | | | | | | | | | | | | | | | | |
|---|---|---|---|---|---|---|---|---|---|---|---|---|---|---|---|---|---|---|---|---|---|---|
| **ERR3989173** | HG03492 | SAS | SAS | SAS | SAS | 0.4896 | 0.9964 | 0.00 | 0.94 | 0.00 | 0.03 | 0.03 | 0.07 | 0.78 | 0.05 | 0.05 | 0.06 | 0.00 | 1.00 | 0.00 | 0.00 | 0.00 |
| **ERR3989196** | HG03654 | SAS | SAS | SAS | SAS | 0.4897 | 0.9964 | 0.00 | 0.98 | 0.00 | 0.02 | 0.00 | 0.06 | 0.78 | 0.05 | 0.05 | 0.06 | 0.00 | 1.00 | 0.00 | 0.00 | 0.00 |
| **ERR3989197** | HG03669 | SAS | SAS | SAS | SAS | 0.4914 | 0.9962 | 0.00 | 0.97 | 0.00 | 0.03 | 0.00 | 0.09 | 0.77 | 0.04 | 0.05 | 0.05 | 0.00 | 1.00 | 0.00 | 0.00 | 0.00 |
| **ERR3242695** | HG03898 | SAS | SAS | SAS | SAS | 0.4939 | 0.9960 | 0.00 | 0.98 | 0.00 | 0.02 | 0.00 | 0.07 | 0.84 | 0.02 | 0.04 | 0.03 | 0.00 | 1.00 | 0.00 | 0.00 | 0.00 |
| **ERR3989200** | HG03688 | SAS | SAS | SAS | SAS | 0.4917 | 0.9973 | 0.00 | 0.98 | 0.00 | 0.02 | 0.00 | 0.07 | 0.84 | 0.02 | 0.05 | 0.03 | 0.00 | 1.00 | 0.00 | 0.00 | 0.00 |
| **ERR3989258** | HG04199 | SAS | SAS | SAS | SAS | 0.4927 | 0.9971 | 0.00 | 0.98 | 0.00 | 0.02 | 0.00 | 0.10 | 0.81 | 0.03 | 0.05 | 0.02 | 0.00 | 1.00 | 0.00 | 0.00 | 0.00 |
| **ERR3989259** | HG04204 | SAS | SAS | SAS | SAS | 0.4917 | 0.9973 | 0.00 | 0.98 | 0.00 | 0.02 | 0.00 | 0.07 | 0.83 | 0.03 | 0.04 | 0.03 | 0.00 | 1.00 | 0.00 | 0.00 | 0.00 |
| **ERR3989262** | HG04228 | SAS | SAS | SAS | SAS | 0.4929 | 0.9980 | 0.00 | 0.98 | 0.00 | 0.02 | 0.00 | 0.11 | 0.81 | 0.02 | 0.03 | 0.03 | 0.00 | 1.00 | 0.00 | 0.00 | 0.00 |

**We note that ntRoot (ntR) predicts both a GAI and LAI-based ancestry fraction, and the former is not derived from the latter. As such, ntRoot LAI-based fraction estimates are not expected to match the highest GAI score.**

**Table S13.** Resource usage breakdown by steps for ADMIXTURE on the balanced set of 100, 1kGP samples.

| Step | Wall-clock run time (h) | Peak RAM (GB) |
|---|---|---|
| **1-Making VCFs** | 4.42 | 87.35 |
| **2-Indexing VCFs** | 0.001715 | 0.030000 |
| **3-Merge Query VCFs** | 0.67 | 2.93 |
| **4-Reheader the VCF** | 0.13 | 0.06 |
| **5-Merge with ref VCF** | 34.44 | 0.14 |
| **6-Filter VCF** | 24.49 | 0.08 |
| **7-Index filtered VCF** | 5.83 | 0.02 |
| **8-plink2** | 2.40 | 14.06 |
| **9-ADMIXTURE** | 201.32 | 449.15 |

**Benchmarks are based on the full sample set, with the exception of creating the VCFs and indexing the individual sample VCFs, which are averaged over the 100 samples.**

### Supplementary Figures

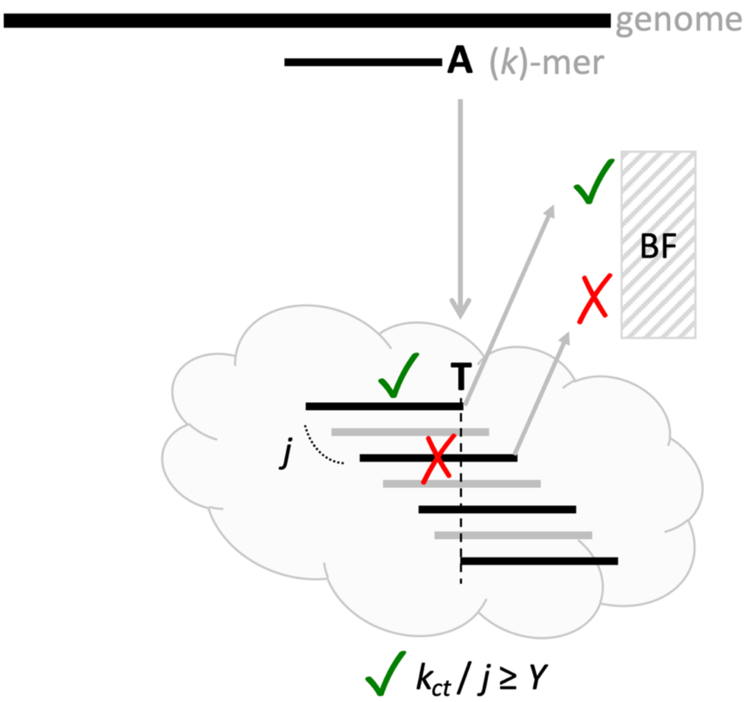

**Figure S1.** Overview of the SNV detection algorithm in ntRoot. Using k-mers (e.g., black line ending with A), sliding along the GRCh38 genome (top, black line), ntEdit v2 interrogates each base in turn by checking for presence (green checks) or absence (red crosses) in a k-mer Bloom filter (BF) built from ntHits (v1.0.2; https://github.com/bcgsc/nthits) for WGS or btllib[17] for WGA datasets, respectively. For each alternate base inspected (e.g., black line ending in T), k-mers overlapping the base under scrutiny (any black lines intersecting with the dashed lined within the cloud) are queried, while jumping over to the next k-mer by j bases. Once $k/j$ k-mers have been queried, a SNV is validated by assessing the proportion of successful Bloom filter k-mer hits ($k_{ct}/j$), and confirming it meets or exceeds the user-defined $-Y$ parameter (set at 0.55 or 55% by default).

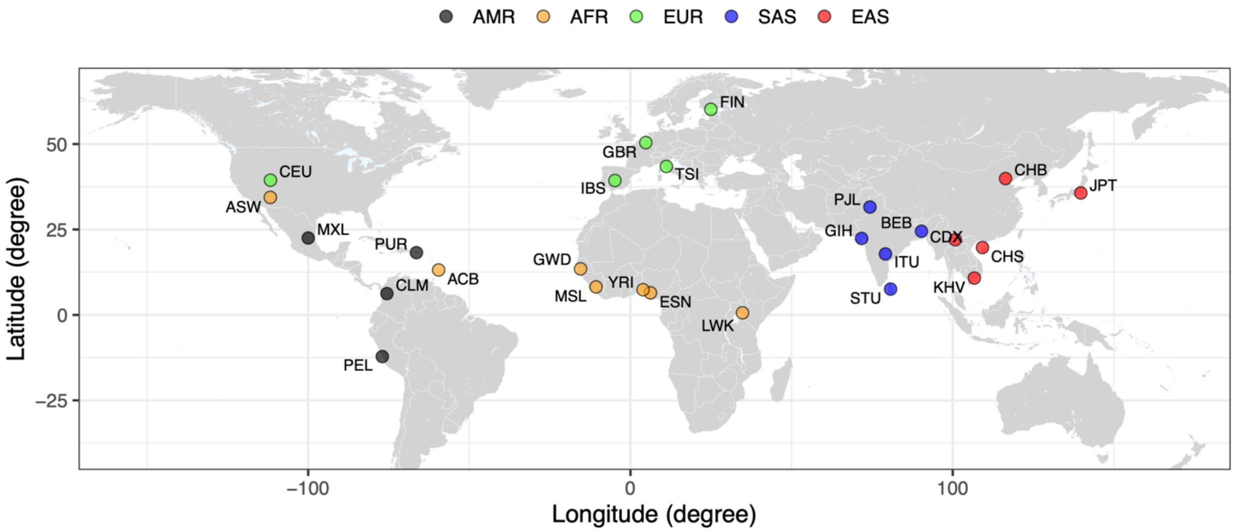

**Figure S2.** Geolocation of the 26 human populations, as defined by the 1000 Genomes Project. Human populations (MXL: Mexican-American, PUR: Puerto Rican, PEL: Peruvian, CLM: Colombian, ACB: African-Caribbean, ASW: African-American South West, YRI: Yoruba, GWD: Gambian, ESN: Esan, MSL: Mende, LWK: Luhya, CEU: CEPH (Centre d'Étude du Polymorphisme Humain – Utah), GBR: British, IBS: Spanish, TSI: Tuscan, FIN: Finnish, STU: Sri Lankan, BEB: Bengali, ITU: Indian, PJL: Punjabi, GIH: Gujarati, CDX: Dai Chinese, KHV: Kinh Vietnamese, CHS: Southern Han Chinese, CHB: Han Chinese, JPT: Japanese), coloured by the super-population they belong to (AMR: Admixed American, black; AFR: African, orange; EUR: European, green; SAS: South Asian, blue; EAS: East Asian, red) are shown in the global context.

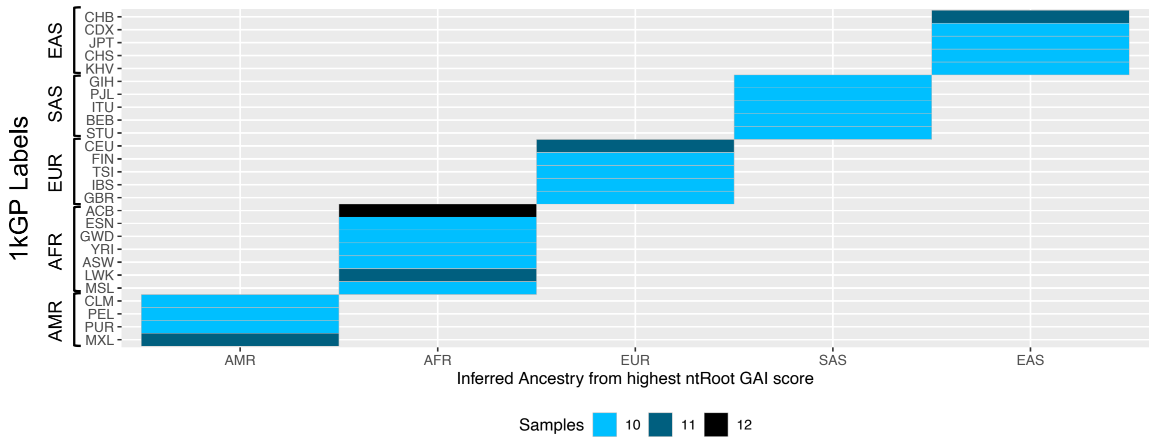

**Figure S3.** ntRoot super-population level GAI compared to 1kGP labels for the 1kGP WGS validation set. ntRoot super-population GAI predictions (x-axis) are compared to the 1kGP population ancestry labels (y-axis). Super-populations include EAS: East Asian; SAS: South Asian; AMR: Admixed American; AFR: African; EUR: European and are further arranged by human populations MXL: Mexican-American, PUR: Puerto Rican, PEL: Peruvian, CLM: Colombian, ACB: African-Caribbean, ASW: African-American South West, YRI: Yoruba, GWD: Gambian, ESN: Esan, MSL: Mende, LWK: Luhya, CEU: CEPH, GBR: British, IBS: Spanish, TSI: Tuscan, FIN: Finnish, STU: Sri Lankan, BEB: Bengali, ITU: Indian, PJL: Punjabi, GIH: Gujarati, CDX: Dai Chinese, KHV: Kinh Vietnamese, CHS: Southern Han Chinese, CHB: Han Chinese, JPT: Japanese. The light-to-dark blue colour indicates the number of individuals processed for each human population.

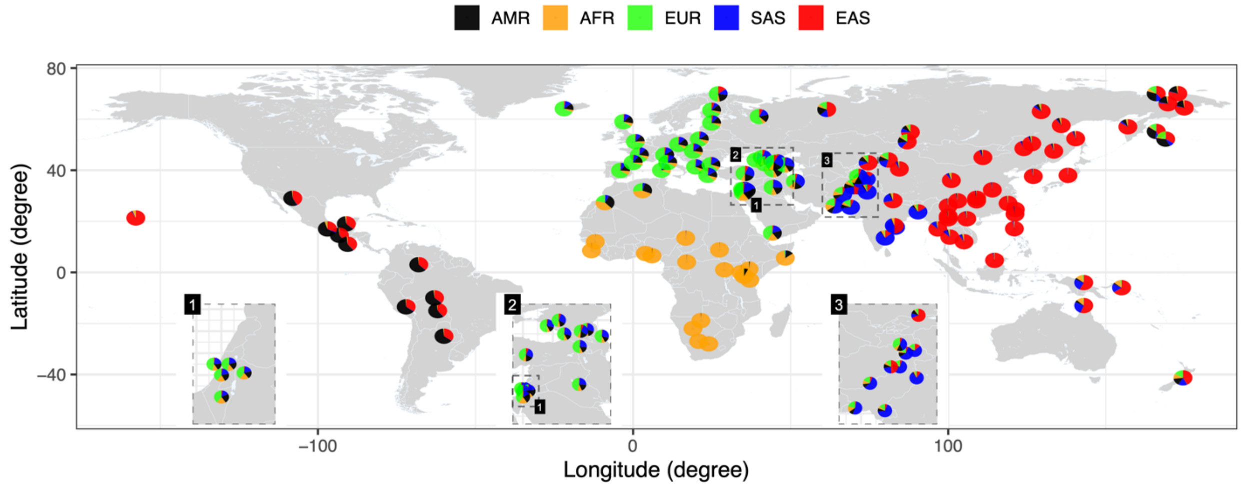

**Figure S4.** ntRoot ancestry composition estimates derived from LAI on SGDP WGS discovery data set from 129 distinct populations shown in the global context, without population labels and with inset regions zoomed-in. LAI-based ntRoot ancestry fraction estimates on the SGDP WGS discovery set for each population, using a representative individual randomly selected from Table S5 data, above. Ancestry fractions are coloured by super-population (AMR: Admixed American, black; AFR: African, orange; EUR: European, green; SAS: South Asian, blue; EAS: East Asian, red).

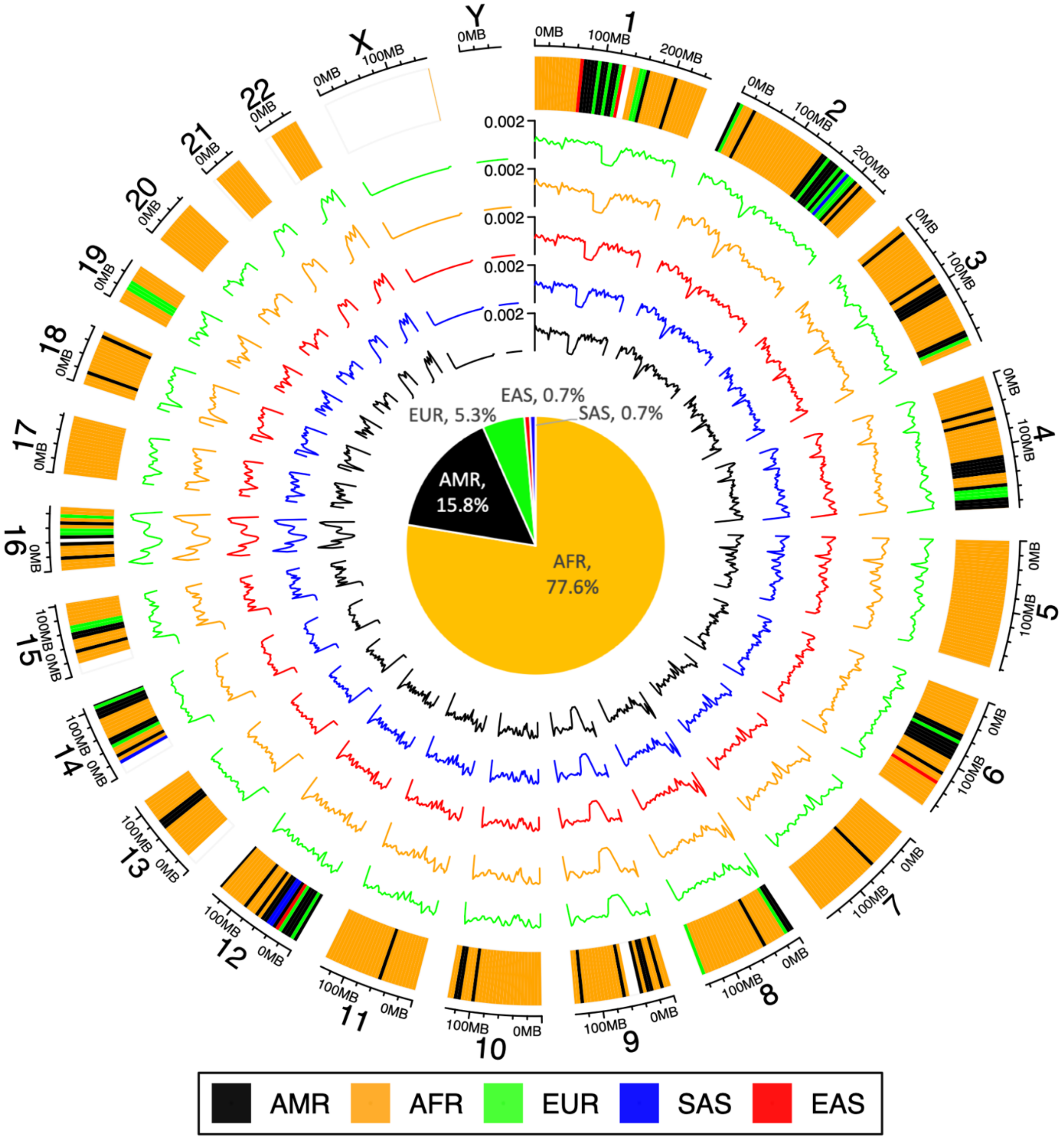

**Figure S5.** ntRoot LAI predictions on HG01243 Illumina whole genome sequencing data from 1kGP for a Puerto Rican individual of mixed, primarily African ancestry, labeling the likely human super-population matching each 5Mbp tile alongside the genome. The inner tracks show the SNV density (overlapping 5Mbp tiles) for EUR (European, green), AFR (African, orange), EAS (East Asian, red), SAS (South Asian, blue) and AMR (American, black). The inner pie chart shows the GAI fraction for each, the sum of each tile size assigned to a super-population over the sum of all tile sizes.

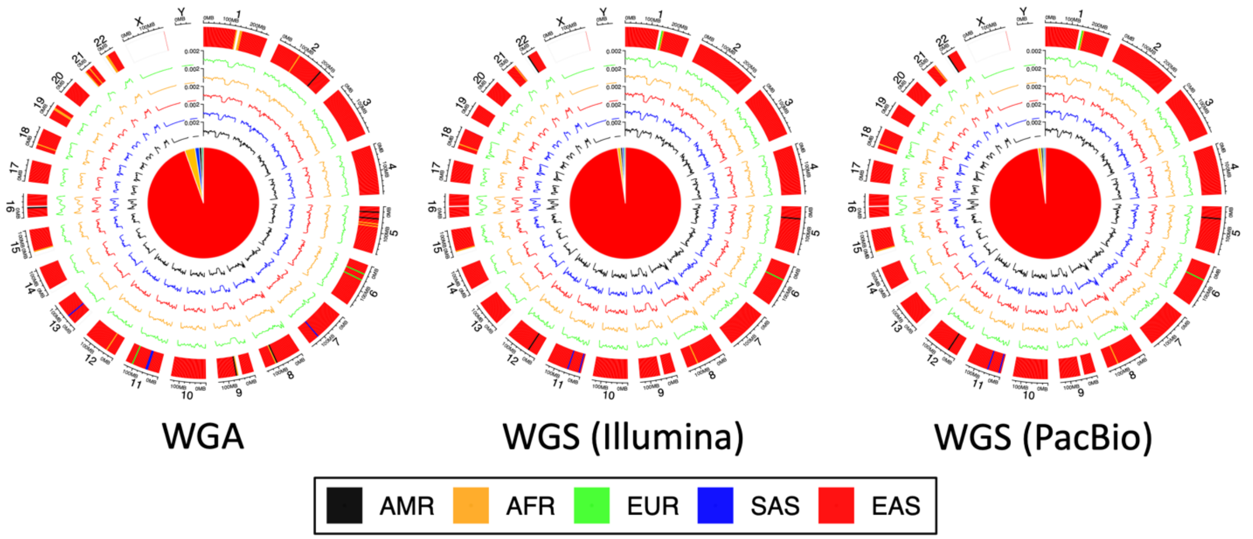

**Figure S6.** ntRoot LAI predictions on KOREF from **a**, Chromosome-scale pseudohaploid whole genome assembly reference, **b**, Illumina whole genome sequencing data and **c**, PacBio Sequel II sequencing data. LAI is shown on the outer track, labeling the likely human super-population matching each 5Mbp tile alongside the genome. The inner tracks show the SNV density (overlapping 5Mbp tiles) for EUR (European, green), AFR (African, orange), EAS (East Asian, red), SAS (South Asian, blue) and AMR (American, black). The inner pie chart shows the GAI fraction for each, the sum of each tile size assigned to a super-population over the sum of all tile sizes.

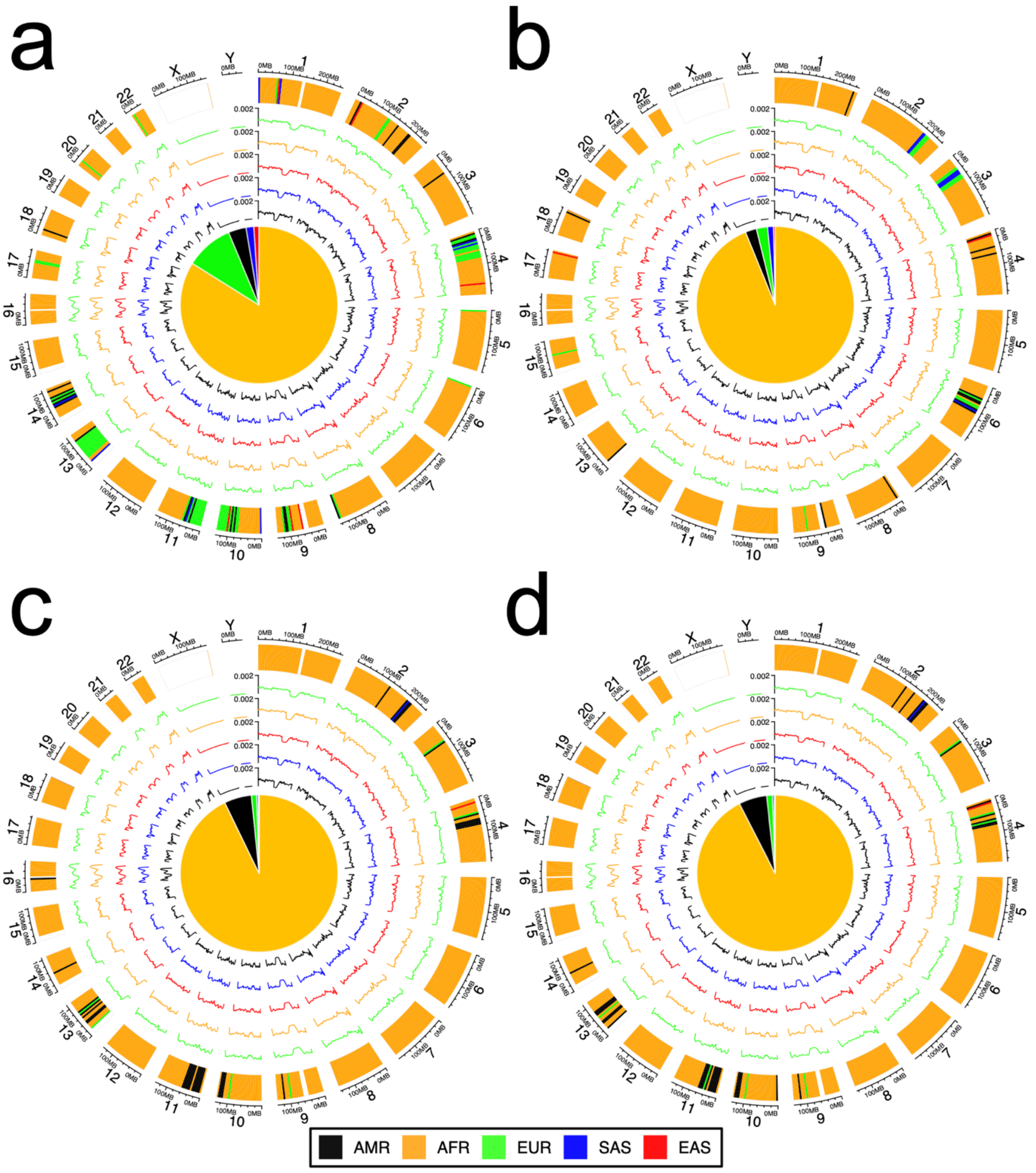

**Figure S7.** ntRoot LAI predictions on HG02055 directly on **a,** Maternal and **b,** paternal chromosome-complete haploid reference genomes and **c,** Flye and **d,** Shasta pseudohaploid genome assembly drafts. LAI is shown on the outer track, labeling the likely human super-population matching each 5Mbp tile alongside each genome. The inner tracks show the SNV density (overlapping 5Mbp tiles) for EUR (European, green), AFR (African, orange), EAS (East Asian, red), SAS (South Asian, blue) and AMR (American, black). The inner pie chart shows the GAI fraction for each, the sum of each tile size assigned to a super-population over the sum of all tile sizes.

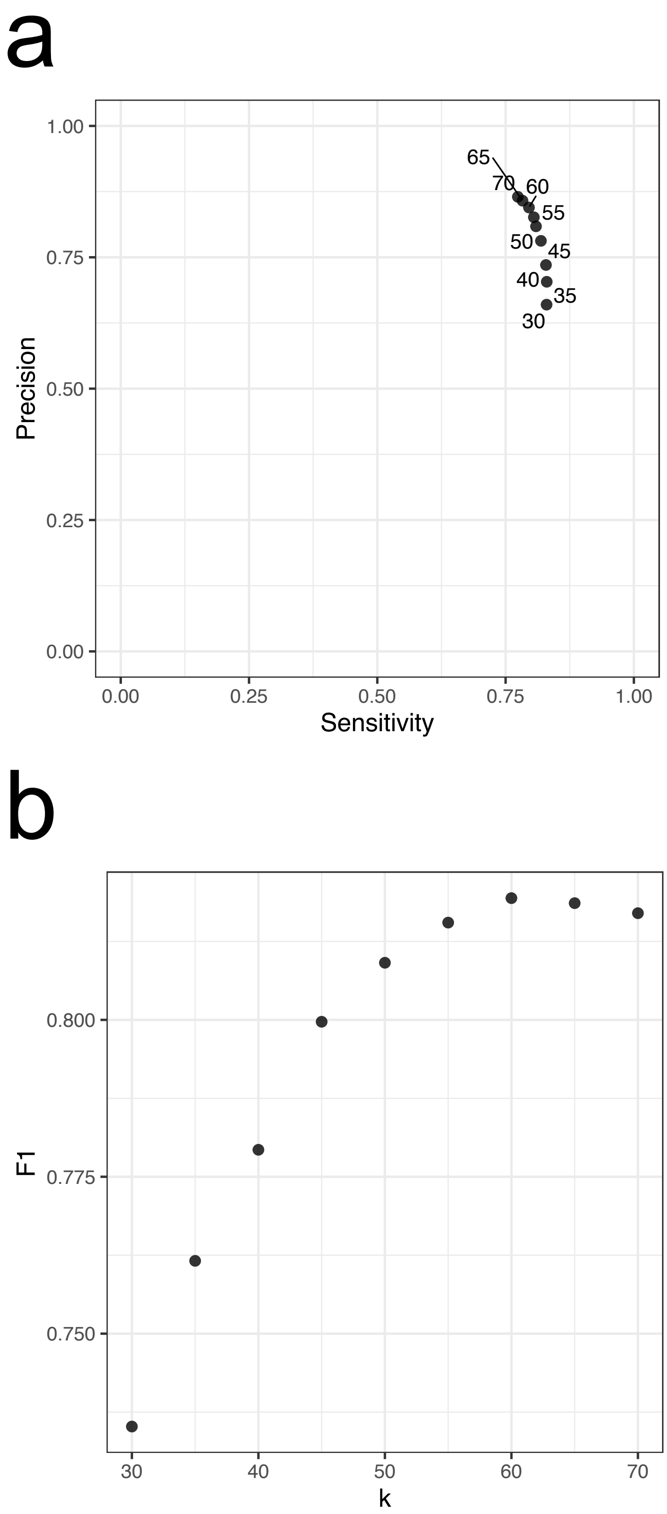

**Figure S8.** Rationale for selecting the parameter k in ntRoot. ntRoot performance a) Sensitivity vs. Precision and b) F1 score in identifying ancestry-discriminant SNVs against a benchmark set of known HG002 variants[12] as a function of k.

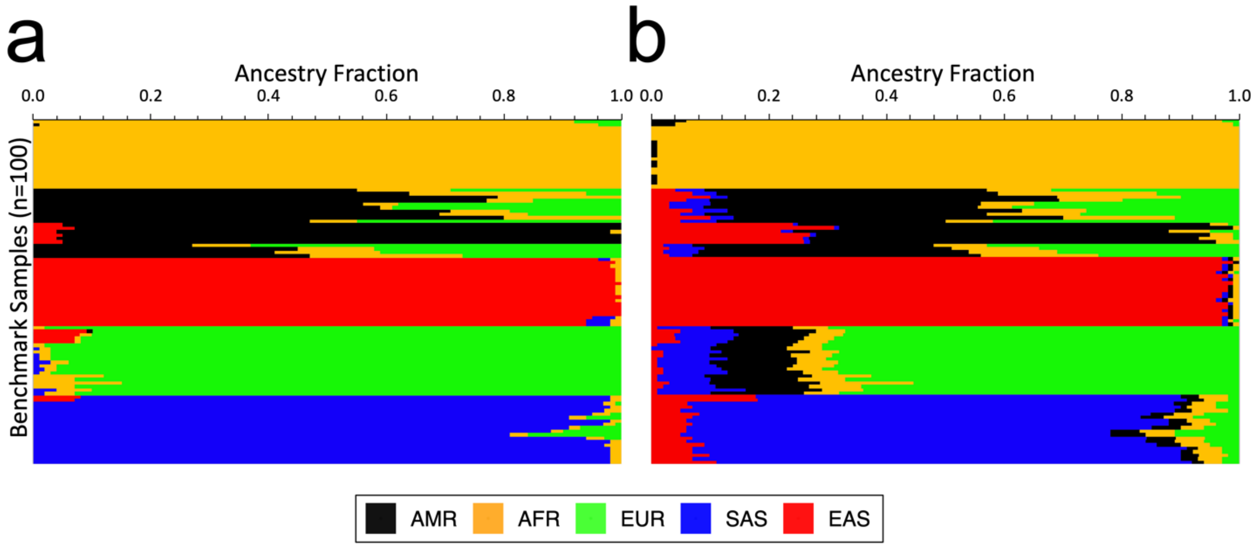

**Figure S9. a,** ADMIXTURE and **b,** ntRoot ancestry fraction predictions (y-axis) on the balanced set of 100, 1kGP samples (x-axis, matching the order of Table S10 from top to bottom). The fractions are represented as horizontal stacked bars for each super-population label: EUR (European, green), AFR (African, orange), EAS (East Asian, red), SAS (South Asian, blue), and AMR (American, black).

a

b

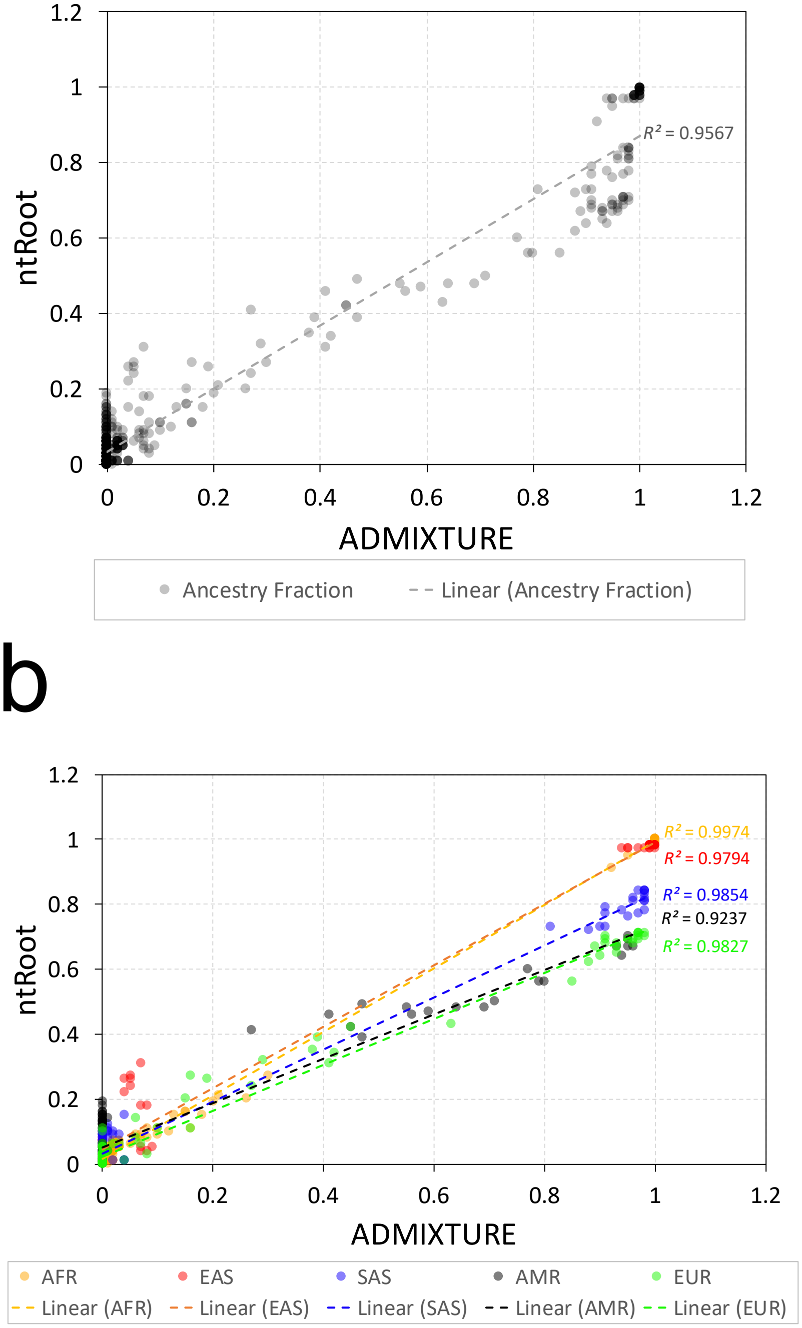

**Figure S10.** Correlation between ntRoot and ADMIXTURE ancestry fractions on the balanced set of 100, 1kGP samples. **a,** Scatter plot showing ancestry fractions estimated by ntRoot (y-axis) against ADMIXTURE (x-axis) across all ancestries and **b,** for each of the five separate ancestry group: AFR (African, orange), EAS (East Asian, red), SAS (South Asian, blue), AMR (American, black), and EUR (European, green). Linear regression lines are shown for each group, with corresponding $R^2$ values displayed.

### References

1. 1000 Genomes Project Consortium. *Nature* **526**, 68 (2015).

2. Byrska-Bishop, M. *et al. Cell* **185**, 3426-3440.e19 (2022).

3. Bollas, A. E. *et al. BMC Bioinformatics* **25**, 76 (2024).

4. Li, H. Preprint at https://doi.org/10.48550/ARXIV.1303.3997 (2013).

5. McKenna, A. *et al. Genome Res.* **20**, 1297–1303 (2010).

6. Van der Auwera, G. A. & O'Connor, B. D. (O'Reilly Media, Inc., 2020).

7. Danecek, P. *et al. GigaScience* **10**, giab008 (2021).

8. Alexander, D. H., Novembre, J. & Lange, K. *Genome Res.* **19**, 1655–1664 (2009).

9. Chang, C. C. *et al. Gigascience* **4**, s13742-015-0047–8 (2015).

10. Mallick, S. *et al. Nature* **538**, 201–206 (2016).

11. Wong, J. *et al. Nat. Commun.* **14**, 2906 (2023).

12. Zook, J. M. *et al. Nat. Biotechnol.* **37**, 561–566 (2019).

13. Levy, S. *et al. PLoS Biol.* **5**, e254 (2007).

14. Yang, C. *et al. Cell Res.* **33**, 745–761 (2023).

15. Kim, H.-S. *et al. GigaScience* **11**, giac022 (2022).

16. Liao, W.-W. *et al. Nature* **617**, 312–324 (2023).

17. Nikolić, V. *et al. J. Open Source Softw.* **7**, 4720 (2022).
